# Evaluating the Effects of Aircraft Noise on Hearing and Physiological Indicators: A Study of Military Personnel Using iTRAQ Proteomics and Cognitive Assessments

**DOI:** 10.1101/2024.09.25.615093

**Authors:** Manish Shukla, Jai Chand Patel, Devasharma Nayak, Meenakshi Shukla, Shutanu Chakravarti, Neeru Kapoor

## Abstract

**Background:** Noise pollution poses a significant public health risk, with prolonged exposure to high levels of noise linked to various adverse outcomes such as annoyance, sleep disturbances, cognitive impairment, hypertension, and cardiovascular diseases. Noise-induced hearing loss (NIHL) is a prominent concern in noisy occupational settings.

**Method:** This study investigated NIHL among 621 male Air Force soldiers aged 18-45, exposed to intense aircraft noise. Auditory assessments included pure-tone audiometry (125 Hz to 8 kHz) to categorize hearing into normal, mild, moderate, and severe impairment. Distortion product otoacoustic emissions (DPOAE) and auditory brainstem response (ABR) were used to evaluate cochlear and auditory nerve function. Heart rate variability (HRV) provided insights into autonomic responses. Cognitive functions were assessed through computerized tests, and blood plasma was analyzed for cardiac biomarkers, oxidative stress indicators, inflammatory markers, and neurotransmitters. Proteomic analysis used iTRAQ labeling, MudPIT, and MALDI-TOF/TOF mass spectrometry for protein quantification and identification, with validation through ELISA.

**Results:** The audiometric tests revealed varying degrees of hearing impairment, with significant threshold differences at 2000, 3000, 4000, and 6000 Hz, especially pronounced at 6000 Hz. The right ear showed greater impairment, and a characteristic high-frequency notch was observed, consistent with noise exposure. Proteomic analysis indicated that NIHL is associated with oxidative stress and systemic inflammation, with differential protein expression related to hearing, coagulation, and inflammation.

**Conclusion:** This study highlights the severe impact of aircraft noise on hearing and systemic health, demonstrating correlations between hearing impairment and biochemical markers. It emphasizes the role of oxidative stress and inflammation in NIHL development and underscores the need for effective noise management and protective measures in noisy work environments.

## Background

Nowadays, noise pollution has become a public health problem [1–4]. Overexposure to noisy environment can cause various kinds of dysfunction, such as annoyance [5], sleep disturbance [6], cognition impairment [7, 8], hypertension and cardiovascular diseases [9, 10], besides hearing impairment [11] which is the primary noise-related dysfunction. Noise (aircraft, road, transportation and railway etc) induces an activation of the cerebral cortex, the the hypotalmus-pitutary-adernal axis, and sympathetic nervous system characterized. Noise-induced hearing loss (NIHL) acquired in leisure or occupational settings is a common cause of hearing impairment in industrialized countries, with a prevalence second only to age-related hearing loss (ARHL) [12, 13]. As an environmental stimulus, noise can induce behavioral changes, autonomic function, and stress responses at receptors composed of multiple neurotransmitter release [14]. The stress response is believed to be one of the main causes of psychiatric disorders such as anxiety [15].

Noise intensity and duration of exposure determine the level of noise damage to an organism. High-intensity noise exposure damages inner hair cells (IHCs), primarily through two pathways: direct mechanical damage, in which noise can destroy the static cilia of hair cells, resulting in hair cell loss and damage to supporting cells and spiral ganglia [16, 17].

Aircraft noise exposure leads to an over activation of the sympathetic system, resulting in elevated levels of noradrenalin (NA), adrenalin (A), angiotensin II (Ang II), and subsequently cortisol [18]. Oxidative stress is defined as the imbalance in the redox characteristics of some cellular environment which can be the result of either biochemical processes leading to the production of reactive species, exposure to damaging agents (i.e., environmental pollutants and radiations), or limited capabilities of endogenous antioxidant systems [19–22]. Reactive oxygen and nitrogen species (ROS/RNS) produced under oxidative stress are known to damage all cellular biomolecules (lipids, sugars, proteins, and polynucleotide) [23, 24]. Thus, several defense systems have been involved within the cells to prevent uncontrolled ROS increase. During the past decade, research has revealed a widespread involvement of oxidative stress in several disease processes, including cancer, cardiovascular disease (CVD), atherosclerosis, diabetes, arthritis, neurodegenerative disorders, and pulmonary, renal, and hepatic diseases[25–27].

Noise-induced cognitive impairment has become a significant public health concern, particularly in urban environments and occupational settings where noise exposure is prevalent. Chronic exposure to high levels of environmental noise, such as traffic, industrial sounds, and even prolonged exposure to loud music, has been linked to various adverse health outcomes, including cognitive decline [28, 29]. Understanding the underlying mechanisms of noise-induced cognitive impairment is crucial for developing effective interventions and public health policies. Cognitive functions encompass various mental processes, including memory, attention, and executive function [30]. Shukla et al. reported that exposure to chronic noise can adversely affect these cognitive domains [7, 8, 28]. One studies have shown that individuals living in noisy urban environments exhibit deficits in cognitive performance compared to those in quieter settings. This cognitive decline is particularly concerning in vulnerable populations such as children, the elderly, and those with preexisting health conditions [28, 31]. Chronic noise exposure is known to induce stress responses that can lead to changes in vascular health. Prolonged exposure to high noise levels can cause endothelial dysfunction, a condition where the lining of blood vessels becomes damaged, impairing their ability to regulate blood flow and blood pressure effectively T [32]. Endothelial dysfunction is a precursor to atherosclerosis, a condition characterized by the buildup of plaques in the arterial walls [33, 34]. These plaques can obstruct blood flow, leading to reduced oxygen and nutrient delivery to the brain, which is essential for maintaining cognitive function. Additionally, impaired cerebral blood flow due to vascular changes can contribute to cognitive decline by affecting brain regions involved in memory and learning [35]. Studies have found that individuals exposed to chronic noise have elevated levels of blood pressure and markers of vascular inflammation [34], both of which are risk factors for cognitive impairment. The cumulative effect of these vascular changes can exacerbate age-related cognitive decline and increase the risk of developing neurodegenerative diseases [36].

To investigate the genesis and mechanism of intense noise-induced hearing loss, a study was conducted involving armed forces personnel exposed to high-intensity noise. The research aimed to correlate auditory impairment with physiological and psychological markers through a comprehensive analysis involving several methodologies. Participants’ hearing was evaluated using pure-tone audiometry, Distortion Product Otoacoustic Emissions (DPOAE), and Auditory Brainstem Response (ABR). These assessments were correlated with Heart Rate Variability (HRV) to gauge autonomic function. Cognitive and Behavioral Assessments: Cognitive and behavioral functions were assessed using the Cambridge Neuropsychological Test Automated Battery (CANTAB), focusing on Spatial Working Memory (SWM), Pattern Alternate Learning (PAL), and Reaction Time (SRT). Plasma and/or serum samples were analyzed to quantify cardiac, stress, and inflammatory markers. This involved the use of advanced mass spectrometry techniques, specifically 4-plex iTRAQ-LC-MS/MS, to identify and quantify plasma proteins. A total of 176 proteins were identified and quantified, with the analysis revealing differential expression patterns across hearing impairment groups categorized as mild, moderate, and severe. The study elucidates the complex interplay between noise exposure, hearing impairment, and associated physiological and psychological responses. The identification of specific plasma proteins related to hearing loss severity provides insights into potential biomarkers for assessing and mitigating noise-induced auditory damage.

## Clinical Relevance

Personnel exposed to high-frequency noises, like alarms and buzzers, may struggle to hear these crucial sounds effectively, especially in noisy environments. This can also make it hard to distinguish specific speech consonants. Noise-Induced Hearing Loss (NIHL) often develops gradually, so individuals might not immediately notice the extent of their hearing loss. The real impact usually becomes clear when these auditory challenges interfere with daily communication. Difficulty with high-frequency sounds can impair the ability to follow training and perform tasks, affecting overall performance and safety. Thus, early detection and intervention are essential to address the risks of NIHL.

## Material and Methods

### Subject selection and ethical considerations in study design

A study of the prevalence and severity of NIHL in ∼621 male Air force soldiers aged between 18-45 years, mean age 28.44±7.49 years and having exposure to intense occupational noise were selected to participate in the study. All of them were explained about the aims, objectives, methodology, and the anticipated benefits of the protocol, which was duly approved by the Institute’s Ethics Committee. The study was approved by the Institutional Human Ethical Committee (IHEC) and adhered to the ethical standards set by the Indian Council of Medical Research (ICMR) [approval no. 01/IEC/DIPAS/14-20]. The participants were explained with the details of the study design and written informed consent was obtained. All the volunteers were asked to fill in a questionnaire comprising information pertaining to health history, occupational history, education level, diet, personal habits etc. Personnel with some history of health problem or with brain surgery, stroke, post-trauma and with night shift before blood draw were excluded from the study. None of the volunteers reported any medicine intake, presence of hereditary or chronic disease. During study, all participants had taken the same type of diet as per the army ration scale and were not undergoing any supplementation with any vitamin. Throughout the study period volunteers had followed daily one-hour physical training protocol as per army training schedule (slow running, body flexibility exercise, and pull ups) in the morning. All the volunteers shared similar lifestyles, age and physical characteristics.

### Audiometer assessment

The 621 subjects underwent audiometric screening in a sound-attenuated room to ensure accurate measurements. The assessment involved recording noise levels at various workstations using a Brüel & Kjær Type 2260 Type 1 Modular Precision Sound Analyzer. The measurements included A-weighted Sound Pressure Level (SPL, dBA), A-weighted Equivalent Sound Level (LAeq), and noise dose. Audiometric evaluations were conducted using a Grason-Stadler Audiometer Model GSI 61, which was calibrated to a resolution of 1 dB. Hearing thresholds were determined for each ear at frequencies of 250 Hz, 500 Hz, 1000 Hz, 2000 Hz, 4000 Hz, 6000 Hz, and 8000 Hz. A threshold greater than 25 dB at any frequency was classified as hearing loss. The severity of hearing loss was graded as follows: Normal: Thresholds below 25 dB, Mild: Thresholds ranging from 26 to 40 dB, Moderate: Thresholds ranging from 41 to 60 dB, Severe: Thresholds of 60 dB or higher(Kapoor, Mani, and Shukla 2019).

### Auditory brain-stem response (ABR) assessment

To assess auditory function, we used Auditory Brainstem Response (ABR) testing. This method measures the brainstem’s response to sound, tracking how the auditory nerve and central pathways react to brief acoustic clicks or tones. It’s a straightforward and reliable approach for evaluating hearing thresholds. The Neurosoft Neurosystem (Russia) was used to generate and record the acoustic stimuli. We placed two-channel ABR electrodes at the mastoid regions of both ears (active electrodes), the scalp vertex (reference electrode), and the forehead (ground electrode) (Fig. 2a).

**Fig. 1.**
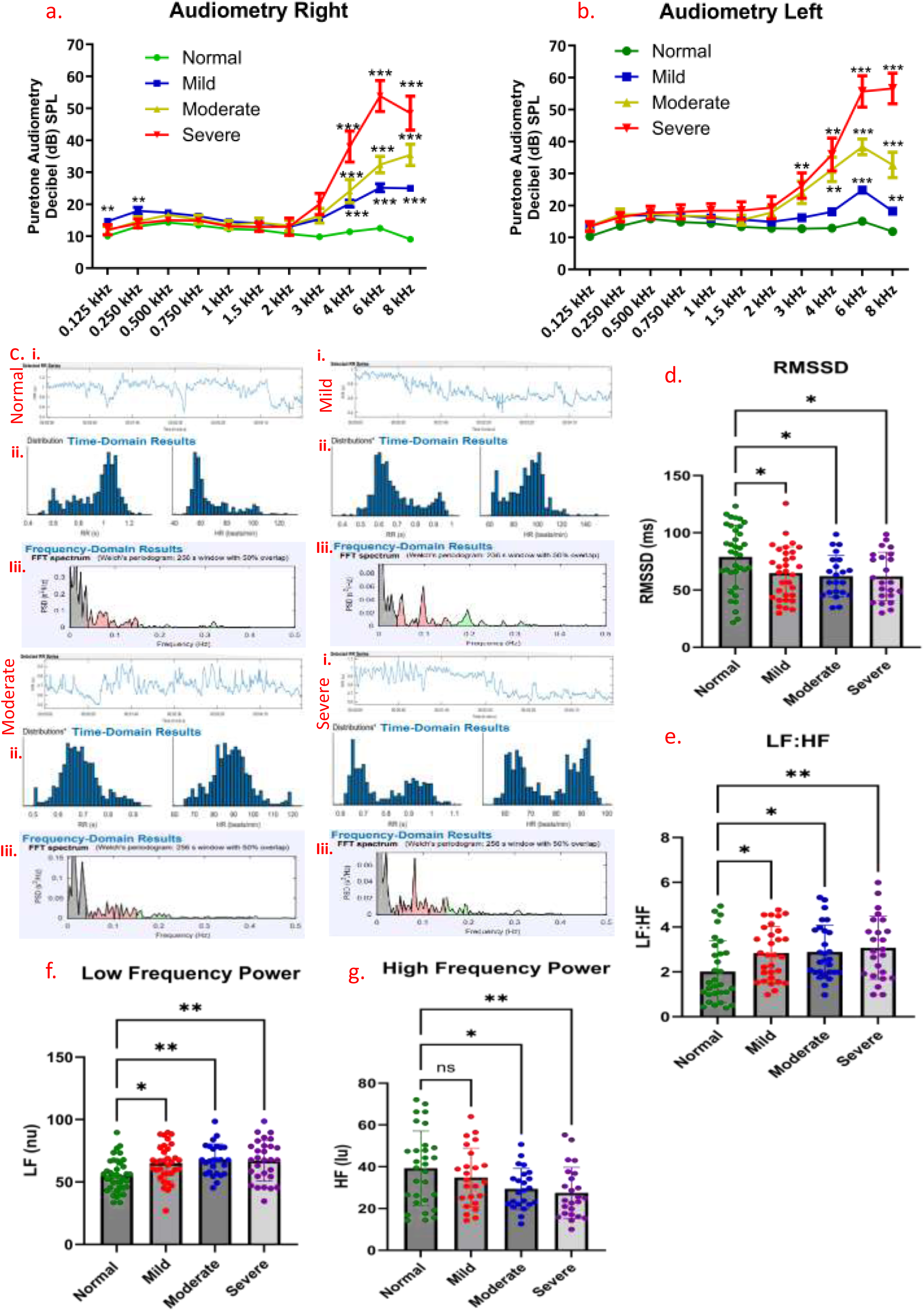
Assessment of Hearing and Physiological Markers in Response to Noise Exposure. (a) Right ear audiometry showing hearing thresholds across different frequencies for study participants. (b) Left ear audiometry depicting hearing thresholds across various frequencies. (c) Heart Rate Variability (HRV) data presented in three panels: (C-i) Normal RR interval measurements. (c-ii) Time domain analysis of HRV. (c-iii) Frequency domain analysis of HRV, illustrating variations in HRV across mild, moderate, and severe hearing impairment groups. (d) Analysis of the Root Mean Square of Successive Differences (RMSSD) in HRV data. (e) Low Frequency to High Frequency ratio (LF: HF) in HRV analysis. (f) High-frequency component of HRV. (g) Low-frequency component of HRV. Audiometry data represented in Mean±SEM Statistical analysis was performed using two-way ANOVA with post-hoc Dunnett’s test for multiple group comparisons against the normal group and HRV data in Mean±SD. Statistical significance was assessed using one-way ANOVA with post-hoc Dunnett’s test. Statistical significance is indicated where *p < 0.05, ** p<0.01 and p<0.001.

**Fig. 2.**
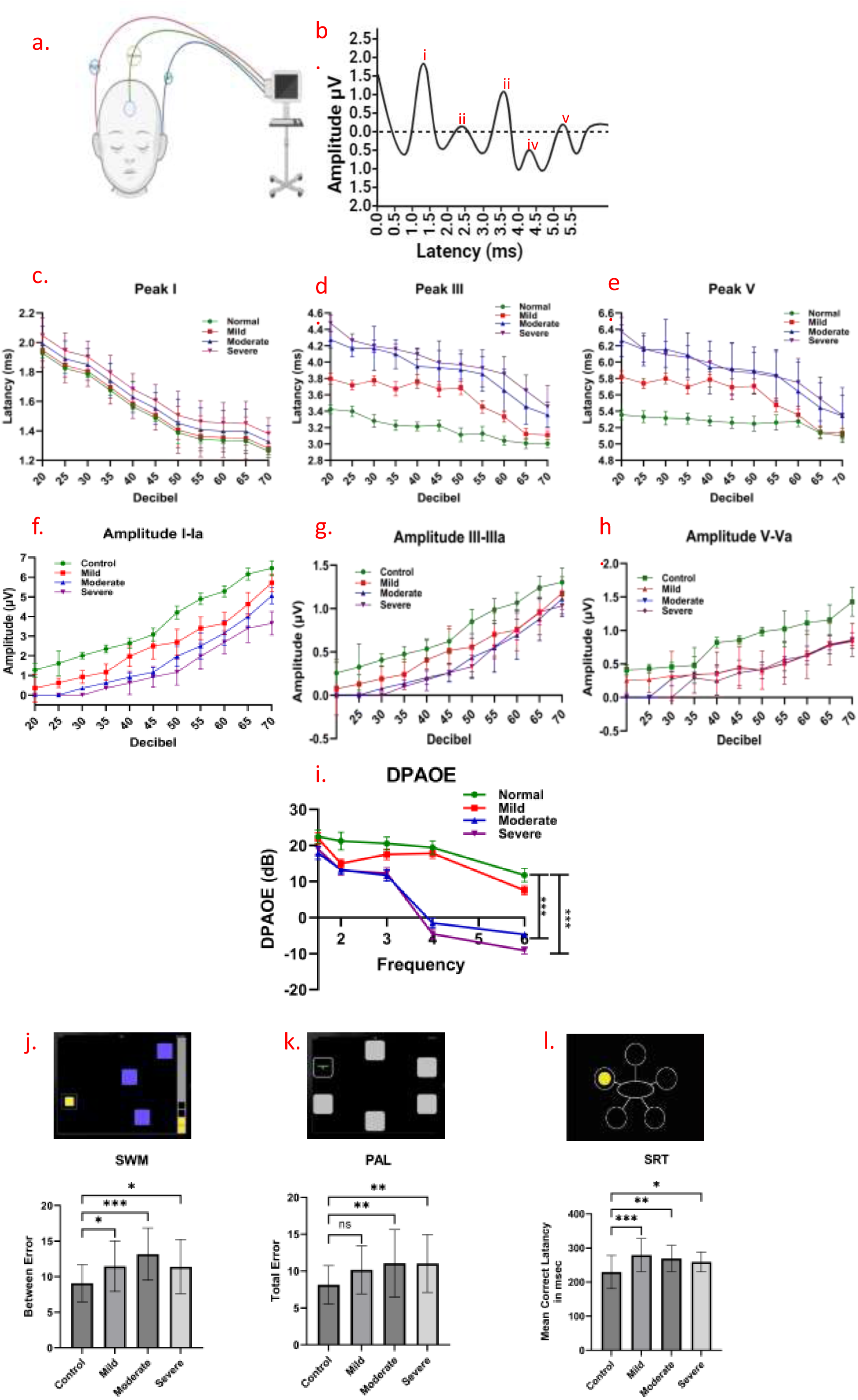
Auditory Brainstem Response (ABR) Data Analysis and Representative Measurements. (a) Representative Fig. showing the data acquisition setup for Auditory Brainstem Response (ABR) testing. (b) Representative data showing ABR peak and latency measurements. (c) Latency of Peak I (d) Latency of Peak III (e) Latency of Peak V (f) Amplitude between Peak I-Ia (g) Amplitude between Peak III-IIIa (h) Amplitude between Peak V-Va (b-h) presented as mean ± SD, across sound intensities ranging from 70 dB to 20 dB in 5 dB intervals. (i) Distortion Product Otoacoustic Emissions (DPOAE) data presented as mean ± SEM across 1.5kHz to 6kHz against f2-f1. (j) the SWM task, (k) PAL task, (l) Reaction Time. SWM, spatial working memory; PAL, Paired Associated Learning. Adapted with permission from Cambridge Cognition. amplitude & latency represented in Mean±SD Statistical analysis was performed using two-way ANOVA with post-hoc Dunnett’s test for multiple group comparisons against the normal group and DPAOE data in Mean±SD. Statistical significance was assessed using two-way ANOVA with post-hoc Dunnett’s test. Data for cognitive assessments are shown as mean ± SD, with statistical significance assessed using one-way ANOVA. Statistical significance is indicated where *p < 0.05, ** p<0.01 and p<0.001.

ABR measurements were taken after audiometry test. We used tone pips (5 ms duration, 1.5 ms rise/fall time) at frequencies from 1 to 8 kHz, with a stimulation rate of 19.1 Hz and a rejection level of ±20 μV. The stimuli were delivered through a tube connected to an electrostatic speaker in a custom-built sound chamber. ABR thresholds were determined by gradually reducing sound intensity in 5 dB steps from 70 dB SPL to 20dB SPL, averaging responses at each level over 1024 repeats with alternating stimulus polarity [8].

### Distortion product-oto-acoustic emissions (DPOAE)

In the DPOAE testing, the DP gram method was employed with an L1-L2 intensity difference of 10 dB (L1=65 dB and L2=55 dB) and a f1/f2 ratio of 1.22. The test duration was capped at 90 seconds. Frequencies tested ranged from 1,000 to 6,000 Hz, and their corresponding distortion products were considered present if the amplitude exceeded −10 dB and the signal-to-noise ratio was at least 6 dB. The pass/fail criterion for DPOAE testing utilized the 3-dB algorithm method to determine the presence of distortion products [16].

### Heart rate variability recordings

Heart rate variability (HRV) was measured in a sitting position at a room temperature of 23 ± 2 °C. After a 5-minute rest, each subject wore an iFeel HRV Sensor (iFeel Healthy Inc., Israel) that recorded data at 256 samples per second for 10 minutes. The data were examined for ectopic beats and artifacts, with artifact-free 5-minute segments analyzed using Kubios 2.2 (Kuopio, Finland). For the analysis, a 5-minute segment from the middle of the ECG recordings (between 180 and 480 seconds) was selected [37].

The study evaluated six HRV measures. Frequency domain indices included low-frequency power (LF, 0.04–0.15 Hz), high-frequency power (HF, 0.15–0.40 Hz), and total power (TP, 0.01–0.40 Hz). Time domain indices included the standard deviation of all normal R-R intervals (SDNN), the root mean square of successive differences between adjacent normal R-R intervals (r-MSSD), and the average SDNN across all 5-minute segments (SDNN index). LF power reflects both sympathetic and parasympathetic activity, while HF power indicates parasympathetic tone. SDNN and SDNN index represent overall HRV by capturing all cyclic components of variability, and r-MSSD measures short-term variability. Reductions in LF, HF, TP, SDNN, r-MSSD, and SDNN index suggest decreased sympathetic and parasympathetic activity, indicating impaired cardiac autonomic function. Details of all HRV parameters are summarized in Additional Table 1.

**Table 1.**
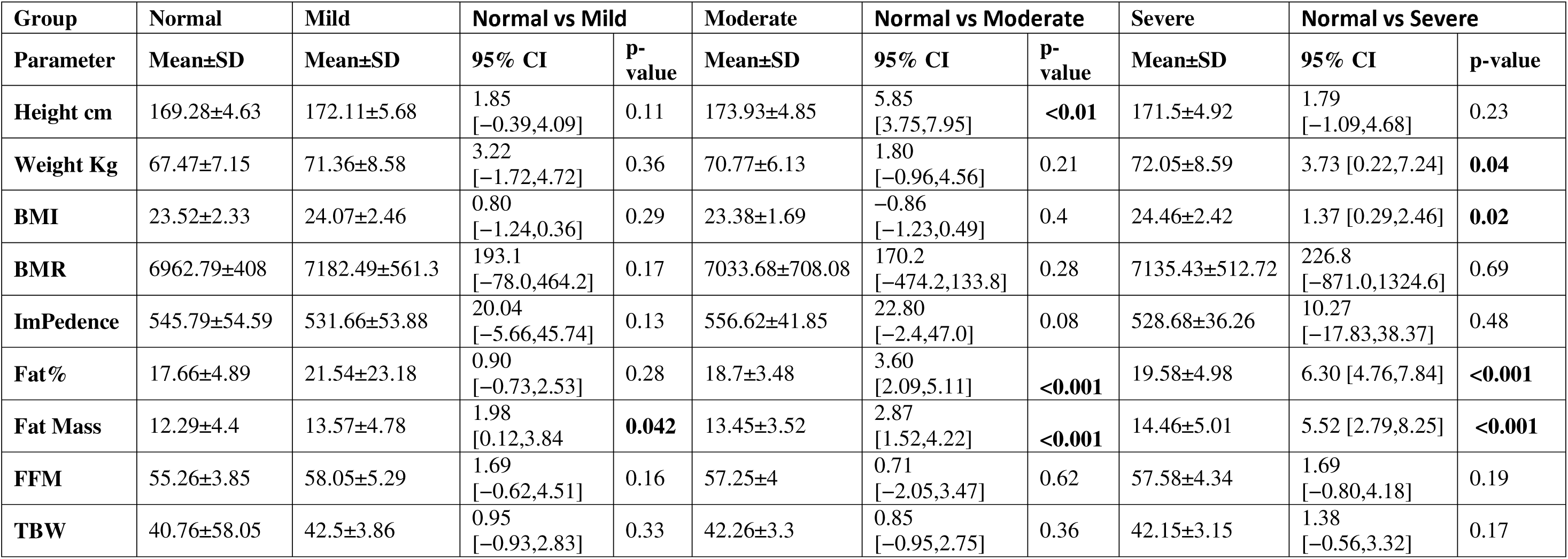
Summary of Physiological Parameters (Body Compostion) provides a comprehensive overview of key metrics related to body composition and metabolic rate, aiding in the assessment of individual health and physiological status.

| Group | Normal | Mild | Normal vs Mild |  | Moderate | Normal vs Moderate |  | Severe | Normal vs Severe |  |
| --- | --- | --- | --- | --- | --- | --- | --- | --- | --- | --- |
| Parameter | Mean±SD | Mean±SD | 95% CI | P-value | Mean±SD | 95% CI | P-value | Mean±SD | 95% CI | p-value |
| Height cm | 169.28±4.63 | 172.11±5.68 | 1.85<br>[-0.39,4.09] | 0.11 | 173.93±4.85 | 5.85<br>[3.75,7.95] | <0.01 | 171.5±4.92 | 1.79<br>[-1.09,4.68] | 0.23 |
| Weight Kg | 67.47±7.15 | 71.36±8.58 | 3.22<br>[-1.72,4.72] | 0.36 | 70.77±6.13 | 1.80<br>[-0.96,4.56] | 0.21 | 72.05±8.59 | 3.73 [0.22,7.24] | 0.04 |
| BMI | 23.52±2.33 | 24.07±2.46 | 0.80<br>[-1.24,0.36] | 0.29 | 23.38±1.69 | -0.86<br>[-1.23,0.49] | 0.4 | 24.46±2.42 | 1.37 [0.29,2.46] | 0.02 |
| BMR | 6962.79±408 | 7182.49±561.3 | 193.1<br>[-78.0,464.2] | 0.17 | 7033.68±708.08 | 170.2<br>[-474.2,133.8] | 0.28 | 7135.43±512.72 | 226.8<br>[-871.0,1324.6] | 0.69 |
| ImPedence | 545.79±54.59 | 531.66±53.88 | 20.04<br>[-5.66,45.74] | 0.13 | 556.62±41.85 | 22.80<br>[-2.4,47.0] | 0.08 | 528.68±36.26 | 10.27<br>[-17.83,38.37] | 0.48 |
| Fat% | 17.66±4.89 | 21.54±23.18 | 0.90<br>[-0.73,2.53] | 0.28 | 18.7±3.48 | 3.60<br>[2.09,5.11] | <0.001 | 19.58±4.98 | 6.30 [4.76,7.84] | <0.001 |
| Fat Mass | 12.29±4.4 | 13.57±4.78 | 1.98<br>[0.12,3.84] | 0.042 | 13.45±3.52 | 2.87<br>[1.52,4.22] | <0.001 | 14.46±5.01 | 5.52 [2.79,8.25] | <0.001 |
| FFM | 55.26±3.85 | 58.05±5.29 | 1.69<br>[-0.62,4.51] | 0.16 | 57.25±4 | 0.71<br>[-2.05,3.47] | 0.62 | 57.58±4.34 | 1.69<br>[-0.80,4.18] | 0.19 |
| TBW | 40.76±58.05 | 42.5±3.86 | 0.95<br>[-0.93,2.83] | 0.33 | 42.26±3.3 | 0.85<br>[-0.95,2.75] | 0.36 | 42.15±3.15 | 1.38<br>[-0.56,3.32] | 0.17 |

### Automated neuropsychological assessment

Volunteers were seated to complete a series of three computerized neuropsychological tests from the CANTAB Battery, as outlined in the CANTAB manual protocols Cognitive score. Following a brief practice session with the touch screen, the three tests were administered. The entire testing process, including instructions, took approximately 50 minutes. The order of the tests was counterbalanced across participants [38].

### Spatial Working Memory (SWM)

Spatial working memory (SWM) is designed to evaluate working memory by requiring participants to search for tokens hidden in colored boxes on a screen. Here’s a detailed breakdown of how the test works and the measures it provides: Participants are shown several colored boxes on the screen, with a single token hidden in one box per trial. The goal is to find the blue tokens. Each box should ideally be searched only once per trial to maximize efficiency. Returning to a box that has already been checked or emptied is penalized. Occur when a participant searches the same box more than once within a single trial. This reflects an inefficiency or failure to remember which boxes have already been checked. Occur when a participant returns to a box that has already been emptied in the course of the trial. This indicates a failure to remember that the box has been checked and is no longer a viable option. Instances where an error can be classified as both a within error and a between error simultaneously. Two major indices were presented: strategy utilization: the number of search sequences starting with a novel box in both 6– and 8-box problems, and errors in total and three different levels of difficulty (4-, 6-, and 8-box problems): the total errors were calculated based on the between errors, within errors, and double errors of particular box problems (i.e., between errors + within errors – double errors) [39].

### Paired Associates Learning (PAL)

The PAL test assesses visual memory and new learning, with particular sensitivity to changes in the temporal and frontal lobes of the brain. In this test, participants are first shown six boxes on a screen, which are opened in a random sequence to reveal their contents. Some of these boxes contain patterns. Afterward, a pattern is displayed in the center of the screen, and the participant must identify which box holds that pattern. If the participant correctly identifies all the patterns, they advance to the next stage, which involves eight boxes. If not, the test ends. The difficulty of the test increases as participants must remember between two and eight patterns.

The PAL test includes the following outcome measures: i. PAL Total Errors Adjusted: This measures the total number of errors made across all stages, adjusted for any stages not attempted due to earlier failures. ii. PAL Total Errors (Six Shapes, Adjusted): This measures the total number of errors made in the eight-stage pattern test, with an adjustment for subjects who did not reach this stage. S Akter[39].

### Reaction Time (RTI)

The RTI test evaluates how quickly a person can respond to visual stimuli on a screen, where the stimuli can be either predictable (simple reaction time) or unpredictable (choice reaction time). The test consists of five progressively complex stages, requiring a series of responses. In this test, a yellow spot may appear in either a single location or in one of five different locations. Participants respond by either pressing a pad, touching the screen, or using both methods. The RTI test uses the following measures: i. RTI Five-Choice Reaction Time: This measure how quickly the participant releases the press pad when a stimulus appears in one of the five possible locations. ii. RTI Five-Choice Movement Time: This measures the duration it takes for the participant to touch the stimulus on the screen after they have released the press pad[39, 40]

### Body Composition and Basal Metabolic Rate (BMR) Assessment

Participants fasted for a minimum of 8 hours before the assessment to eliminate the influence of recent food intake on body composition measurements. Body composition measurement equipment (e.g., bioelectrical impedance analyzer) was used. Participants were seated and at rest for at least 5 minutes before measurement. Electrodes were attached to designated body sites (e.g., hands, feet) following the device’s protocol. The measurement was conducted as per the device’s standard operating procedure, recording body fat percentage, lean body mass, and total body water. All data summarized in Table 1.

### Sample Collection and Processing

All participants received comprehensive instructions on blood collection prior to the procedure. Blood samples were drawn during the fasting period, between 0600 and 0800 hours. Venous blood was collected from the antecubital vein into heparin-treated vials while the subjects were in a resting state. To ensure anonymity, samples were coded and processed according to a standardized protocol. Blood was centrifuged at 1000g for 15 minutes to separate the plasma, which was then aliquoted and stored at −20°C. Upon arrival at the laboratory, the samples were further stored at −80°C until analysis.

## Biochemical/ELISA estimation

### Cardiac Parameters

To quantify HDL and LDL/VLDL cholesterol levels, use the EnzyChrom™ HDL and LDL/VLDL Assay Kit (CatLog EHDL-100). Measure total cholesterol using the EnzyChrom™ Cholesterol Assay Kit (CatLog E2CH-100). Determine triglyceride levels with the EnzyChrom™ Triglyceride Assay Kit (CatLog ECCH-100). Assess glucose concentration using the EnzyChrom™ Glucose Assay Kit (CatLog EBGL-100). Measure CK-MB levels with the Human CK-MB ELISA Kit (CatLog 1005-P1). Quantify C-Troponin using the Human TNNI3/cTn-I (Troponin I Type 3, Cardiac) ELISA Kit (CatLog E-EL-H0649).

### Oxidative Stress Parameters

Measure 4-HNE levels using the ELISA kit (CatLog E-EL-0128). Quantify Vitamin B12 with the ELISA kit (CatLog EIA5848R). Assess cortisol concentration using the ELISA kit (CatLog EIA1887R). Determine superoxide dismutase (SOD) activity with the EnzyChrom™ Superoxide Dismutase Assay Kit (CatLog ESOD-100). Utilize the appropriate assay method for 8-HdOG measurement. Measure malondialdehyde (MDA) using the EnzyFluo™ Myeloperoxidase Assay Kit (CatLog EMPO-100). Quantify glutathione (GSH) and its oxidized form (GSSG) using the EnzyChrom™ GSH/GSSG Assay Kit (CatLog EGTT-100).

### Inflammatory Markers

Use the appropriate assay method for IL-1α and IL-1β measurements. Quantify TNF-α levels with the ELISA kit (CatLog EIA4641) and assess IL-6 concentration using the ELISA kit (CatLog EIA4640). Measure IFN-γ and IL-4 using the appropriate assay methods. Quantify IL-10 with the ELISA kit (CatLog EIA4699). Assess MIP-1α and MIP-1β levels using the relevant assay methods. Determine C-reactive protein (CRP) levels with the ELISA kit (CatLog EIA3954). Measure homocysteine using the appropriate assay method. Quantify prolactin with the ELISA kit (CatLog EIA1291). Serum levels of the experimental biomarker inflammatory panel (sAXL, sTyro3, YKL-40) used from R&D system Inc. as per manufacturer instruction.

### Neurotransmitters

Quantify adrenaline using the DLD Diagnostika GmbH (Hamburg, Germany) assay (CatLog EA604/96). Measure noradrenaline with the DLD Diagnostika GmbH (Hamburg, Germany) assay (CatLog EA610/96). Determine dopamine levels using the DLD Diagnostika GmbH (Hamburg, Germany) assay (CatLog EA608/96). Utilize the appropriate assay method for serotonin measurement. Measure B-type natriuretic peptide (BNP) and atrial natriuretic peptide (ANP) with the relevant assay methods.

### Sample preparation and processing for iTRAQ

Approximately 100 µL plasma from each individual (total 20 subject) per group was pooled to yield four different pools (Normal, mild, moderate and severe) of 2 ml plasma each. For iTRAQ labelling, Samples were processed with the ASKs kit to reduce the level of serum albumin. Total protein content was determined with ToPA Bradford Protein Assay kit. 50µg of total protein from each sample was reduced, alkylated, and precipitated to remove interfering substances. The precipitated proteins were re-suspended in digestion buffer, and trypsin digestion was performed overnight. The digested samples were individually labelled with iTRAQ reagents as follows: Normal (114), Mild (115) Moderate (116), Severe (117) For MudPIT, each sample was fractionated by SCX. The fractions eluted at 75mM, 150mM and 450mM Ammonium Acetate were collected and analyzed individually by nano LC-MS/MS, and the data obtained combined. Prior to MS analysis, samples were desalted using ZipTip and the desalted sample dried in a speedvac, then re-suspended in the appropriate mobile phase for nano-LC-MS/MS analysis [41].

### MOLDI-TOF/TOF analysis and protein identification

Each MudPIT fraction was subjected to in-line Reverse Phase (RP) chromatography and tandem mass spectrometry to identify proteins in each fraction. Briefly, each sample was loaded onto a PicoFrit C18 nanospray column using a Thermo Scientific Surveyor Autosampler operated in the no-waste injection mode. The flow rate was 600nl/min. Peptides were eluted from the column using a linear acetonitrile gradient from 5 to 45% acetonitrile over 180 minutes followed by high and low organic washes for another 30 minutes into the LTQ XL mass spectrometer (Thermo Scientific) via a Nano spray source. The spray voltage was set to 1.8kV and the ion transfer capillary was set at 180°C. A data-dependent Top 5 method was used where a full MS scan from m/z 300-1600 was followed by MS/MS scans on the five most abundant ions. Raw data files were searched against the most recent database for human downloaded from UniProt using the Mud PIT option in Proteome Discoverer 1.4 (Thermo Scientific) and the SyQuest HT search algorithm. For protein identification results, only peptides identified with high confidence were used. Trypsin was the selected enzyme allowing for up to two missed cleavages per peptide. Methylthiolation of cysteine, N terminal iTRAQ4-plex, and Lysine iTRAQ4-plex were used as the static modifications whereas oxidation of Methionine was used as the variable modification. For confidence, the Percolator algorithm was used for PSM (peptide-spectrum match) validation in database searches. The False Discovery Rate (FDR) threshold calculated in Proteome Discoverer Percolator when high confidence peptide was used for protein identification is 0.01. The following comparisons were made for relative protein quantitation (Additional Table 2): 114/113 Mild/Control, 115/113 Moderate/Control, 116/113 Severe/Control. Total 176 proteins were identified unequivocally using only high confidence peptides. A fold-change threshold of 1.2 was used for up/down regulation. Thus, iTRAQ Ratio >1.2 were classified as up-regulated, <0.67 were classified as down-regulated. Ratios from 0.67–1.2 were considered no changes. Ratios above 100 exceed the maximum allowed threshold that is reasonably expected. Fig. 6a and 6b show representative MS/MS spectra for two unique peptides (EDLIWELLNQAQEHFGK and DLIWELLNQAQEHFGK) from the two higher coverage proteins (Protein Serotransferrin and Protein Transthyretin). Supplementary table 2 gives details of the protein IDs and the iTRAQ ratios. The Table includes all proteins identified whether there was an iTRAQ ratio.

### Functional enrichment analysis of the DEGs Gene Ontology (GO) and Kyoto Encyclopedia of Genes and Genomes (KEGG) pathway analysis of DEGs

The Database for Annotation, Visualization, and Integrated Discovery (DAVID) was utilized as a functional annotation tool to gain insights into the biological implications of the gene lists [DAVID: https://david.ncifcrf.gov/]. Gene Ontology (GO), which encompasses biological process (BP), molecular function (MF), and cellular component (CC), was employed for functional studies of the genomic data at a large scale [42]. Additionally, the Kyoto Encyclopedia of Genes and Genomes (KEGG) was utilized as a pathway-related database to understand the molecular networking of genes and molecules [43]. GO-BP terms with an adjusted p-value below 1e-5 were considered overrepresented, and KEGG pathways with a false discovery rate (FDR) below 0.05 were deemed significantly enriched. To perform these analyses, we employed the online DAVID tool [44].

### Construction of the PPI network of the overlapping DEGs

Protein-protein interactions (PPIs) were analyzed to explore the interactions and molecular functions of proteins. PPIs provide valuable insights into the underlying mechanisms of protein function[45]. The Search Tool for the Retrieval of Interacting Genes (STRING) database (http://string-db.org/) was utilized to gather information on predicted and experimentally validated PPIs within a specific cellular context. For this analysis, the dysregulated differentially expressed genes (DEGs) were mapped onto the STRING database to construct a PPI network. The PPI network did not consider text mining or database-based interaction scores. A high combined score threshold was employed as the cutoff value to determine significant protein pairs in the network. Finally, based on the topologies determined by CytoHubba, we extracted the significant protein total of 34 genes [46].

### Validation of Proteins Using ELISA Kit and /or western blot in Plasma or Serum Samples

To validate the presence and quantify the levels of specific proteins in plasma or serum samples, an Enzyme-Linked Immunosorbent Assay (ELISA) kit was utilized. The proteins of interest for this validation included: ORM1 (Alpha-1-acid glycoprotein) ELH-ORM1-1, CLU Clusterin ELH-Clusterin-1, APCS (Serum Amyloid P Component) ELH-APCS-1, LIMK1 (LIM Kinase 1), VTN (Vitronectin) ELH-VTN-1, TTR (Transthyretin) ELH-TTR-1, RBP4 ELH-RBP4-1, TF ELH-Trfrn-1 and CLEC3 (C-Type Lectin Domain Family 3) ELH-CLEC3B-1 all kit from Raybiotech. All experiment performs according to manufacture instruction.

### Statistical analysis using GraphPad Prism 10.3.3

Results are expressed as the means ± SD and/or means ± SEM. Two-way ANOVA (with Dunnett’s correction for comparison of multiple means) was used for audiometry, DPAOE, and ABR test (both amplitude, latency) (Prism for Windows, version 10.3.3, GraphPad Software Inc.). One-way ANOVA (with Dunnett’s correction for comparison of multiple means) or, where appropriate, equivalent non-parametric test (Dunnett/Kruskal-Wallis multiple comparison) was used for comparisons in HRV data (RMSSD, HF, LF LF/HF), other serum/plasma parameters (Cardiac, Oxidative, Neurotransmitter and inflammatory parameter. p-values<0.05 were considered as statistically significant and are either provided in the Figures or by symbol legends of tables.

Use SRplot to compute Pearson correlation coefficients for all Cardiac, Oxidative stress, Inflammatory and neurotransmitter parameter pairs. Evaluate the strength and direction of each correlation: +1 represents a perfect positive correlation, –1 indicates a perfect negative correlation, and 0 signifies no correlation. Conduct significance testing for each coefficient to determine if the results are statistically significant using a p-value threshold of 0.05. In Fig 5, summarize the key correlations and discuss their implications for understanding the impact of hearing impairment on clinical parameters Pearson correlation, (insig p without *, sig p with *) [47]

## Results

### Audiometry Analysis: Impact of Aircraft Noise Exposure on Hearing Sensitivity

Of the 621 participants in the study, 345 subjects were found to have significant threshold shift; 165 subjects had mild, 102 moderate and 78 severe hearing impairment when auditory thresholds were calculated as average of 125Hz, 500 Hz, 750Hz, 1000 Hz, 2000 Hz, 3000 Hz, 4000Hz, 6000 Hz and 8000 Hz frequency (WHO guidelines). According to OSHA guidelines, a notch/dip at 3000, 4000, 6000 and 8000 Hz is indicative of the commencement of auditory impairment. It was found that 276 no. of personnel having normal hearing the defence personnel showed normal hearing, when calculated as average of 500 Hz, 1000 Hz, 2000 Hz, and 4000 Hz but they had started developing the early changes of NIHL in the form of notching at 6000 Hz frequency (Fig. 1a right ear, Fig. 1b left ear) described as threshold shifts. studied showed threshold shifts >25 dB in 3 kHz frequency in Left ear and >25 dB in 2kHz in right ear. Overall Right ear showed in interaction F (30, 6787) = 111, P<0.0001 in Row Factor F (10, 6787) = 419, P<0.0001 and in Column Factor (3, 6787) = 693 P<0.0001 similarly in left ear F (DFn, DFd) Interaction F (30, 1540) = 21.5, P<0.0001; Row Factor F(10, 1540) = 73.2, P<0.0001; Column Factor F (3, 1540) = 175, P<0.0001. Additional Table 3 provides a comprehensive breakdown of the statistical analyses for audiometric thresholds.

### ABR Analysis Impact of Aircraft Noise Exposure on Hearing Sensitivity

Our study utilized Auditory Brainstem Response (ABR) testing to evaluate auditory function in human subjects exposed to noise. Wave I, originating from the peripheral portion of cranial nerve VIII, appeared at approximately 1.5 milliseconds (msec). Wave II, originating from the more proximal section of the nerve, was recorded at around 2.5 msec. Wave III, generated at the cochlear nucleus, occurred at about 3.5 msec, while Wave V, associated with the lateral lemniscus/inferior colliculus, was observed around 5.5 msec.

Dunnett’s multiple comparisons test showed significant differences between Normal and various severity levels (Mild, Moderate, Severe) at all decibel levels, with all p-values <0.001. At 50 dB, wave I the mean latency differences were –0.0197 (95% CI: –0.0644 to 0.0251) p=0.62 for Mild, –0.0647 (–0.113 to –0.0165) p=0.004 for Moderate, and –0.119 (–0.180 to –0.0578) p<0.001 for Severe, indicating a clear trend of increasing amplitude differences with greater severity (Fig. 2c). Wave III amplitude changes at 50 dB were significant: –0.57 (95% CI: –0.625 to –0.516) for Mild, –0.794 (95% CI: –0.852 to –0.736) for Moderate, and –0.854 (95% CI: –0.928 to –0.780) for Severe, all with p-values <0.001 (Fig. 2d). These findings suggest significant functional hearing deficits not always detectable through standard audiometric thresholds. Some subjects exhibited increased cochlear synaptopathy, indicative of synaptic degeneration and hair cell changes due to noise exposure. Additional Table 4 provides a comprehensive breakdown of the statistical analyses for wave I-V latency.

For Amplitude I-Ia, the differences at 50 dB were 1.51 (95% CI: 1.32 to 1.70) for Mild, 2.23 (95% CI: 2.03 to 2.43) for Moderate, and 3.04 (95% CI: 2.78 to 3.30) for Severe (Fig. 2f). For Amplitude III-IIIa, at 50 dB, the differences were 0.294 (95% CI: 0.217 to 0.370) for Mild, 0.415 (95% CI: 0.333 to 0.498) for Moderate, and 0.52 (95% CI: 0.415 to 0.625) for Severe (Fig. 2g). Additional Table 5 provides a comprehensive breakdown of the statistical analyses for wave Ia-Va amplitude. These results illustrate the progressive impact of noise exposure on auditory function, highlighting the importance of ABR in detecting cochlear synaptopathy and functional deficits beyond traditional audiometric assessments.

### DPOAE Analysis: Impact of Aircraft Noise Exposure on Hearing Sensitivity

DPOAE amplitude data provided an indirect measure of OHC integrity at frequency specific regions of the inner ear. DPOAE supra threshold amplitudes were recorded at 2f1-f2. The functional status of cochlear Outer Hair Cells was checked by the recording of Distortion Product-OAEs in the normal, mild, moderate subjects to confirm OHC function. The results obtained were almost like those obtained by pure tone audiometry. The DPOAE amplitudes of the personnel having normal hearing personnel ears are maximal and in order to mild to severe for the personnel with exposed ears the DPOAE amplitudes decreased in column F (3, 1080) = 2558, P<0.001. The DPOAE amplitudes in Exposed ears were significantly smaller than those in normal (P < 0.0001). DPOAE testing revealed a significant difference at 1.5kHz to 6 kHz (P = 0.001) in the mild; 2kHz to 6 kHz (P = 0.001) in the moderate and severe exposed ear and at 2kHz to 6 kHz (P = 0.001) in the mild hearing ear (Fig. 2i).

### Heart Rate Variability Assessment in Response to Aircraft Noise Exposure

In the HRV (Heart Rate Variability) test, we observed a significant decrease in the levels mild 14.2 (95%CI 0.474 27.9) p<0.04, moderate 16.8 (95%CI 1.54 32.0) p<0.03, severe 16.8 (95%CI 1.98 32.1) p<0.02) of RMSSD (Fig. 1d) and high-frequency power (HF nu) across subjects with moderate (p<0.04), and severe (p<0.008) exposure to aircraft noise (Fig. 1g). The reduction in RMSSD indicates a decline in parasympathetic nervous system activity and overall vagal tone, reflecting a diminished ability to regulate heart rate variability in response to stress. Similarly, the decrease in HF nu suggests a reduction in the parasympathetic component of heart rate variability. Conversely, there was a significant increase in the LF: HF mild (p<0.04), moderate (p<0.03), severe (p<0.009) (Fig. 1e) and LF nu mild (p<0.01), moderate (p<0.002), severe (p<0.003) (Fig. 1f), indicating a shift towards higher sympathetic nervous system dominance relative to parasympathetic activity. This shift implies an overall increase in sympathetic activity and a reduced balance between sympathetic and parasympathetic influences, highlighting the impact of chronic noise exposure on autonomic regulation and cardiovascular health.

### Cognitive Assessment of the Impact of Aircraft Noise Exposure: Evaluating Spatial Working Memory, Paired Associates Learning, and Simple Reaction Time

We administered the CANTAB battery to assess cognitive parameters, specifically evaluating Spatial Working Memory (SWM), Paired Associates Learning (PAL), and Simple Reaction Time (SRT) across subjects with normal hearing, mild hearing impairment, moderate hearing impairment, and severe hearing impairment. The results revealed a significant increase in the number of errors in the SWM task among subjects with mild, moderate, and severe hearing impairments, indicating impaired spatial working memory Mild –2.41 (–4.37-0.260) p=0.02; Moderate –4.10 (–6.26 to –1.95) <0.001; Severe –2.90 (–4.5 to –0.191) p=0.03 (Fig. 2j). Similarly, PAL performance showed increased total errors F 4.10 (3, 112); p=0.008 and longer mean correct latencies measured in milliseconds across all levels of hearing impairment in mild CI95% was –2.03 (–4.32 –0.252) p value 0.09 in moderate –2.93 (–5.22 –0.645) p value 0.008 and in severe –2.90 (–5.18 –0.611) p value 0.009 suggesting deterioration in associative learning and memory recall (Fig. 2k). The SRT results also demonstrated longer reaction times Mild –50 (–76.3 to –23.8) p<0.001; Moderate –39.9 (–66.2 to –13.6) <0.001; Severe –30.0 (–56.2 to –3.71) p=0.02 (Fig. 2l) in individuals with hearing impairments, reflecting delays in simple cognitive processing. These findings collectively highlight that hearing impairment is associated with significant declines in cognitive function, as evidenced by increased errors and longer response times across multiple cognitive tasks.

### BCA Anthropometric characteristics and body composition of Indian males stratified by hearing groups

BCA data analysis revealed significant differences in several parameters across severity groups (Normal, Mild, Moderate, Severe). Height significantly differed only between Normal and Moderate groups (Mean difference = 5.85 cm, p < 0.01). Weight and BMI were notably higher in the Severe group compared to Normal (Mean differences = 3.73 kg and 1.37, p = 0.04 and p = 0.02, respectively). Fat Percentage and Fat Mass increased significantly with severity, especially in the Severe group (Mean differences of 6.30% and 5.52 kg, p < 0.001). Basal Metabolic Rate, Impedance, Fat-Free Mass, and Total Body Water showed no significant variations across groups all parameter summarized in Table 1.

### Redox Signaling: Assessing Oxidative Stress and Antioxidant Responses in the Context of Aircraft Noise Exposure

Noise is mainly linked with the phenomenon/occurrence of Oxidative stress in an individual. Hence, the levels of oxidative stress parameters and antioxidants in human plasma were analysed to validate this postulate. C-Reactive Protein (CRP) –4.47 (–5.70 –3.24); –1.56 (– 2.80 –0.332); –1.40 (–2.65 –0.143); an acute-phase reactant, also showed higher levels, indicating systemic inflammation (Fig. 3a). The levels of cortisol increased with increase in hearing a stress hormone, was elevated, reflecting an increased physiological stress response associated with prolonged noise exposure in mild –6.86 (–10.3 –3.45), (p<0.001); moderate –2.88 (–8.70 –1.97), (p<0.001); and severe –4.43 (–10.8 –3.81), (p<0.001) (Fig. 3b) likewise, the MDA levels, markers of lipid peroxidation also increased with increase in hearing impairment in mild and moderate group but normalized in the severely hearing-impaired group mild –4.14 (–5.37 –2.91), (p<0.001); moderate –2.88 (–4.11 –1.66), (p<0.001); and severe –4.43 (–5.67 –3.19), (p<0.001) (Fig. 3d). The levels of oxidative DNA damage were found to increase in all exposed groups from –13.0(–16.8 – 9.14); –6.44 (–10.3 –2.62); –9.21(–13.1-5.29) mild to severe. (Fig. 3c).

**Fig. 3.**
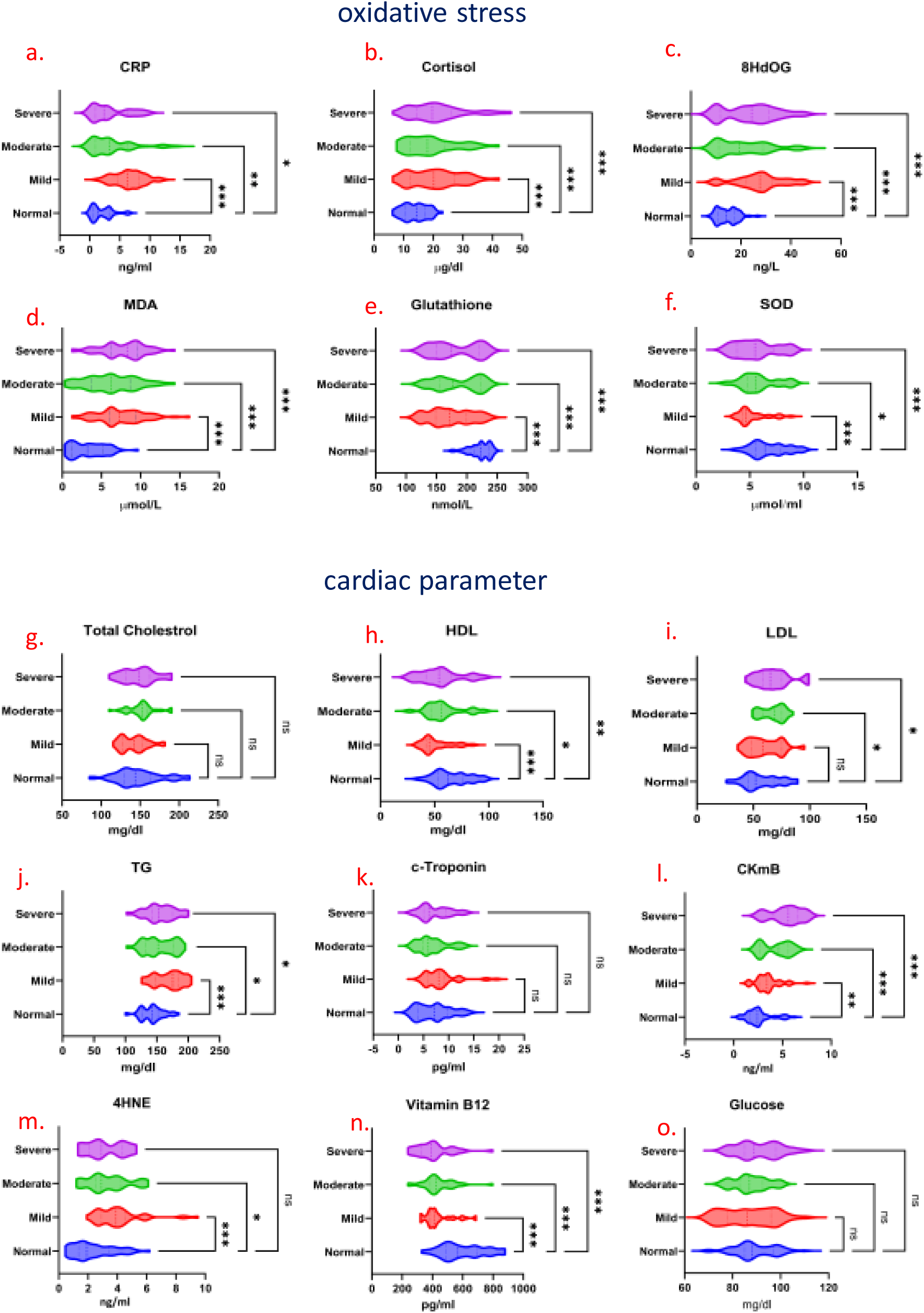
Oxidative Stress and Cardiac Parameters Across Hearing Impairment Groups. Oxidative stress parameters: (a) C-reactive protein (CRP) levels, (b) Cortisol levels, (c) 8-Hydroxydeoxyguanosine (8-OHdG) levels, (d) Malondialdehyde (MDA) levels, (a-d) shown as mean ± SD, with elevated levels observed across mild to severe hearing impairment groups. (e) Glutathione levels, (f) Superoxide Dismutase (SOD) levels, (e-f) data presented as mean ± SD, with reduced levels observed in the mild to severe hearing impairment groups. **Cardiac parameters:** (g) Cholesterol (h) High-Density Lipoprotein (HDL) levels, (i) Low-Density Lipoprotein (LDL) levels, (j) Triglycerides (TG) levels, (k C-Troponin level, (l) Creatine Kinase MB (CK-MB) levels, (m) 4-Hydroxy-2-nonenal (4-HNE) levels, (n) Vitamin B12 levels, depicted as mean ± SD, with (o) Cholesterol, Troponin, and Glucose levels. All data represented as mean ± SD with. statistical significance assessed using one-way ANOVA followed by Dunnett’s post-hoc test for comparisons against the normal hearing group. Where (i, j, l, m) showing increased levels from mild to severe hearing impairment groups, HDL & Vitamin B12 reduced levels from mild to severe hearing impairment groups, Cholesterol, C-Troponin, and Glucose levels with no significant change across the hearing impairment groups. Statistical significance is indicated where *p < 0.05, ** p<0.01 and p<0.001.

Since antioxidants offer protection from the damage caused by oxidative stress, the activity/expression of various antioxidants such as reduced glutathione and superoxide dismutase was observed. The levels of reduced glutathione declined with increase in the severity of hearing impairment from 48.8 (35.0 62.7) p p<0.001; 36.6 (22.4 50.8) p<0.01; 41.0 (26.7 55.3) p<0.001 mild, moderate, and severe group as compared to normal (Fig. 3e). Similarly, the levels of superoxide dismutase also declined with increasing hearing impairment 1.52 (0.823 2.22) p p<0.001; 0.915 (0.150 1.68) p<0.01; 1.21 (0.446 1.98) p<0.001 mild, moderate, and severe group (Fig. 3f). It is observed that noise results in an increase in lipid peroxidation as evinced by increase in MDA concentration in plasma, coupled with a decrease in the erythrocytes activity of SOD, reduced GSH in blood, and plasma total antioxidant status in all the hearing-impaired groups.

In our study, analysis of serum/plasma samples revealed significant increases in various oxidative stress parameters among individuals with mild, moderate, and severe hearing lo ss due to chronic noise exposure suggests that noise exposure leads to significant oxidative stress, causing cellular and genetic damage was also significantly increased mild, moderate, and severe group, indicating heightened oxidative stress and damage to cell membranes. These findings collectively demonstrate that chronic noise exposure induces considerable oxidative stress and inflammation, contributing to the physiological changes observed in hearing loss.

### Modulation of cardiac Expression in Response to Aircraft Noise Exposure

Chronic noise exposure significantly impacts cardiac parameters, as evidenced by elevated levels of LDL (low-density lipoprotein) –2.72 (–9.37 3.94) p=0.67; –10.3 (–18.9 –1.70) p<0.01; –10.6 (–19.2 –2.05) p<0.01 in mild, moderate and severe (Fig. 3i) and CK-MB (creatine kinase-MB) across all levels of hearing loss severity—mild, moderate, and severe –0.872 (–1.48 – 0.262) p=0.002; –1.69 (–2.57 –0.806) p<0.001; –2.75 (–3.63 –1.87) p<0.001 (Fig.3l). Elevated LDL levels are associated with increased risk of atherosclerosis and cardiovascular diseases, reflecting potential adverse effects of chronic noise on lipid metabolism. Elevated CK-MB, a marker of myocardial injury, suggests possible subclinical damage to cardiac tissue due to prolonged noise stress. Additionally, 4-HNE (4-hydroxy-2-nonenal), a marker of oxidative stress, was elevated only in individuals with mild and moderate hearing loss –1.69 (–2.36 –1.02) p<0.001; –1.03 (–1.99 –0.0661) p<0.03; –0.848 (–1.81 0.116) p=0.10, indicating early oxidative damage (Fig. 3m). Conversely, HDL (high-density lipoprotein) levels were decreased across all exposure groups mild 14.2 (6.94 21.5) p<0.001; moderate7.66 (0.137 15.2) p=0.04; severe10.5 (2.94 18.0) p=0.003, which further implicates disrupted lipid profiles and increased cardiovascular risk (Fig. 3h). Vitamin B12 levels were also reduced in all subjects, potentially reflecting impaired nutritional status or increased oxidative stress mild 162 (108 216) p<0.001; moderate 160 (100 219) p<0.001; severe 191 (132 250) p<0.001 (Fig. 3n). Total cholesterol (Fig. 3g) p=0.40; p=0.69; p=0.99 and glucose levels p=0.35, p=0.70, p=0.71(Fig. 3o) and c-troponin p=0.08, p=0.99, p=0.96 remained unchanged from mild to severe, suggesting that the observed changes are specific to certain lipid and cardiac markers rather than general metabolic alterations. These findings underscore the detrimental cardiovascular effects of chronic noise exposure, highlighting the need for strategies to mitigate its impact on heart health.

### Modulation of Inflammatory Cytokine Expression in Response to Aircraft Noise Exposure

The significant up-regulation of these pro-inflammatory cytokines and related markers in individuals exposed to noise underscores the inflammatory impact of chronic noise exposure. The elevated levels of TNF-α, IFN-γ, IL-6, IL-1α, IL-1β, homocysteine, MIP-1β, MIP-1α, IL-10, and IL-4 highlight a complex inflammatory response, involving both pro-inflammatory and regulatory mechanisms. This chronic inflammation can lead to various health issues, including cardiovascular problems and cognitive impairments.

Chronic noise exposure can trigger persistent inflammation, leading to increase from mild, moderate and severe TNF-α –5.66 (–7.37 –3.95) p<0.001, –2.02 (–3.79 –0.252) p=0.02 –3.02 (–4.79 –1.25) p<0.001 production as part of the body’s defense mechanism against perceived threats (Fig. 4a). Elevated IFN-γ –7.39 (–10.6 –4.20) p<0.001, –4.06 (–7.59 –0.530) p=0.02, –4.81 (–8.34 –1.28) p=0.004(Fig. 4b) and IL-6 levels –3.84 (–6.19 –1.49) p<0.001, –3.76 (–6.56 –0.969) p=0.005 –2.92 (–5.71 –0.124) p=0.04 (Fig. 4c). Elevated IL-1α –2.63 (–3.46 –1.80) p<0.001, –1.12(–1.98 –0.264) p=0.006, –0.955 (–1.81 –0.0986) p=0.02 (Fig. 4d and IL-1β levels –2.03 (–3.29 –0.771) p<0.001, – 1.73 (–3.30 –0.152) p=0.03 –1.89 (–3.34 –0.444) p=0.006 due to noise exposure suggest that noise acts as a stressor, stimulating inflammatory pathways and exacerbating inflammatory responses (Fig. 4e). Higher homocysteine levels mild –1.31 (–2.46 –0.162) p=0.02 –1.58 (–2.86 –0.301) p=0.01, –1.68 (–3.06 –0.291), p=0.01 due to noise exposure may reflect increased oxidative stress and inflammation, which can negatively impact vascular health (Fig. 4f). Elevated levels of MIP-1α –32.9 (–43.2 –22.5) p<0.001, –16.1 (–27.1 –5.11) p=0.002, –13.7 (–24.7 –2.73) p=0.010 (Fig. 4g) and MIP-1β –31.9 (–43.4 –20.5) p<0.001, –16.6 (–28.3 –4.92) p=0.003 –18.1 (–29.8 –6.40) p<0.001 (Fig. 4h) suggest that noise exposure enhances the recruitment and activation of immune cells, contributing to the inflammatory response.

**Fig. 4.**
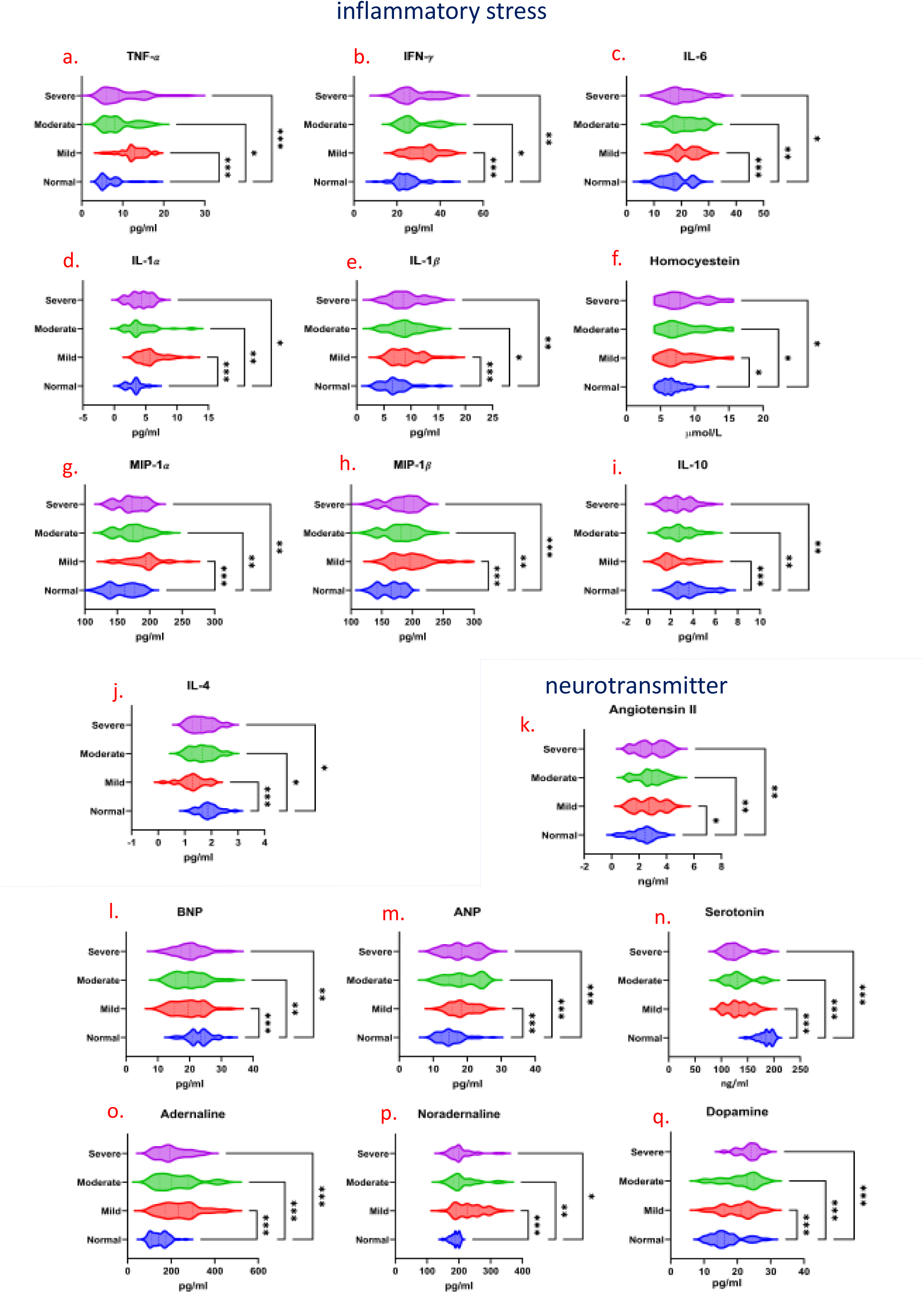
inflammatory and neurotransmitter Parameters Across Hearing Impairment Groups. inflammatory stress parameters: (a) TNF-α levels, (b) IFN-levels, (c) IL-6levels, (d) IL-1 α levels, (e) IL-1 β levels, (f) Homocysteine levels, (g) MIP-1 α levels, (h) MIP-1 β levels, (i) IL-10 levels and (j) IL-4 levels, (a-h) shown as mean ± SD, with elevated levels observed across mild to severe hearing impairment groups, (I, j) data presented as mean ± SD, with reduced levels observed in the mild to severe hearing impairment groups. **Neurotransmitter parameters**: (k) Angiotensisn-II (l) BNP (m) ANP levels, (n) Serotonin levels, (o) Adernaline levels, (p) Nordernaline level, (q) Dopamine. All data represented as mean ± SD with. statistical significance assessed using one-way ANOVA followed by Dunnett’s post-hoc test for comparisons against the normal hearing group. Where (k, m, o, p, q) showing increased levels from mild to severe hearing impairment groups, (l, n) reduced levels from mild to severe hearing impairment groups. Statistical significance is indicated where *p < 0.05, ** p<0.01 and p<0.001.

Chronic noise exposure leads to a pronounced inflammatory response, as evidenced by significantly elevated levels of pro-inflammatory markers such as TNF-α, IL-1α, IL-1β, IL-6, MIP-1β, and MIP-1α. TNF-α and IL-1β are pivotal in driving systemic inflammation, while IL-6 and MIP-1β/α contribute to the recruitment and activation of immune cells, exacerbating the inflammatory process. The increase in homocysteine further underscores the inflammatory and oxidative stress induced by noise. Conversely, there is a notable decrease in mild to severe anti-inflammatory cytokines IL-4 0.599 (0.404 0.793) p<0.001, 0.237 (0.0158 0.458) p=0.03, 0.239 (0.0173 0.460) p=0.03 (Fig. 4i) and IL-10 1.28 (0.725 1.83) p<0.001, 0.845 (0.225 1.47) p=0.004, 0.865 (0.245 1.49) p=0.003 (Fig. 4j). IL-10, which normally helps to dampen inflammation, and IL-4, involved in promoting anti-inflammatory responses, are reduced, indicating an impaired regulatory mechanism that fails to counterbalance the heightened inflammatory state induced by chronic noise exposure. This imbalance between pro-inflammatory and anti-inflammatory cytokines highlights the disruptive impact of persistent noise on the body’s ability to manage and resolve inflammation.

### Modulation of Neurotransmitter Expression in Response to Aircraft Noise Exposure

Following exposure to aircraft noise, we observed a notable decrease in the levels of several key neurotransmitters and hormones, including Brain Natriuretic Peptide (BNP), serotonin. BNP, which is involved in regulating blood pressure and fluid balance, showed reduced levels mild 3.56 (1.52 to 5.61) p<0.001, moderate 3.22 (0.908 5.52) p=0.003, severe 2.89 (0.579 5.19) p=0.009, potentially reflecting disrupted cardiovascular and stress responses (Fig. 4l). Similarly, decreased serotonin 49.4 (40.2 58.6) p<0.001, 48.7 (35.4 61.9) <0.001, 55.6 (42.3 68.9) <0.001 (Fig. 4n) suggest a negative impact on mood regulation and cognitive function, as these neurotransmitters are crucial for emotional stability and cognitive processes.

Ang II, a key regulator of blood pressure and fluid balance, also exhibited increase levels mild 0.481 (0.0647 0.898) p=0.02, moderate 0.656 (0.0792 0.872) p=0.03, severe 0.758 (0.341 1.17) p<0.001 indicating potential alterations in the renin-angiotensin system due to chronic noise stress (Fig. 4k). In contrast, we found increased levels of Atrial Natriuretic Peptide (ANP), adrenaline, and noradrenaline. ANP, which is involved in reducing blood volume and pressure, was elevated mild –3.57 (–5.48 to –1.67) p<0.001, moderate –3.71 (–5.71 to –1.70) p<0.001, severe –3.52 (–5.51 to –1.53) p<0.001 (Fig. 4m), possibly as a compensatory response to disrupted cardiovascular homeostasis. The rise in adrenaline –94.8 (–126 –63.7) p<0.001, –74.0 (–109 –38.5 p<0.001, –57.0 (–92.5 –21.5) p<0.001 (Fig. 4o) and noradrenaline mild –46.7 (–62.3 –31.1) p<0.001, moderate –25.0 (–42.7 –7.26) p=0.003, severe –19.3 (–37.0 –1.62) p=0.03 (Fig. 4p), critical for the body’s fight-or-flight response, dopamine levels 4.12 (2.02 6.23) <0.001, 2.88 (0.713 5.05) p=0.005, 3.16 (0.995 5.33) p=0.002 (Fig. 4q) suggests heightened sympathetic nervous system activity and stress response due to persistent noise exposure. These changes in neurotransmitter and hormone levels underscore the physiological impact of chronic noise exposure on both the central nervous system and cardiovascular health.

### Correlation Analysis Using Spearman’s Rank Correlation Coefficient

Reduced Glutathione negatively correlated with MDA, CRP, and 4HNE, suggesting that decreased antioxidant levels are associated with higher oxidative stress and inflammation. SOD Negatively correlated with MDA and CRP, implying that lower SOD activity is linked with higher oxidative stress and inflammation. CRP: Strongly positively correlated with TNF-α, IL-6, IL-1β, and MIP-1β, reflecting systemic inflammation. TNF-α positively correlated with IL-6 and IL-1β, indicating a cohesive inflammatory response. TNF-α, IL-1α, IL-1β, IFN-γ, MIP-1α showed significant positive correlations with adrenaline and noradrenaline (p<0.01), indicating that higher levels of these inflammatory markers are associated with increased sympathetic activity. MIP-1β and MIP-1α exhibited positive correlations with CRP, homocysteine (p<0.01), reflecting an association between heightened immune cell recruitment and increased oxidative stress. Homocysteine positively correlated with CRP and TNF-α, showing its role in systemic inflammation and oxidative stress. LDL (Low-Density Lipoprotein): Positively correlated with CRP, indicating that elevated LDL levels are associated with increased systemic inflammation. HDL negatively correlated with LDL, suggesting that lower HDL levels may be related to higher LDL and overall cardiovascular risk. CK-MB positively correlated with CRP and LDL, reflecting myocardial injury and its association with systemic inflammation. Serotonin negatively correlated with CRP and MDA, indicating that lower serotonin levels are associated with higher oxidative stress and inflammation. Dopamine negatively correlated with CRP and 4HNE, suggesting decreased dopamine levels are linked with increased oxidative stress. Angiotensin II positively correlated with CRP and LDL, indicating its involvement in cardiovascular risk and systemic inflammation. ANP negatively correlated with LDL and CRP, suggesting its role in cardiovascular regulation and response to inflammation. Glucose generally weakly correlated with most parameters, indicating that its role might be less directly related to the oxidative and inflammatory markers measured. Vitamin B12 negatively correlated with CRP and 4HNE, suggesting that lower Vitamin B12 levels are associated with higher oxidative stress and inflammation. These correlations underscore the complex interactions between oxidative stress, inflammation, cardiovascular health, and neurotransmitter levels in the context of chronic noise exposure (Fig. 5).

**Fig. 5.**
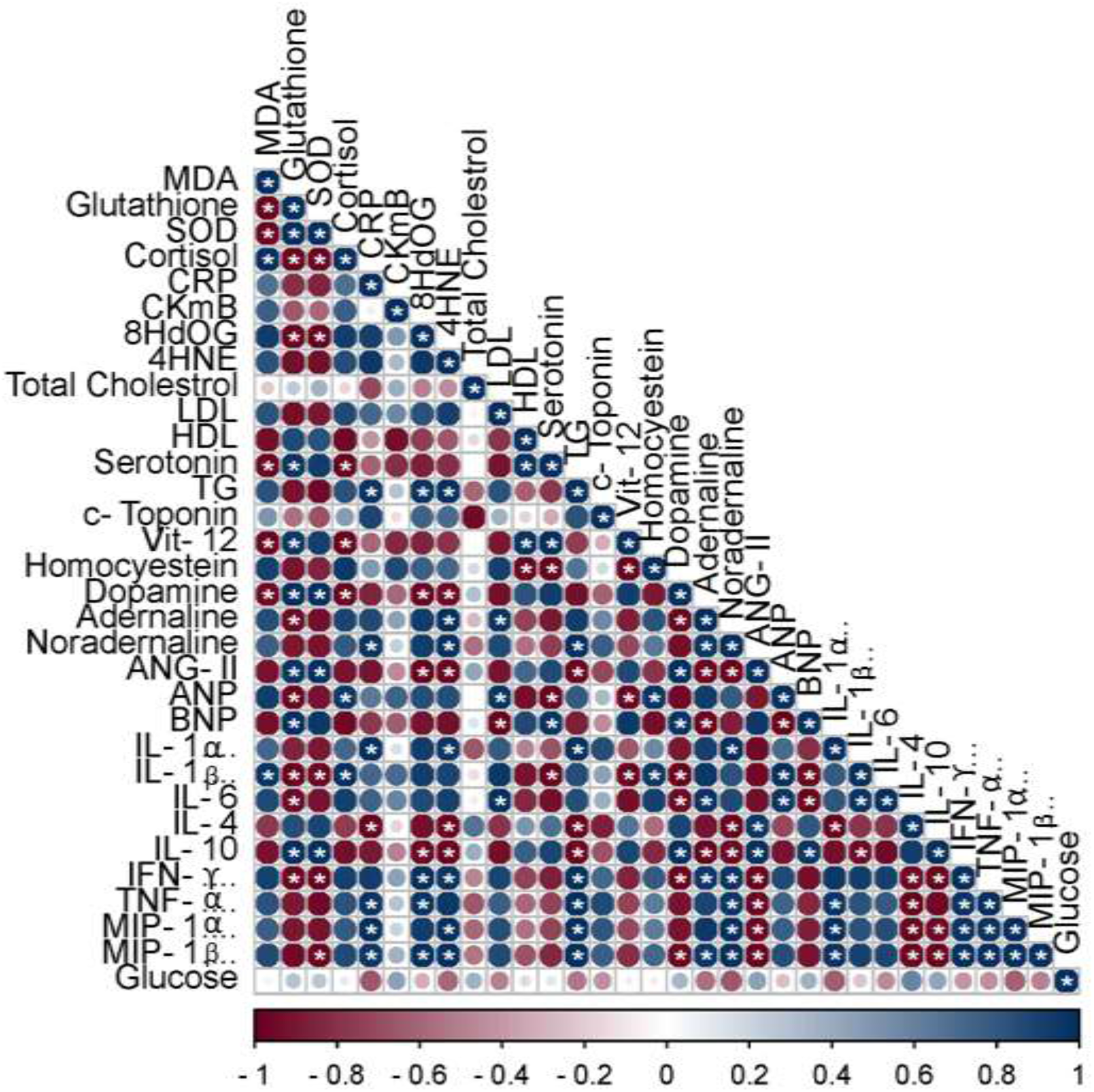
Pearson Correlation Analysis of Key Variables. Statistical significance is indicated with asterisks, where p < 0.05 is considered significant (insig p without *, sig p with *). Correlation strengths are categorized as follows: 0.1 ≤ |r| < 0.3 (weak), 0.3 ≤ |r| < 0.5 (moderate), and |r| ≥ 0.5 (strong).

### Quantitative human proteome analysis using iTRAQ Label

Our investigation began with the iTRAQ labeled; LC-MS/MS analysed human plasma proteome. An exhaustive list of proteins with quantification was obtained from LC-MS/MS to understand the casual events occurring due to noise exposure. In total 176 proteins were identified, out of which 157 proteins were differentially expressed in the experimental/hearing impaired groups when compared with normal hearing group (Additional Table 1). A total of 176 proteins were identified in the study, of which 157 were differentially expressed. Specifically, 123 differentially expressed proteins were identified in the mild hearing impairment group, comprising 67 up-regulated and 14 down-regulated proteins. In the moderate hearing impairment group, 142 differentially expressed proteins were found, including 38 up-regulated and 25 down-regulated proteins. The severe hearing impairment group also exhibited 142 differentially expressed proteins, with 63 up-regulated and 6 down-regulated proteins. Notably, a Venn diagram (Fig. 6c) revealed that 109 proteins were common across all three hearing-impaired groups, as also illustrated in the heat map (Fig. 6d).

**Fig. 6.**
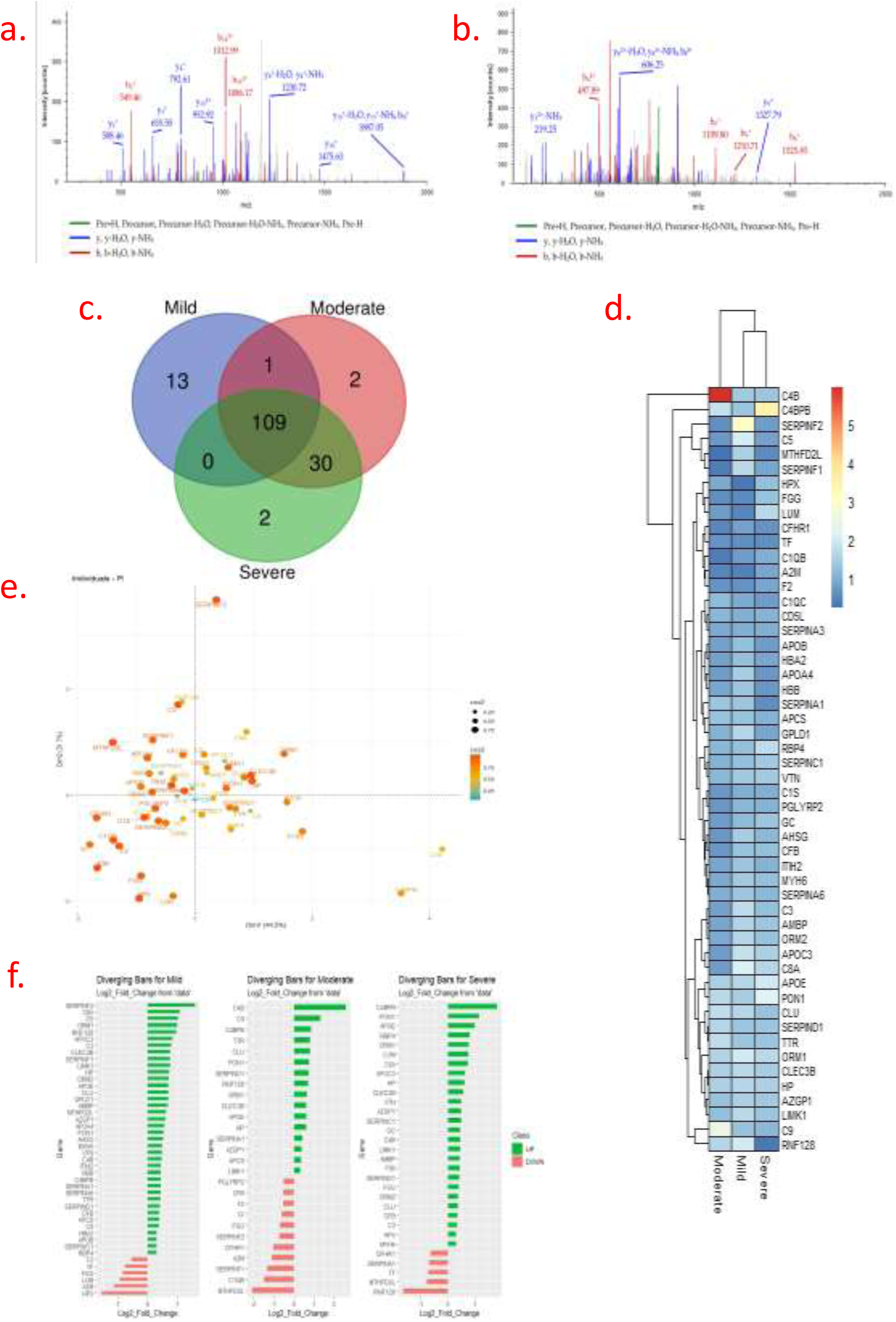
Proteomic Analysis and Protein Expression Across Hearing Impairment Groups. (a) Representative MS/MS spectra showing two unique peptides identified from serotransferrin (EDLIWELLNQAQEHFGK) (b) Representative MS/MS spectra showing two unique peptides identified from transthyretin (DLIWELLNQAQEHFGK). (c) Venn diagram illustrating the overlap of proteins identified and differentially expressed across hearing impairment groups: 157 out of 176 totals differentially expressed proteins identified in all impairment. 123 differentially expressed proteins in the mild group. 142 differentially expressed proteins in the moderate group. 157 differentially expressed proteins in the severe group. 109 proteins common to all groups. (d) Heat map depicting the expression levels of differentially expressed proteins across mild, moderate, and severe hearing impairment groups. Expression levels are shown as log2 fold changes, with color gradients indicating relative abundance. (e) Principal Component Analysis (PCA) plot showing the distribution of differentially expressed proteins across mild, moderate, and severe hearing impairment groups. The plot illustrates the variance and clustering of protein expression profiles. (f) Diverging bar chart representing the log2 fold changes of differentially expressed proteins in the mild, moderate, and severe groups. The bars indicate the magnitude and direction of change in protein expression relative to the normal hearing group. Data are presented as mean ± SD, with statistical significance evaluated using appropriate methods as specified in the methods section.

The principal component analysis (PCA) is a statistical tool for sample classification based on multivariate data. Each point in the PCA graph represents the whole protein profile of one biological sample. Samples with similar behavior in their protein profile are grouped together. In our work the PCA graph for iTRAQ data showed a clear separation between mild, moderate, and severe hearing-impaired group samples (Fig. 6e).

The protein symbol corresponding to their protein accession number was considered for network construction, although some of the proteins were omitted from the study due to their unknown characteristics. The node size and edge thickness represent fold change value and interaction score (experimental only) obtained from STRINGdb. White, pink and green colors indicate proteins with fold change values >0.8 to <1.2 (insignificant), >1.2 (Up-regulated) and <0.8 (down-regulated) respectively in control, moderate and severe hearing-impaired groups (left to right) (Fig. 6f)

### Functional enrichment analysis of DEGs after sample filtration

An examination of 157 dysregulated proteins revealed significant findings through Gene Ontology Biological Process (GO-BP) terms. Functional and pathway enrichment analyses using the DAVID tool identified several noteworthy results. The GO KEGG analysis highlighted that the most significantly enriched modules were related to the complement and coagulation system, coronavirus disease (COVID-19), Staphylococcus aureus infection, systemic lupus erythematosus (SLE), extracellular matrix (ECM) structural constituents, and cholesterol metabolism in the hearing-impaired groups.

In terms of biological processes (BPs), the mild hearing impairment group showed significant enrichment in the humoral immune response, acute inflammatory response, blood coagulation, hemostasis, acute phase response, complement activation, humoral response mediated by circulating immunoglobulins, and complement cascade classical pathway (Fig. 7Ai, 7D). The moderate group exhibited enrichment in the humoral immune response, complement activation, wound healing, acute phase response, acute inflammatory response, blood coagulation, coagulation, and hemostasis (Fig. 7b.i, 7e). The severe group showed similar enrichment with additional involvement in the B cell-mediated immune response (Fig. 7c.i, 7f).

**Fig. 7.**
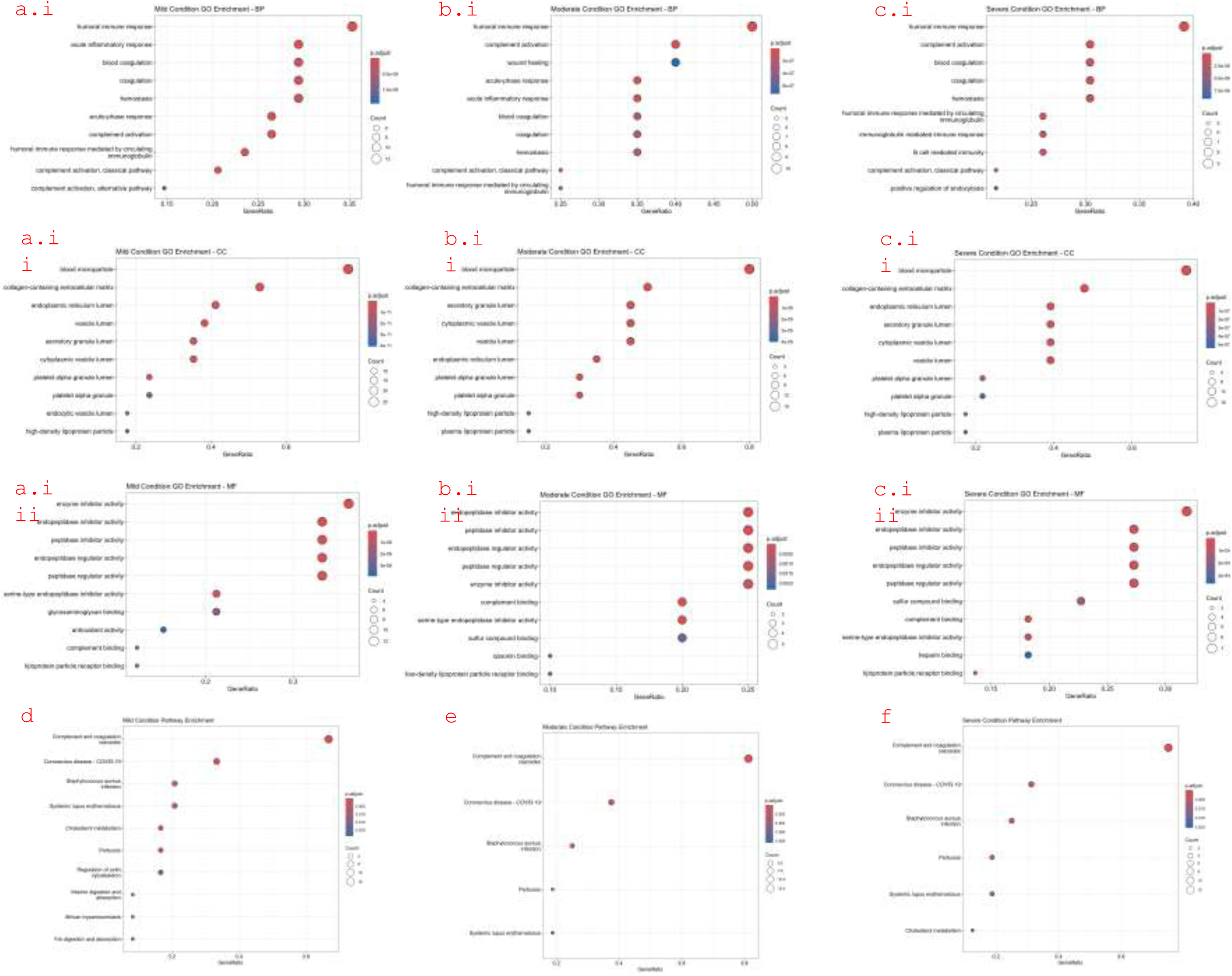
GO-KEGG Pathway Analysis of Hearing Impairment Severity Panels a, b, and c present the results of Gene Ontology (GO) term enrichment analysis for different severities of hearing impairment. Panel a illustrates the findings for the mild hearing loss group, showing Biological Processes (BP) in a.i, Cellular Components (CC) in a.ii, and Molecular Functions (MF) in a.iii. Panel b depicts the moderate hearing loss group, with BP in b.i, CC in b.ii, and MF in b.iii. Panel c details the severe hearing loss group, showing BP in c.i, CC in c.ii, and MF in c.iii. Panels d, e, and f provide insights into the KEGG pathway analysis for each severity level. d highlights the KEGG pathways enriched in the mild hearing loss group, e displays the pathways for the moderate hearing loss group, and f outlines the pathways for the severe hearing loss group. These panels collectively illustrate the biological processes, cellular components, and molecular functions associated with different levels of hearing impairment, alongside the key metabolic and signaling pathways affected at each severity level.

Regarding cellular components (CCs), dysregulated proteins were predominantly enriched in blood microparticles, collagen-containing extracellular matrix, endoplasmic reticulum lumen, vesicle lumen, secretory vesicle lumen, cytoplasmic vesicle lumen, and platelet alpha granules across all hearing impairment levels (Fig. 7a.ii, b.ii, c.ii).

GO analysis of dysregulated proteins indicated distinct molecular functions (MF) associated with different levels of hearing impairment. The mild group (Fig. 7a.iii) exhibited significant changes in enzyme inhibitor activity, endopeptidase inhibitor activity, peptidase inhibitor activity, endopeptidase regulator activity, and peptidase regulator activity. The moderate group (Fig. 7b.iii) showed alterations in enzyme inhibitor activity, endopeptidase inhibitor activity, peptidase inhibitor activity, endopeptidase regulator activity, peptidase regulator activity, complement binding, serine-type endopeptidase inhibitor activity, and sulfur compound binding. The severe group (Fig. 7c.iii) highlighted changes in enzyme inhibitor activity, endopeptidase inhibitor activity, peptidase inhibitor activity, endopeptidase regulator activity, peptidase regulator activity, complement binding, serine-type endopeptidase inhibitor activity, sulfur compound binding, and heparin binding.

Furthermore, seven differentially expressed proteins were implicated in the complement and coagulation cascade pathway. These results suggest that proteins significantly affected by noise exposure are primarily associated with stress, coagulation, and inflammatory pathways. Based on interaction and involvement in these processes, common 52 different proteins were selected for further analysis.

## Establishment of the PPI network

The StringDB constructed the predicted protein-protein associations based on the following information such as experimental validation, co-expression, coregulation and shared domains [48]. The DEGs were mapped using STRING to evaluate the connections between genes, with a confidence score threshold set at > 0.4 to determine significance. Cytoscape, a software for network biology analysis and visualization [46], was employed to generate protein-protein interaction (PPI) networks. Utilizing the STRING database and Cytoscape software, a protein-protein interaction (PPI) network was constructed (Fig. 8a, 8b, 8c). 52 protein from each group mild moderate and severe were identified from the PPI network using the CytoHubba plugin and the maximal clique centrality method. Data mining revealed that both the significant module and the hub genes predominantly consisted of upregulated protein. The protein-protein weighted graph network assigns a topological values foe each note in the network. These topological parameters are critical for determining important proteins and information flow networks. The PPI network information obtained from StringDB was used to construct a weighted network. CytoHubba, a cytoscape plugin analyzed the weighted PPI network Based on the topologies of the network. We extracted the 52 proteins corresponding to each topology mentioned in the Fig. mild 8a, moderate 8b, sever 8c. Betweenness, Bottleneck, Closeness, Clustering Coefficient, Degree, Density of Maximum Neighborhood Component (DMNC), EcCentricity, Edge Percolated Component (EPC), Maximal Clique Centrality (MCC), MNC, Radiality and Stress are classified as local properties of a nodes based on direct connection to its neighborhood nodes. These topological values don’t affect the network but restrict to the node specific itself. For example, degree is simply the number of total edges connected to a particular node. While the global based nodes impact the whole network such as bottleneck, betweenness, and closeness etc. CLU, C5, HP, C3, C4B, TTR are the top bottleneck proteins indicating the most trafficking point in the network in mild to severe group. Betweenness graph is showing that C4B, C9, HP and F2 are the proteins shares maximum pathways in a network while walking over the whole network. In a similar way, C5, APOE, CLU, C3, APOB, SERPINA1, ORM2 showed a measure of minimum distance against all the nodes in respective network termed as closeness topology.

**Fig. 8.**
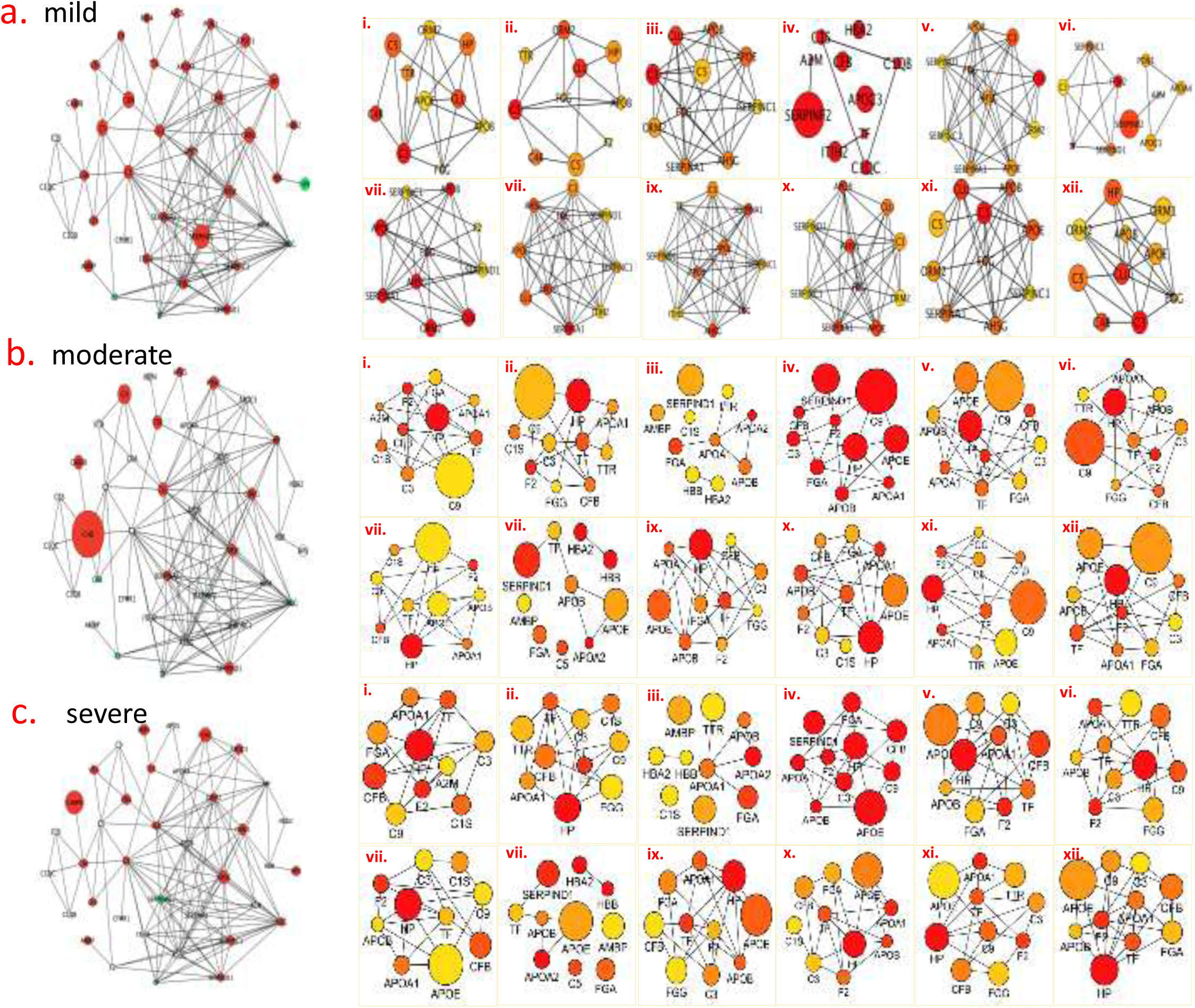
Protein-Protein Interaction (PPI) Network Analysis Based on Top 52 Proteins with High and Low Expression Levels. PPI network representation showing interactions among the top 52 proteins with the highest and lowest expression levels across (a) mild, (b) moderate, (csevere) hearing impairment groups. The network visualizes the relationships and interactions between these proteins. Where in each group **(i) Betweenness Centrality:** Indicates the extent to which a protein lies on the shortest path between other proteins, highlighting key intermediaries in the network. **(ii) Bottleneck Centrality:** Measures the frequency with which a protein acts as a bottleneck in the network, indicating its role in network flow. **(iii) Closeness Centrality:** Reflects how close a protein is to all other proteins in the network, representing its ability to access other proteins quickly. **(iv) Clustering Coefficient:** Shows the degree to which proteins cluster together, indicating the local density of connections. **(v) Degree Centrality:** Represents the number of direct connections a protein has, illustrating its overall connectivity within the network. **(vi) Degree of Minimum Node Centrality (DMNC):** Assesses the minimal number of connections required for a protein to maintain network integrity **(vii) Eccentricity:** Measures the maximum distance from a protein to any other protein in the network, indicating its centrality. **(viii) Eigenvector Centrality (EPC):** Indicates a protein’s influence based on its connections and the influence of its neighbors. **(ix) Maximal Clique Centrality (MCC):** Identifies proteins involved in the largest cliques, or fully connected sub-networks. **(x) Minimum Node Centrality (MNC):** Reflects the minimum number of connections necessary for a protein’s involvement in the network. **(xi) Radiality:** Measures how central a protein is in terms of network topology and shortest paths. **(xii) Stress Centrality:** Assesses the extent to which a protein is involved in the network’s shortest paths, indicating its role in network stability. The network visualizations and metrics provide insights into the functional significance and interconnectivity of the top 52 proteins with varying expression levels.

### Modulation of proteins activating responses to stress, hearing, cognition, coagulation and inflammatory pathways

Twelve distinct proteins have been identified as significantly associated with the activation of stress responses, coagulation, and inflammatory pathways. Vitronectin (VTN) and serum amyloid protein A (APCS) are linked with coagulation, stress, and inflammation. Complement component 5 (C5), hemopexin (HPX), and alpha-1-acid glycoprotein-1a (ORM1) are associated with stress and inflammatory responses. Retinol binding protein 4 (RBP4) and alpha-2-microglobulin (A2M) are related to inflammation and coagulation, respectively. Among these proteins, HPX, APCS, ORM1, and VTN show increased expression, while A2M shows decreased expression. C5a is elevated in cases of mild hearing loss but decreases in moderate and severe cases. RBP4 consistently increases with greater hearing loss severity. Additionally, Transthyretin (TTR), Isoform 4 of Clusterin (CLU), and Tetranectin (CLEC3B) are significantly upregulated with worsening hearing loss, whereas Sero-transferrin (TF) is downregulated. LIM Kinase 1 (LIMK1) is a serine/threonine kinase crucial for regulating actin cytoskeleton dynamics, which affects cell movement, shape, and differentiation. Transthyretin (TTR) is involved in thyroid hormone and retinol transport and can accumulate abnormally in conditions such as amyloidosis. While TTR’s direct role in hearing loss is less clear, it may influence hearing through systemic and local metabolic processes. C-Type Lectin Domain Family 3 (CLEC3), also known as Tetranectin, plays a role in inflammation and tissue remodeling. Its upregulation in the context of hearing loss indicates it may contribute to the pathophysiology of hearing impairments.

### Validation of plasma proteomic differential levels between normal and impaired group sample by Elisa and western blot

For validation, ten proteins that showed dys-regulation in at least one of the impaired group samples and belonging to different pathways were selected for western blotting (Fig. 9a). The results are in accordance with the proteomic analysis. To investigate the plausible effect of noise exposure on the expression of these ten selected proteins, plasma level of protein was measured. Though the concentration and expression pattern varied, a significant change in the plasma concentration was observed in all the proteins. Fig. 9 presents the mean values for the plasma levels of these proteins at each study time point.

**Fig. 9.**
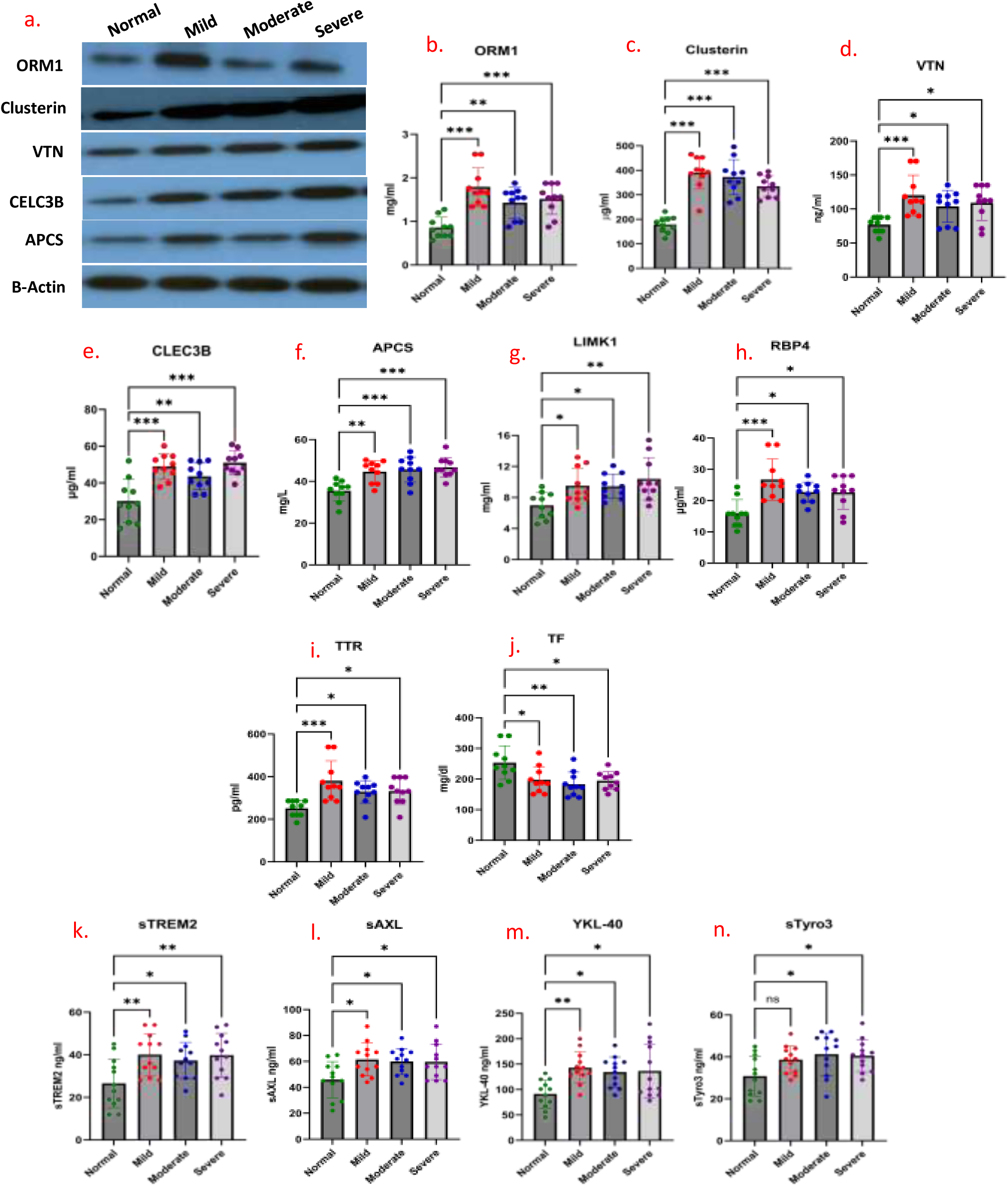
Western Blot and ELISA Validation of Protein Dysregulation Across Hearing Impairment Groups. (a) Western blot results showing the expression of the following proteins: ORM1 (Orosomucoid 1), Clusterin, VTN (Vitronectin), CLEC3B (C-Type Lectin Domain Family 3 Member B), APCS (Apolipoprotein C1) Each protein is shown with its respective bands, indicating expression levels across different hearing impairment groups. (b) – (j) ELISA validation of protein dysregulation across hearing impairment groups: (b) ORM levels in mild, moderate, and severe hearing impairment groups compared to normal hearing group. (c) Clusterin (d) VTN levels (e) CLEC3B levels (f) APCS levels (g) LIMK1 (LIM Domain Kinase 1), (h) RBP4 (Retinol-Binding Protein 4) levels, (i) TTR (Transthyretin), (j) TF (Transferrin) levels in mild, moderate, and severe hearing impairment groups. (k-n) Validation of inflammatory serum panel: sTREM2, sAXL, YKL-40, sTyro3. Data for ELISA are presented as mean ± SD, with statistical significance determined using one way. ELISA results are compared between mild, moderate, and severe groups, highlighting the dysregulation of proteins identified in the iTRAQ analysis. All measured unit mentioned in each ELISA Y-axis. Statistical significance was assessed using one-way ANOVA with post-hoc Dunnett’s test. Statistical significance is indicated where *p < 0.05, ** p<0.01 and p<0.001.

A significant change (p<0.001) in the plasma concentration was obtained for all protein molecules. However, the expression pattern of each protein in response to noise exposure varied. Plasma levels of ORM1 –0.933 (CI 95%-1.32 –0.546) p<0.001, –0.570 (CI 95% –0.957 –0.183) p=0.003, –0.655 (CI 95%-1.04 to –0.268) <0.001 (Fig. 9b); CLU –212 (–273 to –150) p<0.001, – 194 (–255 to –132) p<0.001, –156 (–217 to –94.4) p<0.001 (Fig. 9c); VTN –43.1 (–68.9 to –17.3) p<0.001, –26.9 (–52.8 to –1.12) p=0.04 –32.1 (–57.9 to –6.28) p=0.01 (Fig. 9d); CLEC3-18.8 (–28.3 –9.42) p<0.001, –13.4 (–22.8 –3.98) p=0.003, –20.8 (–30.2 –11.4) p<0.001 (Fig. 9e); APCS, –9.17 (–14.9 –3.42) p=0.001, –10.1 (–15.9 –4.37) p<0.001, –11.2 (–17.0 –5.47) p<0.001 (Fig. 9f); LIMK1 –2.53 (–4.87 to –0.190) p=0.03, –2.44 (–4.78 to –0.100) p=0.04, –3.40 (–5.74 to –1.06) p=0.003 (Fig. 9g); Rbp4 –10.9 (–16.5 to –5.22) p<0.001, –6.84 (–12.5 to –1.20) p=0.01, –6.72 (–12.4 to –1.08) p=0.01 (Fig. 9h); TTR –129 (–200 to –58.5) p<0.001, –78.1 (–149 to –7.53) p=0.03, –80.2 (–151 to –9.66) p=0.02 (Fig. 9i); and increased with increasing severity of hearing loss, while TF 56.0 (7.76 104) p=0.02, 70.3 (22.1 119) p=0.002, 58.7 (10.5 to 107) p=0.01 was downregulated.

## Discussion

The present study was conducted on military personnel exposed to aircraft noise. Ground duty personnel concerned with the maintenance and operation of high-performance aircrafts served as study sample. Consequent on their auditory evaluation for assessment of hearing status through pure tone audiometry in the frequency range of 125 Hz to 8 kHz for both ears, they were categorized as those having normal hearing with Hearing level up to 25 dB, mild hearing impaired with Hearing level between 25-40dB, moderate hearing impaired with Hearing level between 40-60dB. Personnel with Hearing levels higher than 60dB were adjudged as severe hearing impaired. In the distinct analysis of frequencies, significant differences in PTA 2000, 3000, 4000 and the most significantly difference at 6000 Hz was observed in both ears in all the groups. There were significant differences between the two groups viz. normal and with impaired hearing in relation to threshold changes suggesting NIHL and with Right ear more affected than the left ear. The notch/dip in hearing threshold at higher frequencies has long been recognized as a clinical sign of exposure to noise, and although the classic association is between continuous exposure to noise and a notch at 4 kHz, notches have been also observed at 6 kHz in people/individuals exposed to high frequency noise and at 3 kHz with low frequency noise. Audiometric variability is greater at 6 than at 4 kHz.

ABR testing revealed significant changes in wave latencies and amplitudes, correlating with the severity of hearing impairment our previous study also showed similarly result in rat [8]. These findings suggest that ABR can detect functional deficits and cochlear damage that are not always evident through standard audiometric thresholds. DPOAE testing demonstrated a reduction in outer hair cell (OHC) function in individuals exposed to aircraft noise, aligning with the findings from audiometry and ABR tests. This reduction in DPOAE amplitudes confirms the impact of noise on cochlear function and supports the use of DPOAE as a valuable tool for detecting early OHC damage and assessing auditory health our previous study also showed similarly result in armed forced exposed to intense noise during training [16].

Current study focused on changes in expression of plasma proteome with the increase in noise induced hearing impairment. The expression was comparatively evaluated between normal subjects and different hearing-impaired groups, namely mild, moderate and severely hearing impaired. iTRAQ-LCMS/MS was performed to determine the proteins with differential expression exhibiting plausible association with the manifestation of hearing impairment in noisy environment. The experiment was conducted simultaneously on individual samples and pooled samples as well. Further, the relative expression of proteins/peptides was compared between impaired and normal groups in all the samples along with protein-protein interaction and ontology. Proteins involved in hearing, coagulation and inflammation with response towards noise were chosen for further evaluation. Results of the current investigations were suggestive of hearing loss inducing oxidative stress leading to inflammation and altered plasma proteome. The pathophysiological basis of NIHL has been extensively studied; factors leading to hearing loss after noise exposure include direct mechanical trauma, oxidative stress, metabolic exhaustion, ischemia and ionic imbalance in the inner ear fluids. In this study, an important aspect redox homeostasis, a key process in noise exposure has been explored in plasma through estimation of oxidative stress parameters and antioxidants levels using biochemical assay in the noise induced mild, moderate and severe hearing-impaired subjects.

### Impact of Noise Exposure on Inflammation and Oxidative Stress: Analyzing Circulating Inflammatory Cytokines and Oxidative Stress Markers in Relation to Hearing Loss Severity

The influence of noise on inflammation and oxidative stress was evaluated by examining circulating levels of inflammatory cytokines and oxidative stress markers in subjects with varying degrees of hearing loss or impairment. Our analysis aimed to determine the extent to which chronic noise exposure contributes to systemic inflammation and oxidative damage, which are known to have significant implications for both auditory and overall health [49, 50], Inflammation also plays an important role in hearing disorders induced by noise, drugs, and advancing age[51]. Noise exposure can damage cochlear function by inducing inflammation in animal models[52]. This is supported by the fact that exposure to noise increased the levels of intracellular adhesion molecules and migration of leukocytes [53]. We measured circulating levels of several key inflammatory cytokines, including TNF-α, IL-6, IFN-γ, IL-4, and IL-10. Our findings indicate a marked increase in pro-inflammatory cytokines (TNF-α, IL-6, and IFN-γ) across all severity levels of hearing impairment [54, 55]. Specifically, subjects with moderate to severe hearing loss showed significantly higher levels of TNF-α and IL-6 compared to those with mild hearing loss or normal hearing. levels of GSH and SOD, which are critical antioxidants that protect cells from oxidative damage, were found to be significantly reduced in subjects with moderate to severe hearing impairment. The decrease in these antioxidants suggests a compromised ability to neutralize reactive oxygen species (ROS), further contributing to oxidative stress and cellular damage[56]. Correlation analysis revealed a positive relationship between levels of pro-inflammatory cytokines and oxidative stress markers. Specifically, higher levels of TNF-α and IL-6 were correlated with increased MDA levels, suggesting that inflammation may contribute to heightened oxidative stress. Conversely, there was a negative correlation between anti-inflammatory cytokines and oxidative stress markers, indicating that impaired anti-inflammatory responses may exacerbate oxidative damage.

Increased inflammation in the elder individuals has been associated with both age-related hearing loss [57]and non-age-related sensorineural hearing loss [58]. An epidemiologic study found that prolonged elevation of serum C-reactive protein (CRP), a marker of general inflammation, was associated with increased risk of developing hearing impairment in individuals under but not over the age of 60 years [59]. In another epidemiologic study, the levels of C-reactive protein (CRP), IL-6, and TNF-alpha have been directly correlated with hearing loss. Evidence for chronic inflammation has been found in aging cochlea [60].

### Impact of Noise Exposure on cognitive impairment

Our study investigated the cognitive impact of hearing impairment using the CANTAB battery, focusing on Spatial Working Memory (SWM), Paired Associates Learning (PAL), and Simple Reaction Time (SRT). The results provide compelling evidence that hearing impairment significantly affects cognitive functions, particularly in spatial working memory, associative learning, and simple cognitive processing. The mechanisms linking hearing loss to dementia are predominantly associated with sensorineural hearing changes, which involve dysfunction of the cochlea. This is most observed as age-related hearing loss (ARHL) or presbycusis over 65 years old [61]. Various causes of hearing loss, including presbycusis, noise-induced hearing loss (NIHL), and ototoxicity, lead to permanent hearing impairment and subsequently limit available management options [62]. Meta-analyses pooling observational studies have reinforced the link between age-related hearing loss (ARHL) and cognitive decline [63–65]. Similar correlations have also been observed in rats, supporting this association across different models [7, 8].

### Impact of Noise Exposure on autonomic regulation and cardiac

In the HRV test, we observed a significant decrease in RMSSD and high-frequency power (HF nu) across subjects exposed to aircraft noise. The reduction in RMSSD indicates diminished parasympathetic nervous system activity and vagal tone, reflecting a decreased ability to regulate heart rate variability under stress [66, 67]. Similarly, the decrease in HF nu suggests a reduction in the parasympathetic component of heart rate variability. In contrast, there was a notable increase in the LF: HF ratio and LF nu, indicating a shift towards greater sympathetic nervous system dominance relative to parasympathetic activity[68]. This shift suggests an overall increase in sympathetic activity and a reduced balance between sympathetic and parasympathetic influences, underscoring the impact of chronic noise exposure on autonomic regulation and cardiovascular health.

Epidemiological studies have linked traffic noise exposure with cardiovascular diseases, such as hypertension, myocardial infarction, and stroke [18, 69, 70]. Noise acts as a non-specific stressor, stimulating both the autonomic and endocrine systems. Chronic exposure to low levels of noise can disrupt activity, sleep, and communication, leading to emotional responses such as annoyance and subsequent stress [71] Our findings align with previous research, which highlights the adverse effects of chronic noise exposure on cardiovascular function and autonomic regulation.

### The influence of noise on circulating plasma protein

The analysis of 157 dysregulated proteins in the context of hearing impairment has provided substantial insights into the biological processes, molecular functions, and cellular components associated with various severities of auditory dysfunction. GO-BP terms, enriched through DAVID tool analyses, reveal a prominent involvement of pathways related to inflammation, coagulation, and stress responses across different levels of hearing impairment. Through gene ontology analysis, two prominent yet unrelated pathways namely, complement and coagulation cascade pathways were noticed [72]. The complex interplay of proteins associated with stress responses, coagulation, and inflammation provides insight into their roles in hearing loss. Vitronectin (VTN) and serum amyloid protein A (APCS) are integral to coagulation, stress responses, and inflammatory pathways [73, 74], with their elevated expression suggesting heightened systemic responses to these conditions. The presence of APCS in all cerebral Aβ plaques and most neurofibrillary tangles in Alzheimer’s disease effectively distinguishes AD brains from cognitively normal ones [75]. Vitronectin is a glycoprotein that is found in the extracellular matrix of various tissues, including the inner ear. one study found that exposure to loud noise increased the expression of vitronectin in the cochlea of rats [76]. Another study found that vitronectin levels were significantly higher in the blood of workers who had been exposed to occupational noise [76]. It is believed that vitronectin may play a role in the pathogenesis of NIHL by contributing to the inflammatory response that occurs following noise exposure. Specifically, it is thought that vitronectin may bind to damaged hair cells in the inner ear and activate immune cells, leading to an inflammatory response that can contribute to further damage [77].

Complement component 5 (C5), hemopexin (HPX), and alpha-1-acid glycoprotein-1a (ORM1) are particularly implicated in stress and inflammation [78]; their increased levels are consistent with an active inflammatory state or stress response. Conversely, alpha-2-microglobulin (A2M) shows decreased expression, possibly reflecting altered protease inhibition or changes in inflammation-related pathways [79]. The differential regulation of C5a and retinol binding protein 4 (RBP4) with hearing loss severity highlights their potential roles in auditory dysfunction: C5a decrease in moderate to severe hearing loss might indicate a shift in inflammatory response or complement activation, while RBP4 consistent increase with worsening hearing loss suggests its involvement in the progressive nature of hearing impairments. Additionally, transthyretin (TTR), isoform 4 of Clusterin (CLU), and Tetranectin (CLEC3B) are markedly upregulated with greater hearing loss severity. TTR, known for its role in thyroid hormone and retinol transport, might influence hearing through broader metabolic or systemic effects, though its direct role remains ambiguous [80]. Tetranectin (CLEC3B), implicated in inflammation and tissue remodeling, appears to be a key player in the pathophysiology of hearing loss, potentially contributing to the inflammatory and structural changes observed in auditory tissues. The downregulation of Sero-transferrin (TF) further supports the notion of complex alterations in iron homeostasis and oxidative stress in hearing loss. Collectively, these findings underscore the multifaceted roles of these proteins in the pathogenesis of hearing impairments, emphasizing the need for further investigation into their precise mechanisms and interactions.

TTR, known for its roles in cognition, memory, psychological health, and emotion [81, 82], is also linked to oxidative stress, a major factor in Alzheimer’s disease (AD) and other neurodegenerative disorders [83] Our findings highlight elevated TTR expression with worsening hearing loss, reflecting its primary function in transporting thyroxine and retinol bound to retinol binding protein (RBP4), and its broader biological roles related to antioxidant and oxidant properties, making TTR a potential biomarker for oxidative stress.

## Conclusion

This study assessed the impact of aircraft noise on military personnel by analyzing hearing sensitivity, inflammation, oxidative stress, and cognitive function. Audiometric evaluations categorized subjects into normal, mild, moderate, and severe hearing impairment groups, revealing significant threshold shifts, especially at high frequencies, consistent with noise-induced hearing loss (NIHL). ABR testing showed altered wave latencies and amplitudes correlating with hearing loss severity, highlighting ABR’s effectiveness in detecting cochlear damage. DPOAE testing further confirmed impaired outer hair cell function due to noise exposure. Protein analysis identified upregulation of haptoglobin, alpha-1-antiproteinase, APCS, VTN, LIMK1, transthyretin, and other proteins linked to oxidative stress, coagulation, cognitive and inflammation. The study underscores the multifaceted effects of noise exposure on auditory and systemic health. The results highlight the importance of using comprehensive diagnostic approaches, including protein biomarkers, to assess and manage noise-induced hearing damage effectively. Addressing these factors is crucial for developing preventive strategies and interventions to mitigate the adverse effects of chronic noise exposure on hearing and overall health.

## Acknowledgements

We would like to express our sincere gratitude to all the personnel from the armed forces who participated in this study. Their cooperation and dedication were instrumental in making this research possible. Our appreciation extends to supported by the Defence Research and Development Organization (Project DIP-266) for their financial support, which was crucial in conducting this research. Additionally, we acknowledge DIPAS for providing the facilities and resources necessary for this study. Finally, we wish to thank the editorial team and reviewers for their time and constructive feedback, which greatly contributed to the refinement of this manuscript.

## Funding

This work was supported by grants from the Defence Research and Development Organization (Project DIP-266), Ministry of Defence, Government of India, Delhi.

## Author information

Manish Shukla, Jai Chand Patel, Meenakshi Shukla, Shutanu Chakravarti, Neeru Kapoor Occupation Health, Defence Institute of Physiology & Allied Science (DIPAS), DRDO, Timarpur, Delhi 110054; Devasharma Nayak Proof and Experimental Establishment (PXE), DRDO, Chandipur, Balasore, Odisha 756025

## Contributions

MS conceived the study, performed experiment; MS, NK designed experiments; JCP performed bioinformatics & statistical analysis; MS, NK, DS field duty, CANTAB, HRV; MS, MS ABR, DPOAE; MS audiometry data acquired; DSN Blood sample collection, processing, ELISA performed; MS, MS performed experiments and collected and analyzed data. SC performed HRV data analysis; MS, JCP prepared Fig. composites. MS and NK wrote the manuscript. All authors read, provided feedback, and approved the final manuscript text and Figures for publication.

## Corresponding author

Correspondence to Manish Shukla, Neeru Kapoor.

## Ethics declarations

The study was approved by the Institutional Human Ethical Committee (IHEC) and adhered to the ethical standards set by the Indian Council of Medical Research (ICMR) [approval no. 01/IEC/DIPAS/14 to 20]. The participants were explained with the details of the study design and written informed consent was obtained.

## Consent for publication

Not applicable.

## Competing interests

The authors declare that they have no competing interests.

---

**Additional Table 1.**
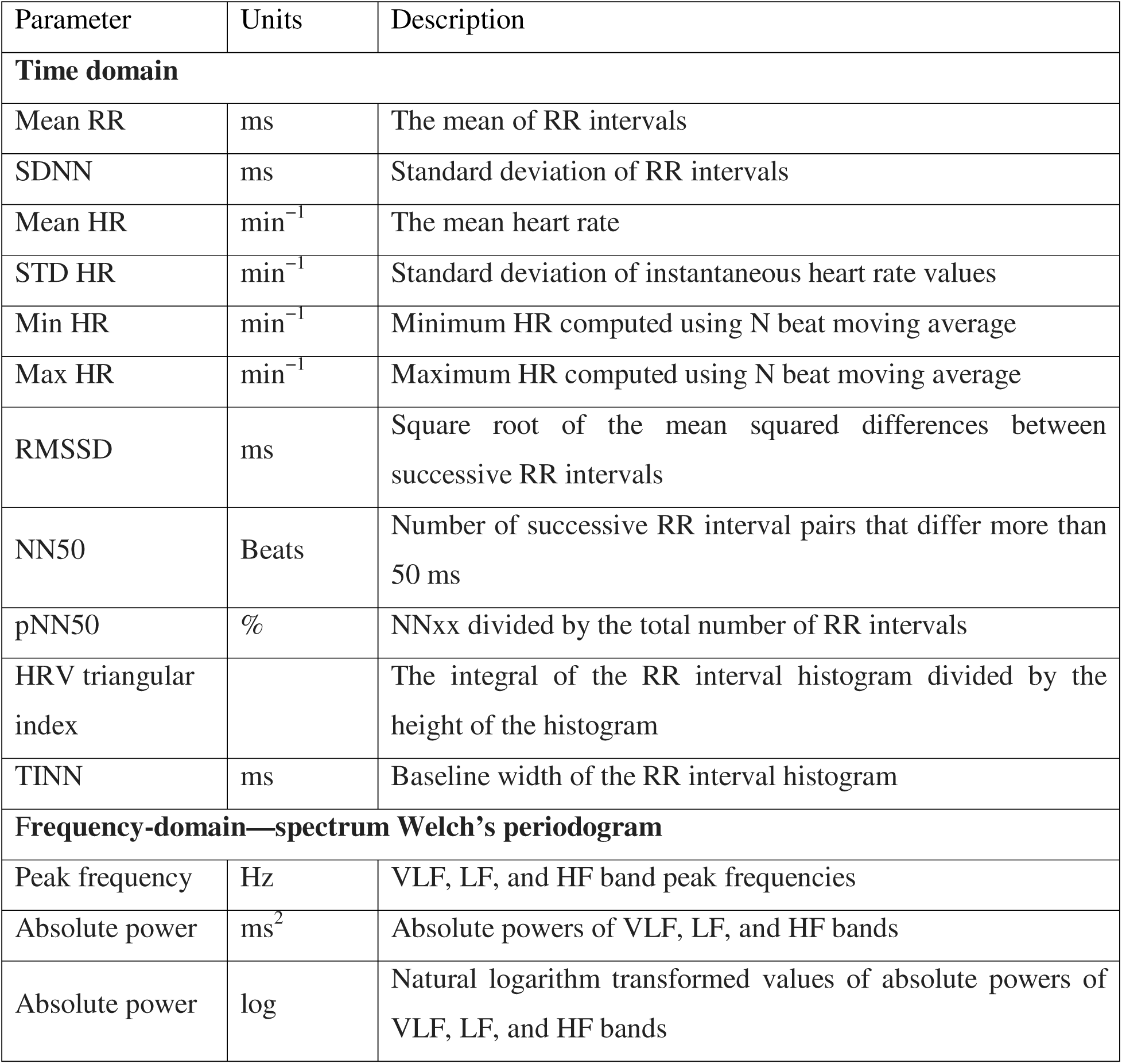
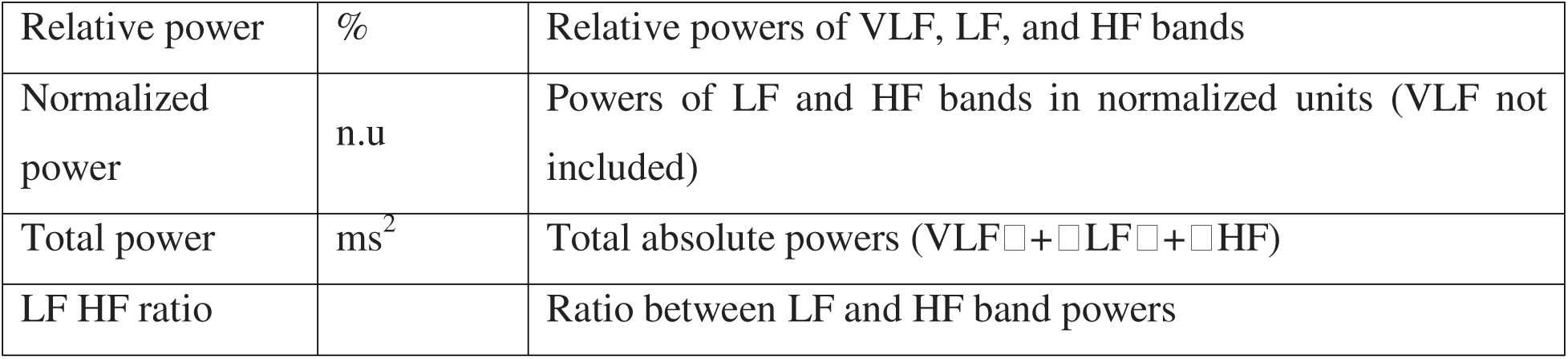
Summary of executing HRV parameters.

**Additional Table 2:**
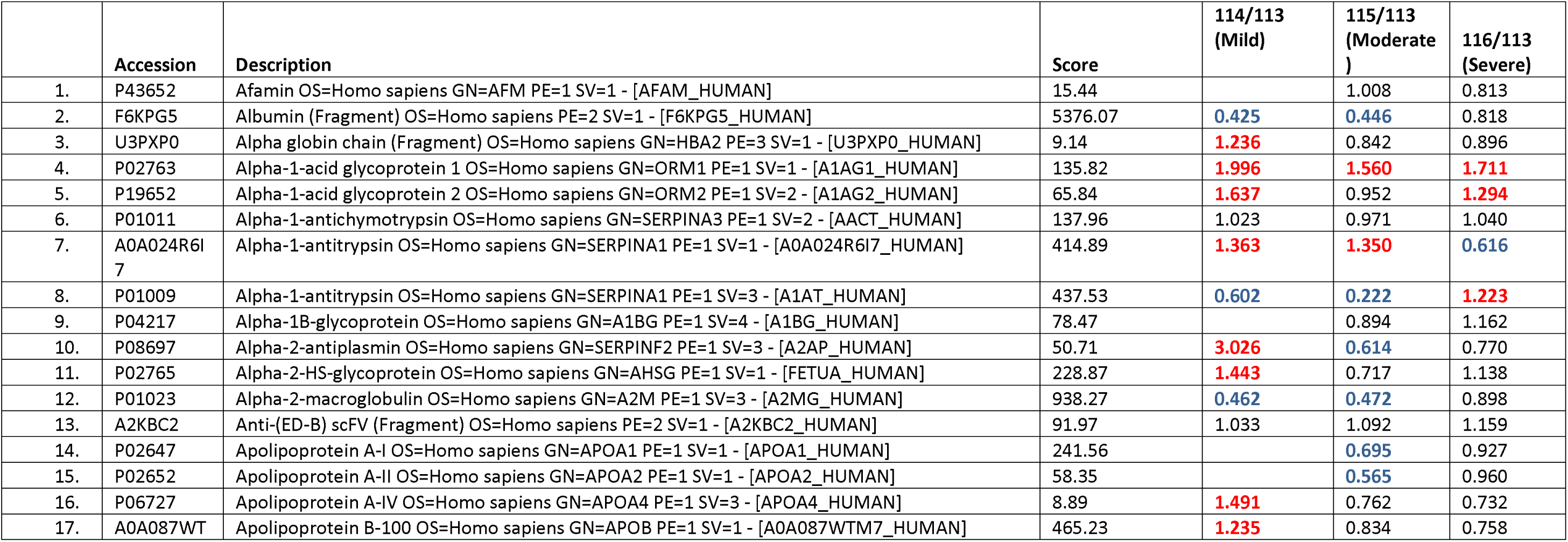
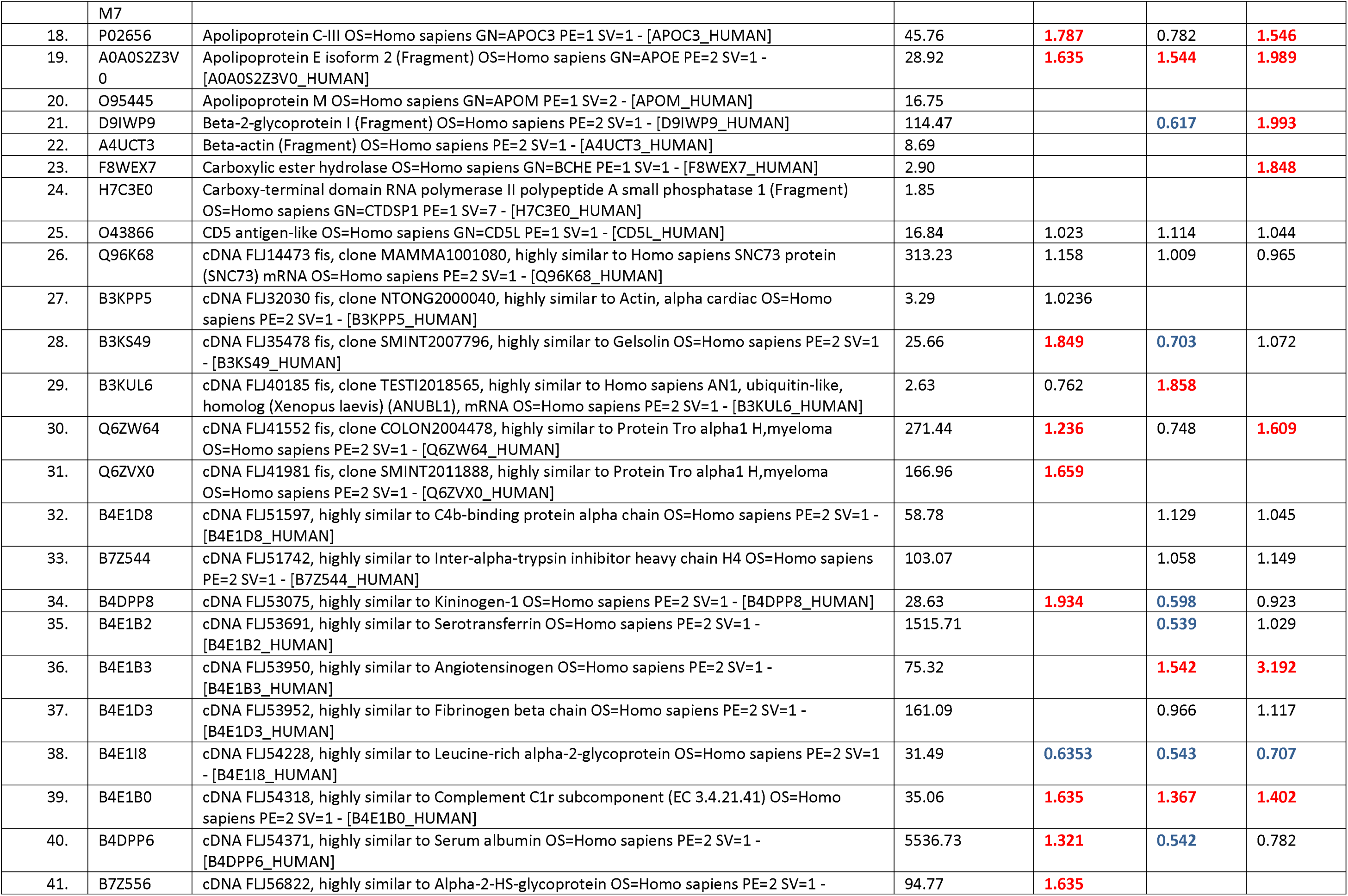
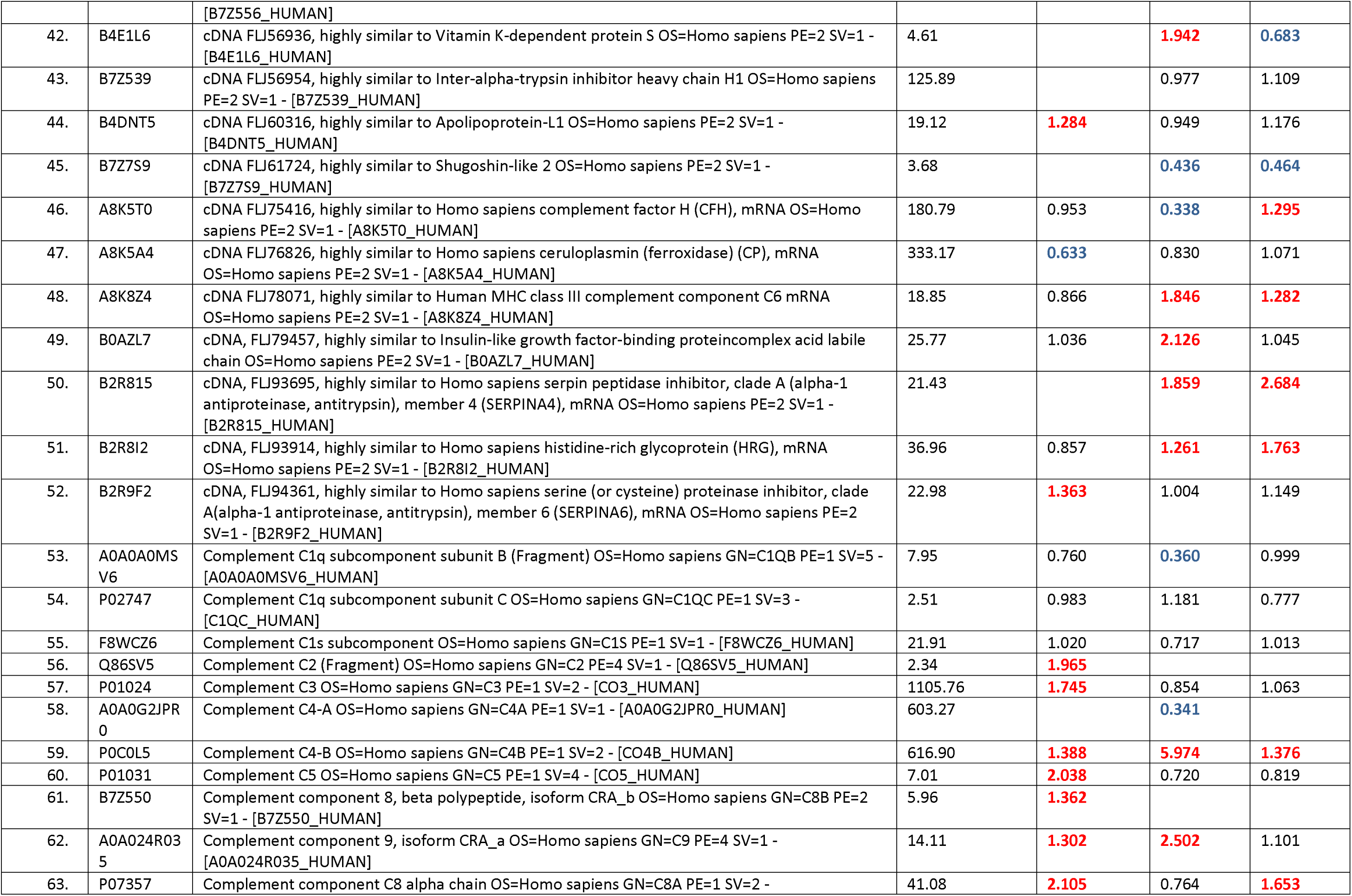
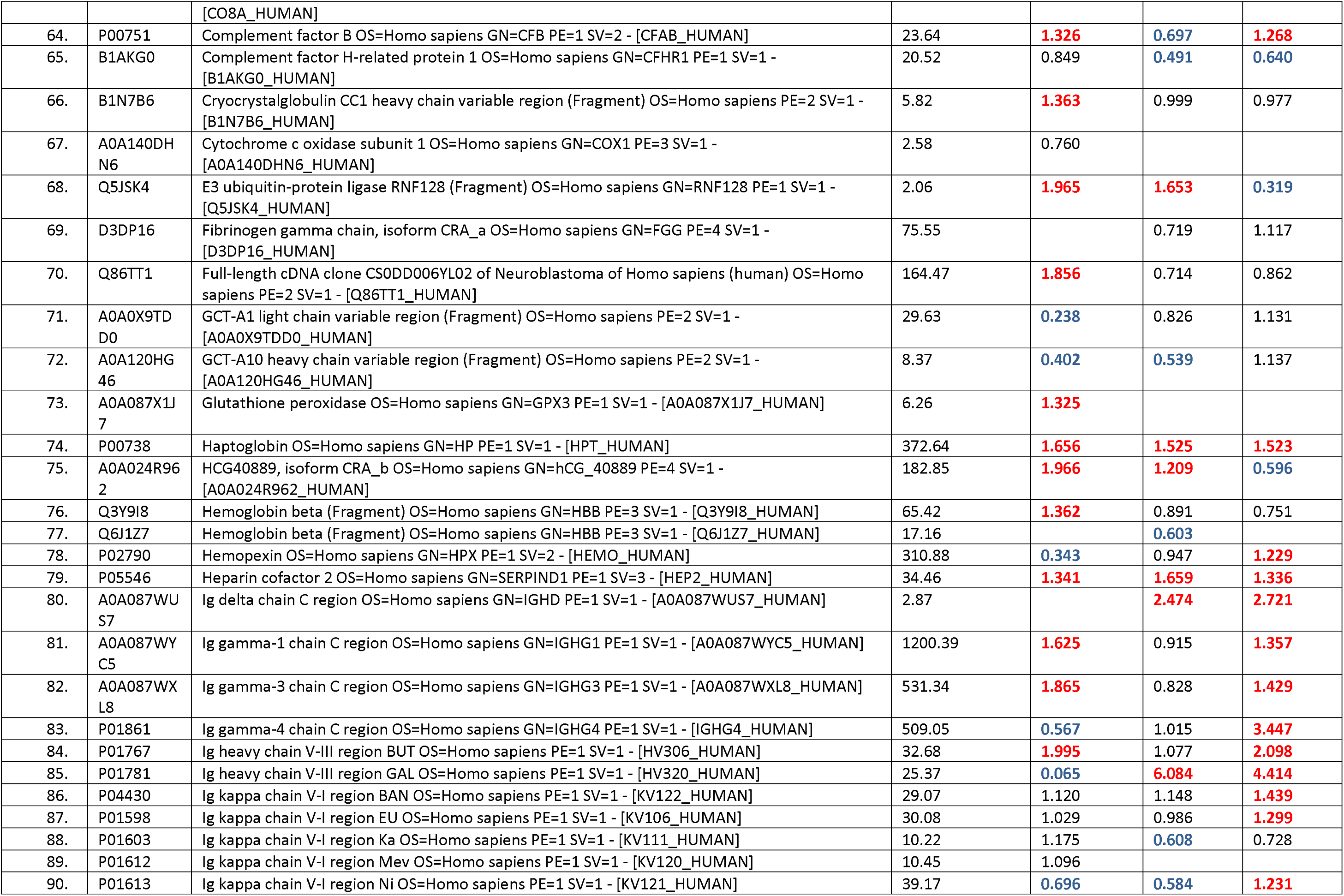
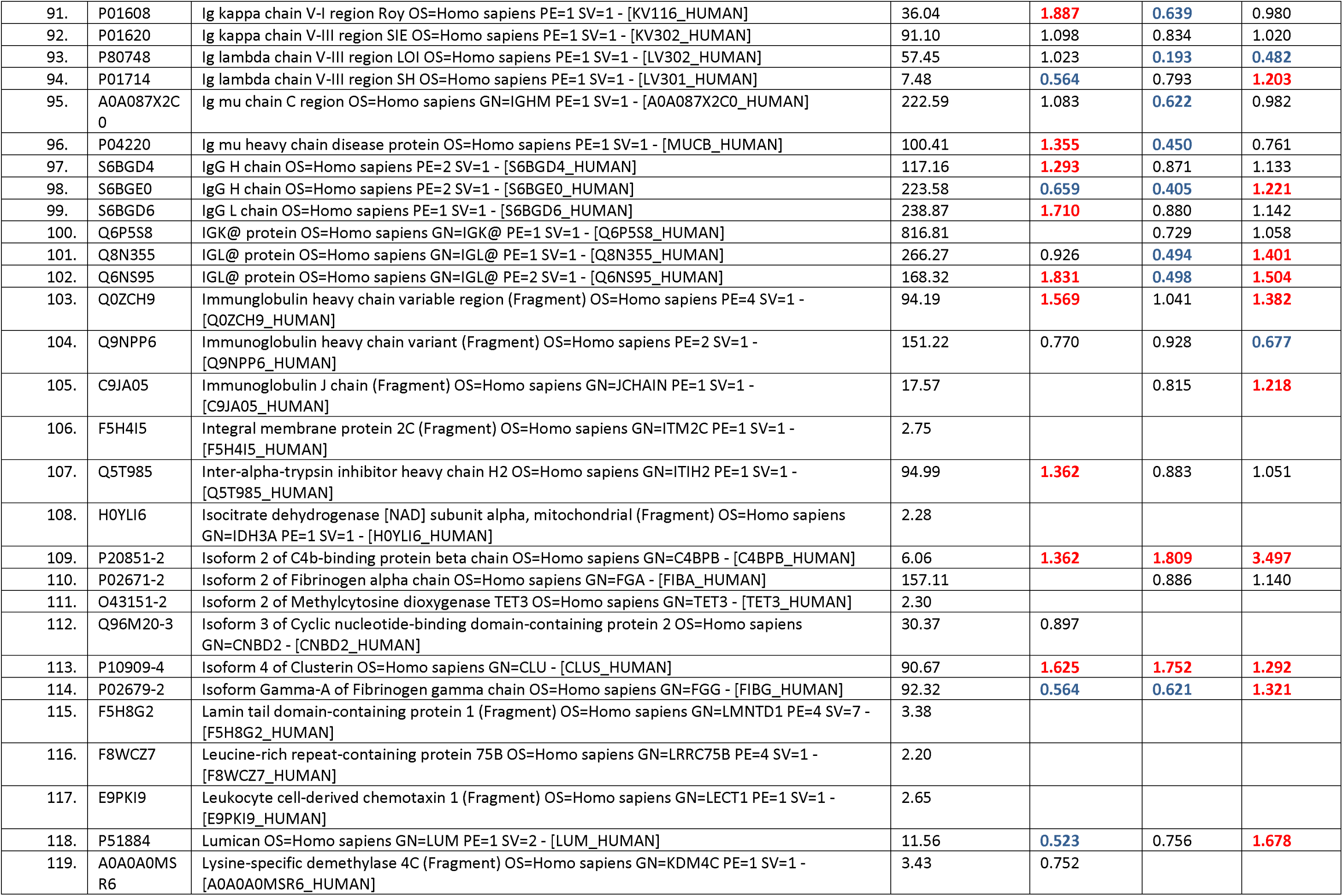
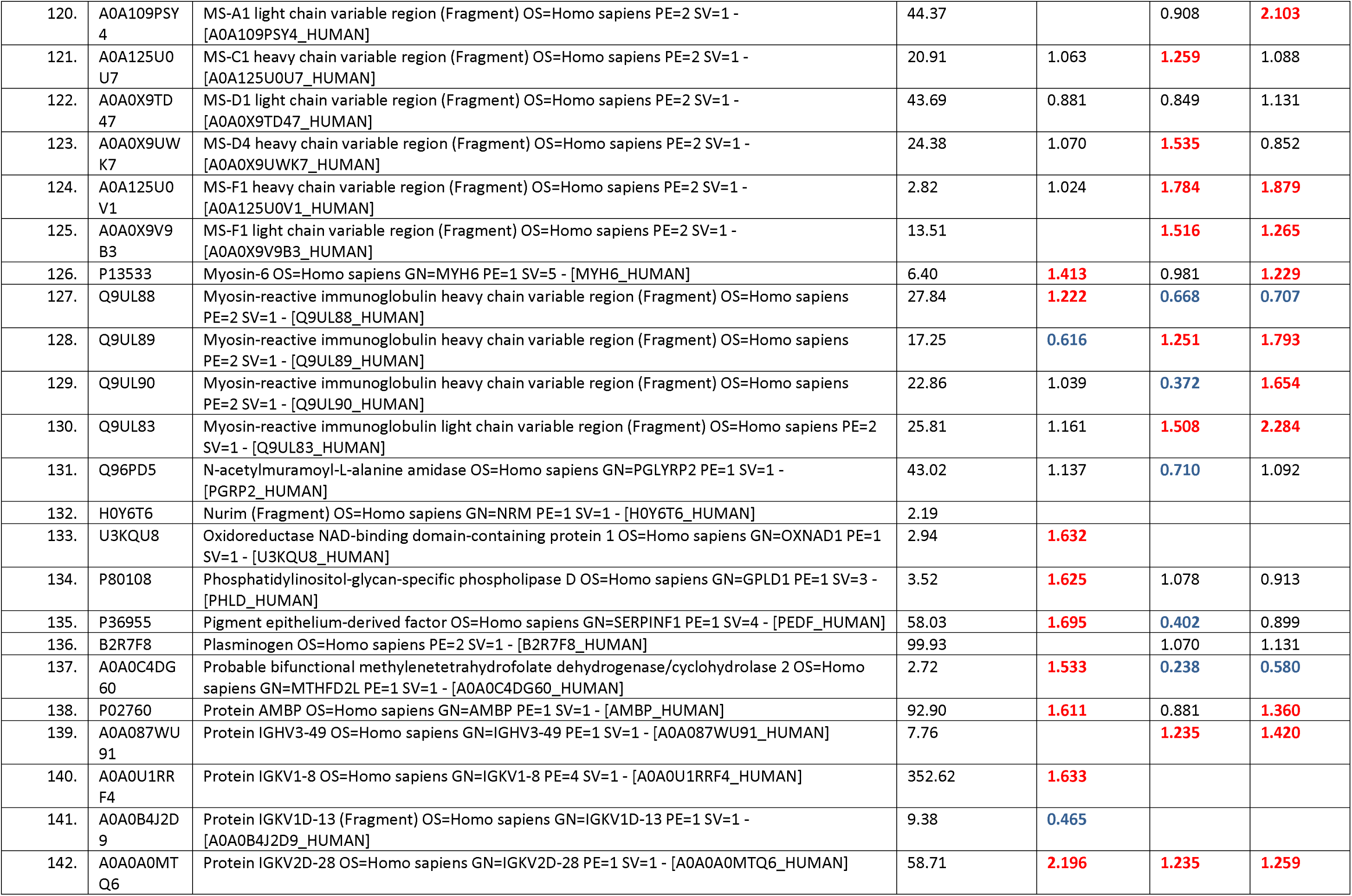
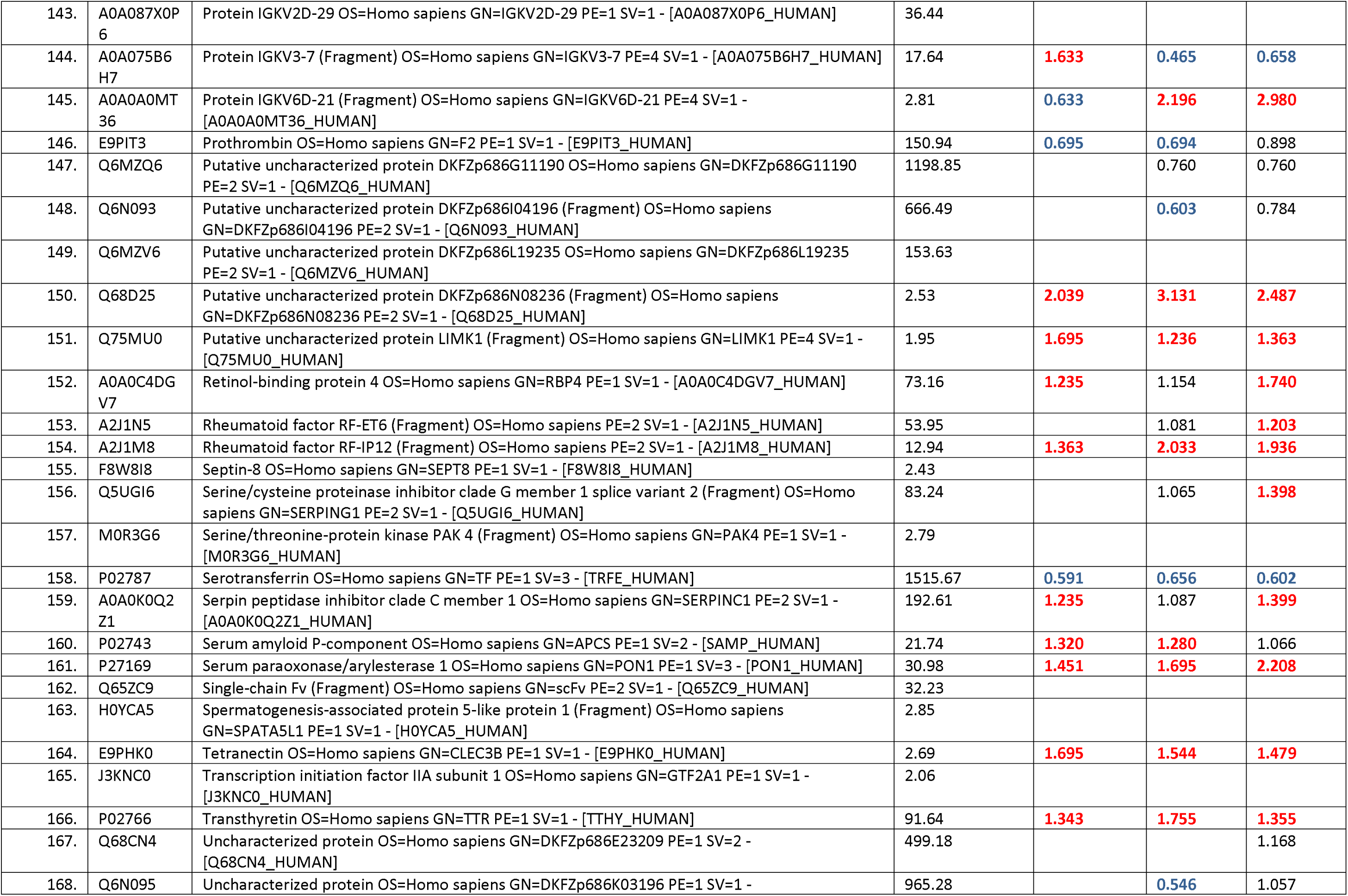
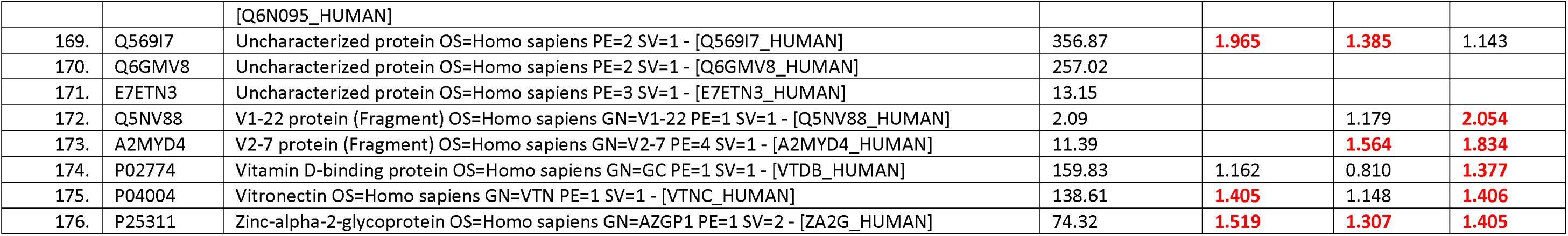
Differential Protein Expression Across Hearing Loss Severity Levels. Table displays the differential protein expression profiles for 176 proteins across three severity levels of hearing loss: mild, moderate, and severe. Each panel represents the fold change in protein expression levels compared to the normal hearing group. Blue color reparented downregulated and red color upregulated.

**Additional Table 3.** Summary of audiometry statistical data analysis repot.

**Additional Table 4.**
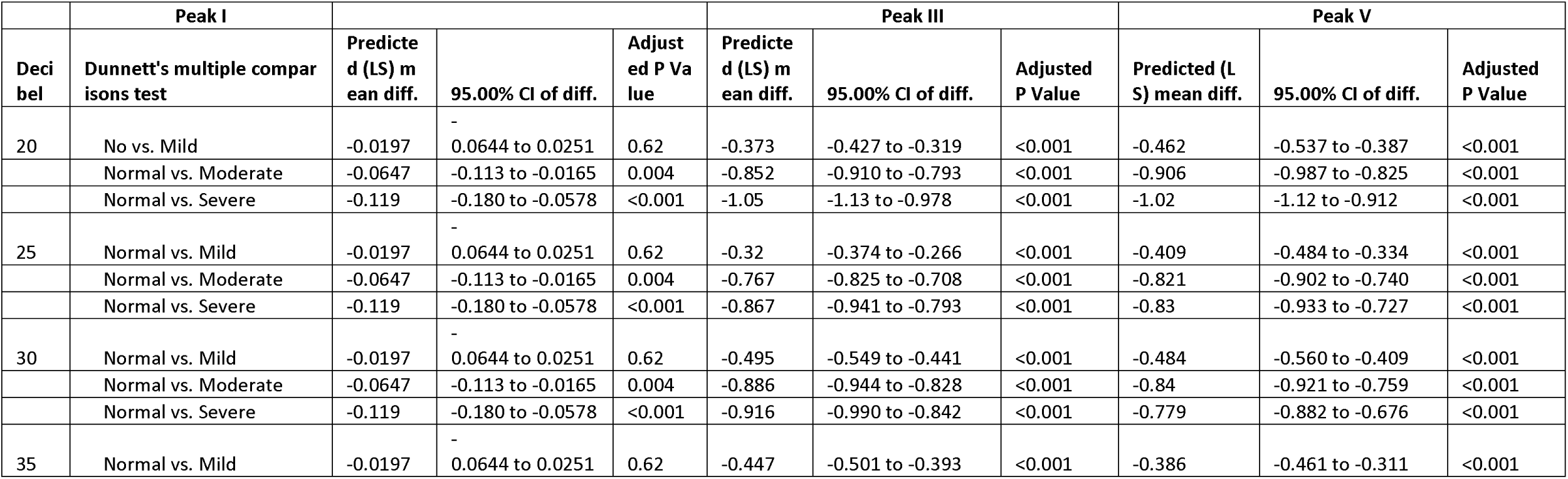
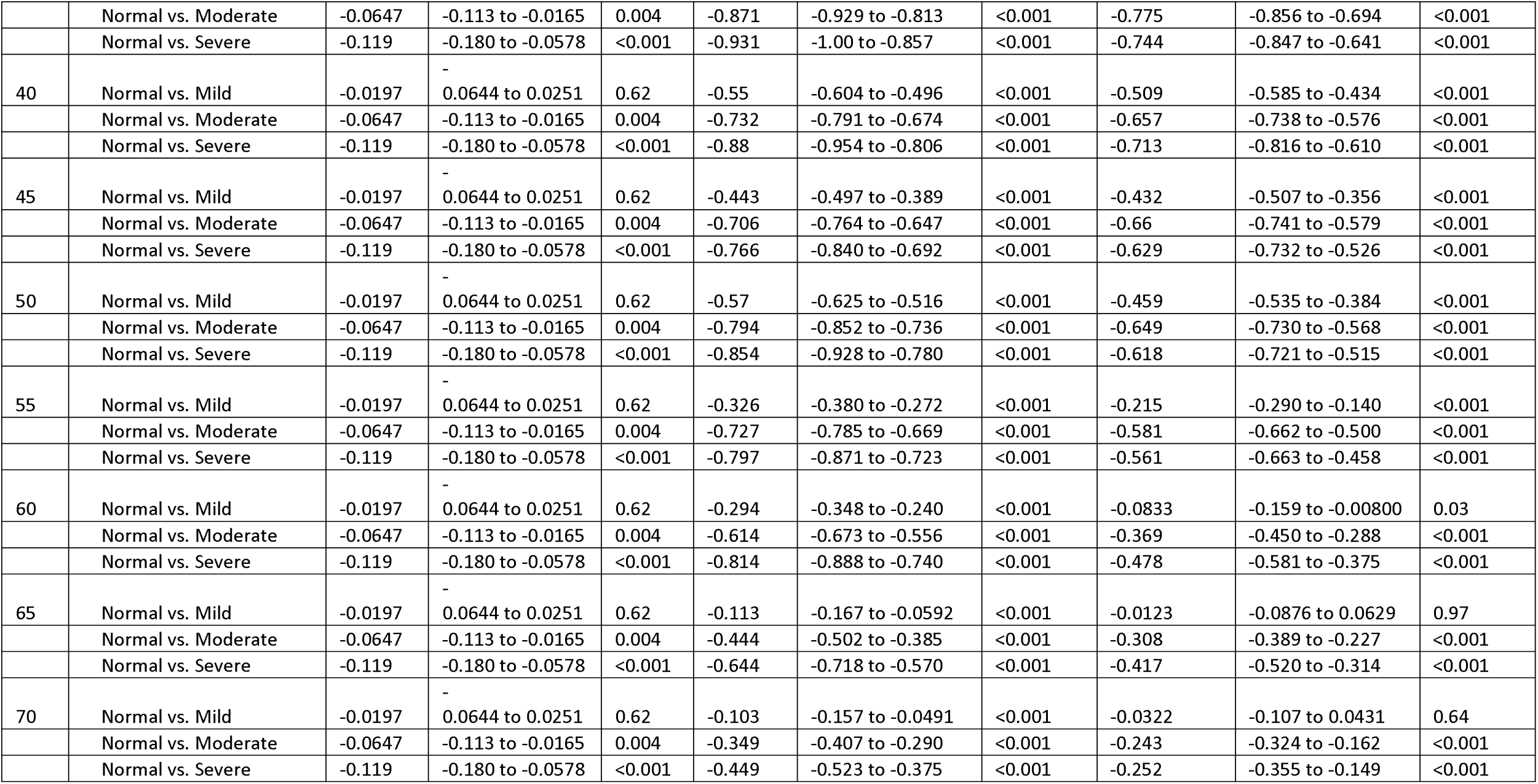
Individual Wave I-V Latency Statistical Analysis. The table presents the mean latencies, standard deviations, and results from statistical tests comparing latencies between normal and impaired hearing groups (mild, moderate, severe). For each wave, latency values are displayed along with their corresponding 95% confidence intervals (CI) and p-values, reflecting differences in wave latencies at each severity level.

**Additional Table 5.**
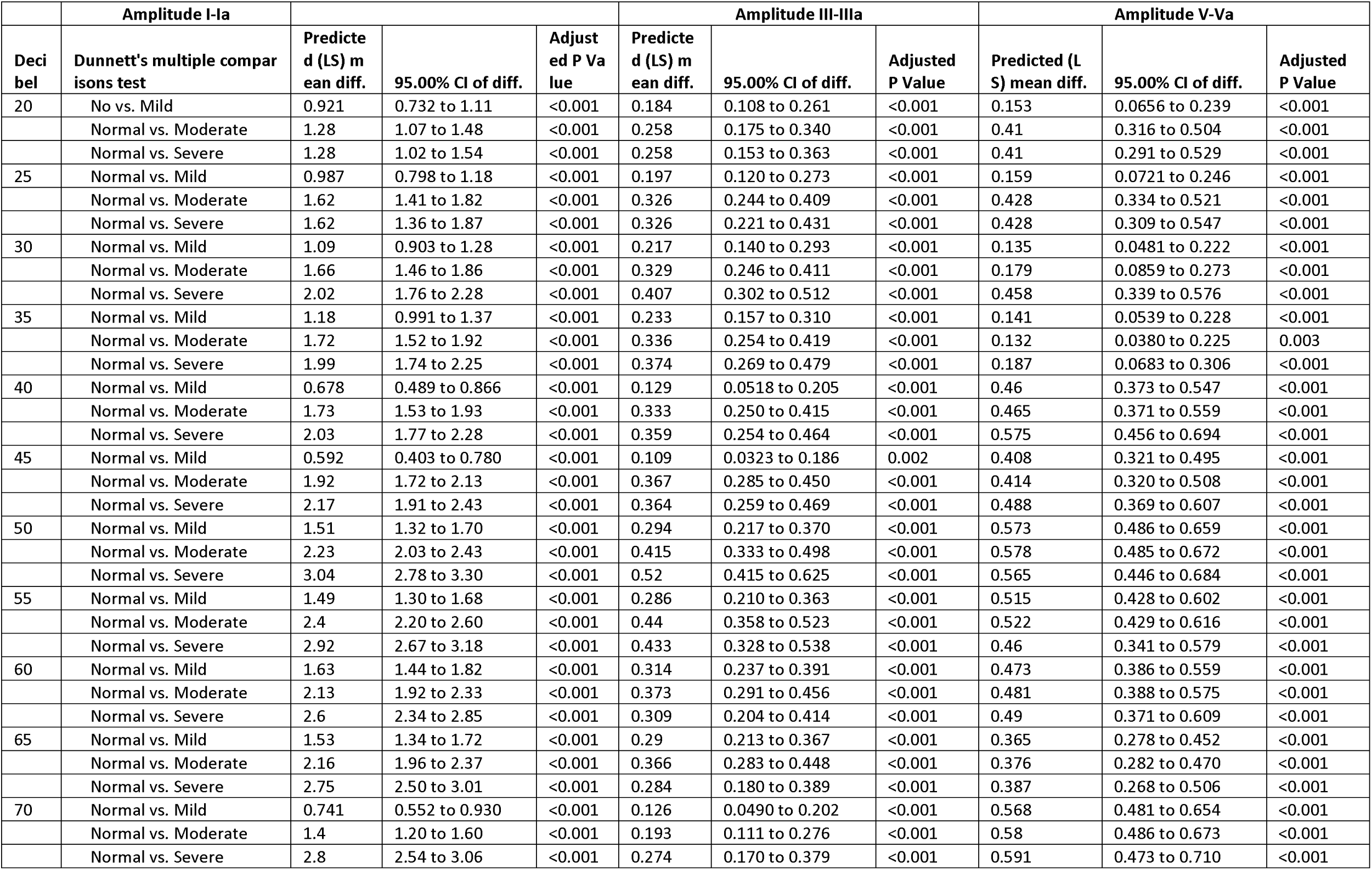
Individual Wave I-V Amplitude Statistical Analysis. The table presents the mean amplitude, standard deviations, and results from statistical tests comparing latencies between normal and impaired hearing groups (mild, moderate, severe). For each wave, amplitude values are displayed along with their corresponding 95% confidence intervals (CI) and p-values, reflecting differences in wave amplitude at each severity level.

## Notes

### Competing Interest Statement

The authors have declared no competing interest.

