## Supplemental Table for "Evaluating the Effects of Aircraft Noise on Hearing and Physiological Indicators: A Study of Military Personnel Using iTRAQ Proteomics and Cognitive Assessments"

**Additional Table 1 Summary of executing HRV parameters**

| Parameter | Units | Description |
| --- | --- | --- |
| **Time domain** | | |
| Mean RR | ms | The mean of RR intervals |
| SDNN | ms | Standard deviation of RR intervals |
| Mean HR | min^−1^ | The mean heart rate |
| STD HR | min^−1^ | Standard deviation of instantaneous heart rate values |
| Min HR | min^−1^ | Minimum HR computed using N beat moving average |
| Max HR | min^−1^ | Maximum HR computed using N beat moving average |
| RMSSD | ms | Square root of the mean squared differences between successive RR intervals |
| NN50 | Beats | Number of successive RR interval pairs that differ more than 50 ms |
| pNN50 | % | NNxx divided by the total number of RR intervals |
| HRV triangular index |  | The integral of the RR interval histogram divided by the height of the histogram |
| TINN | ms | Baseline width of the RR interval histogram |
| F**requency-domain—spectrum Welch’s periodogram** | | |
| Peak frequency | Hz | VLF, LF, and HF band peak frequencies |
| Absolute power | ms^2^ | Absolute powers of VLF, LF, and HF bands |
| Absolute power | log | Natural logarithm transformed values of absolute powers of VLF, LF, and HF bands |
| Relative power | % | Relative powers of VLF, LF, and HF bands |
| Normalized power | n.u | Powers of LF and HF bands in normalized units (VLF not included) |
| Total power | ms^2^ | Total absolute powers (VLF + LF + HF) |
| LF HF ratio |  | Ratio between LF and HF band powers |

**Additional Table 2:** Differential Protein Expression Across Hearing Loss Severity Levels. Table displays the differential protein expression profiles for 176 proteins across three severity levels of hearing loss: mild, moderate, and severe. Each panel represents the fold change in protein expression levels compared to the normal hearing group. Blue color reparented downregulated and red color upregulated.

|  | **Accession** | **Description** | **Score** | **114/113 (Mild)** | **115/113 (Moderate)** | **116/113 (Severe)** |
| --- | --- | --- | --- | --- | --- | --- |
| 1. | P43652 | Afamin OS=Homo sapiens GN=AFM PE=1 SV=1 - [AFAM_HUMAN] | 15.44 |  | 1.008 | 0.813 |
| 2. | F6KPG5 | Albumin (Fragment) OS=Homo sapiens PE=2 SV=1 - [F6KPG5_HUMAN] | 5376.07 | **0.425** | **0.446** | 0.818 |
| 3. | U3PXP0 | Alpha globin chain (Fragment) OS=Homo sapiens GN=HBA2 PE=3 SV=1 - [U3PXP0_HUMAN] | 9.14 | **1.236** | 0.842 | 0.896 |
| 4. | P02763 | Alpha-1-acid glycoprotein 1 OS=Homo sapiens GN=ORM1 PE=1 SV=1 - [A1AG1_HUMAN] | 135.82 | **1.996** | **1.560** | **1.711** |
| 5. | P19652 | Alpha-1-acid glycoprotein 2 OS=Homo sapiens GN=ORM2 PE=1 SV=2 - [A1AG2_HUMAN] | 65.84 | **1.637** | 0.952 | **1.294** |
| 6. | P01011 | Alpha-1-antichymotrypsin OS=Homo sapiens GN=SERPINA3 PE=1 SV=2 - [AACT_HUMAN] | 137.96 | 1.023 | 0.971 | 1.040 |
| 7. | A0A024R6I7 | Alpha-1-antitrypsin OS=Homo sapiens GN=SERPINA1 PE=1 SV=1 - [A0A024R6I7_HUMAN] | 414.89 | **1.363** | **1.350** | **0.616** |
| 8. | P01009 | Alpha-1-antitrypsin OS=Homo sapiens GN=SERPINA1 PE=1 SV=3 - [A1AT_HUMAN] | 437.53 | **0.602** | **0.222** | **1.223** |
| 9. | P04217 | Alpha-1B-glycoprotein OS=Homo sapiens GN=A1BG PE=1 SV=4 - [A1BG_HUMAN] | 78.47 |  | 0.894 | 1.162 |
| 10. | P08697 | Alpha-2-antiplasmin OS=Homo sapiens GN=SERPINF2 PE=1 SV=3 - [A2AP_HUMAN] | 50.71 | **3.026** | **0.614** | 0.770 |
| 11. | P02765 | Alpha-2-HS-glycoprotein OS=Homo sapiens GN=AHSG PE=1 SV=1 - [FETUA_HUMAN] | 228.87 | **1.443** | 0.717 | 1.138 |
| 12. | P01023 | Alpha-2-macroglobulin OS=Homo sapiens GN=A2M PE=1 SV=3 - [A2MG_HUMAN] | 938.27 | **0.462** | **0.472** | 0.898 |
| 13. | A2KBC2 | Anti-(ED-B) scFV (Fragment) OS=Homo sapiens PE=2 SV=1 - [A2KBC2_HUMAN] | 91.97 | 1.033 | 1.092 | 1.159 |
| 14. | P02647 | Apolipoprotein A-I OS=Homo sapiens GN=APOA1 PE=1 SV=1 - [APOA1_HUMAN] | 241.56 |  | **0.695** | 0.927 |
| 15. | P02652 | Apolipoprotein A-II OS=Homo sapiens GN=APOA2 PE=1 SV=1 - [APOA2_HUMAN] | 58.35 |  | **0.565** | 0.960 |
| 16. | P06727 | Apolipoprotein A-IV OS=Homo sapiens GN=APOA4 PE=1 SV=3 - [APOA4_HUMAN] | 8.89 | **1.491** | 0.762 | 0.732 |
| 17. | A0A087WTM7 | Apolipoprotein B-100 OS=Homo sapiens GN=APOB PE=1 SV=1 - [A0A087WTM7_HUMAN] | 465.23 | **1.235** | 0.834 | 0.758 |
| 18. | P02656 | Apolipoprotein C-III OS=Homo sapiens GN=APOC3 PE=1 SV=1 - [APOC3_HUMAN] | 45.76 | **1.787** | 0.782 | **1.546** |
| 19. | A0A0S2Z3V0 | Apolipoprotein E isoform 2 (Fragment) OS=Homo sapiens GN=APOE PE=2 SV=1 - [A0A0S2Z3V0_HUMAN] | 28.92 | **1.635** | **1.544** | **1.989** |
| 20. | O95445 | Apolipoprotein M OS=Homo sapiens GN=APOM PE=1 SV=2 - [APOM_HUMAN] | 16.75 |  |  |  |
| 21. | D9IWP9 | Beta-2-glycoprotein I (Fragment) OS=Homo sapiens PE=2 SV=1 - [D9IWP9_HUMAN] | 114.47 |  | **0.617** | **1.993** |
| 22. | A4UCT3 | Beta-actin (Fragment) OS=Homo sapiens PE=2 SV=1 - [A4UCT3_HUMAN] | 8.69 |  |  |  |
| 23. | F8WEX7 | Carboxylic ester hydrolase OS=Homo sapiens GN=BCHE PE=1 SV=1 - [F8WEX7_HUMAN] | 2.90 |  |  | **1.848** |
| 24. | H7C3E0 | Carboxy-terminal domain RNA polymerase II polypeptide A small phosphatase 1 (Fragment) OS=Homo sapiens GN=CTDSP1 PE=1 SV=7 - [H7C3E0_HUMAN] | 1.85 |  |  |  |
| 25. | O43866 | CD5 antigen-like OS=Homo sapiens GN=CD5L PE=1 SV=1 - [CD5L_HUMAN] | 16.84 | 1.023 | 1.114 | 1.044 |
| 26. | Q96K68 | cDNA FLJ14473 fis, clone MAMMA1001080, highly similar to Homo sapiens SNC73 protein (SNC73) mRNA OS=Homo sapiens PE=2 SV=1 - [Q96K68_HUMAN] | 313.23 | 1.158 | 1.009 | 0.965 |
| 27. | B3KPP5 | cDNA FLJ32030 fis, clone NTONG2000040, highly similar to Actin, alpha cardiac OS=Homo sapiens PE=2 SV=1 - [B3KPP5_HUMAN] | 3.29 | 1.0236 |  |  |
| 28. | B3KS49 | cDNA FLJ35478 fis, clone SMINT2007796, highly similar to Gelsolin OS=Homo sapiens PE=2 SV=1 - [B3KS49_HUMAN] | 25.66 | **1.849** | **0.703** | 1.072 |
| 29. | B3KUL6 | cDNA FLJ40185 fis, clone TESTI2018565, highly similar to Homo sapiens AN1, ubiquitin-like, homolog (Xenopus laevis) (ANUBL1), mRNA OS=Homo sapiens PE=2 SV=1 - [B3KUL6_HUMAN] | 2.63 | 0.762 | **1.858** |  |
| 30. | Q6ZW64 | cDNA FLJ41552 fis, clone COLON2004478, highly similar to Protein Tro alpha1 H,myeloma OS=Homo sapiens PE=2 SV=1 - [Q6ZW64_HUMAN] | 271.44 | **1.236** | 0.748 | **1.609** |
| 31. | Q6ZVX0 | cDNA FLJ41981 fis, clone SMINT2011888, highly similar to Protein Tro alpha1 H,myeloma OS=Homo sapiens PE=2 SV=1 - [Q6ZVX0_HUMAN] | 166.96 | **1.659** |  |  |
| 32. | B4E1D8 | cDNA FLJ51597, highly similar to C4b-binding protein alpha chain OS=Homo sapiens PE=2 SV=1 - [B4E1D8_HUMAN] | 58.78 |  | 1.129 | 1.045 |
| 33. | B7Z544 | cDNA FLJ51742, highly similar to Inter-alpha-trypsin inhibitor heavy chain H4 OS=Homo sapiens PE=2 SV=1 - [B7Z544_HUMAN] | 103.07 |  | 1.058 | 1.149 |
| 34. | B4DPP8 | cDNA FLJ53075, highly similar to Kininogen-1 OS=Homo sapiens PE=2 SV=1 - [B4DPP8_HUMAN] | 28.63 | **1.934** | **0.598** | 0.923 |
| 35. | B4E1B2 | cDNA FLJ53691, highly similar to Serotransferrin OS=Homo sapiens PE=2 SV=1 - [B4E1B2_HUMAN] | 1515.71 |  | **0.539** | 1.029 |
| 36. | B4E1B3 | cDNA FLJ53950, highly similar to Angiotensinogen OS=Homo sapiens PE=2 SV=1 - [B4E1B3_HUMAN] | 75.32 |  | **1.542** | **3.192** |
| 37. | B4E1D3 | cDNA FLJ53952, highly similar to Fibrinogen beta chain OS=Homo sapiens PE=2 SV=1 - [B4E1D3_HUMAN] | 161.09 |  | 0.966 | 1.117 |
| 38. | B4E1I8 | cDNA FLJ54228, highly similar to Leucine-rich alpha-2-glycoprotein OS=Homo sapiens PE=2 SV=1 - [B4E1I8_HUMAN] | 31.49 | **0.6353** | **0.543** | **0.707** |
| 39. | B4E1B0 | cDNA FLJ54318, highly similar to Complement C1r subcomponent (EC 3.4.21.41) OS=Homo sapiens PE=2 SV=1 - [B4E1B0_HUMAN] | 35.06 | **1.635** | **1.367** | **1.402** |
| 40. | B4DPP6 | cDNA FLJ54371, highly similar to Serum albumin OS=Homo sapiens PE=2 SV=1 - [B4DPP6_HUMAN] | 5536.73 | **1.321** | **0.542** | 0.782 |
| 41. | B7Z556 | cDNA FLJ56822, highly similar to Alpha-2-HS-glycoprotein OS=Homo sapiens PE=2 SV=1 - [B7Z556_HUMAN] | 94.77 | **1.635** |  |  |
| 42. | B4E1L6 | cDNA FLJ56936, highly similar to Vitamin K-dependent protein S OS=Homo sapiens PE=2 SV=1 - [B4E1L6_HUMAN] | 4.61 |  | **1.942** | **0.683** |
| 43. | B7Z539 | cDNA FLJ56954, highly similar to Inter-alpha-trypsin inhibitor heavy chain H1 OS=Homo sapiens PE=2 SV=1 - [B7Z539_HUMAN] | 125.89 |  | 0.977 | 1.109 |
| 44. | B4DNT5 | cDNA FLJ60316, highly similar to Apolipoprotein-L1 OS=Homo sapiens PE=2 SV=1 - [B4DNT5_HUMAN] | 19.12 | **1.284** | 0.949 | 1.176 |
| 45. | B7Z7S9 | cDNA FLJ61724, highly similar to Shugoshin-like 2 OS=Homo sapiens PE=2 SV=1 - [B7Z7S9_HUMAN] | 3.68 |  | **0.436** | **0.464** |
| 46. | A8K5T0 | cDNA FLJ75416, highly similar to Homo sapiens complement factor H (CFH), mRNA OS=Homo sapiens PE=2 SV=1 - [A8K5T0_HUMAN] | 180.79 | 0.953 | **0.338** | **1.295** |
| 47. | A8K5A4 | cDNA FLJ76826, highly similar to Homo sapiens ceruloplasmin (ferroxidase) (CP), mRNA OS=Homo sapiens PE=2 SV=1 - [A8K5A4_HUMAN] | 333.17 | **0.633** | 0.830 | 1.071 |
| 48. | A8K8Z4 | cDNA FLJ78071, highly similar to Human MHC class III complement component C6 mRNA OS=Homo sapiens PE=2 SV=1 - [A8K8Z4_HUMAN] | 18.85 | 0.866 | **1.846** | **1.282** |
| 49. | B0AZL7 | cDNA, FLJ79457, highly similar to Insulin-like growth factor-binding proteincomplex acid labile chain OS=Homo sapiens PE=2 SV=1 - [B0AZL7_HUMAN] | 25.77 | 1.036 | **2.126** | 1.045 |
| 50. | B2R815 | cDNA, FLJ93695, highly similar to Homo sapiens serpin peptidase inhibitor, clade A (alpha-1 antiproteinase, antitrypsin), member 4 (SERPINA4), mRNA OS=Homo sapiens PE=2 SV=1 - [B2R815_HUMAN] | 21.43 |  | **1.859** | **2.684** |
| 51. | B2R8I2 | cDNA, FLJ93914, highly similar to Homo sapiens histidine-rich glycoprotein (HRG), mRNA OS=Homo sapiens PE=2 SV=1 - [B2R8I2_HUMAN] | 36.96 | 0.857 | **1.261** | **1.763** |
| 52. | B2R9F2 | cDNA, FLJ94361, highly similar to Homo sapiens serine (or cysteine) proteinase inhibitor, clade A(alpha-1 antiproteinase, antitrypsin), member 6 (SERPINA6), mRNA OS=Homo sapiens PE=2 SV=1 - [B2R9F2_HUMAN] | 22.98 | **1.363** | 1.004 | 1.149 |
| 53. | A0A0A0MSV6 | Complement C1q subcomponent subunit B (Fragment) OS=Homo sapiens GN=C1QB PE=1 SV=5 - [A0A0A0MSV6_HUMAN] | 7.95 | 0.760 | **0.360** | 0.999 |
| 54. | P02747 | Complement C1q subcomponent subunit C OS=Homo sapiens GN=C1QC PE=1 SV=3 - [C1QC_HUMAN] | 2.51 | 0.983 | 1.181 | 0.777 |
| 55. | F8WCZ6 | Complement C1s subcomponent OS=Homo sapiens GN=C1S PE=1 SV=1 - [F8WCZ6_HUMAN] | 21.91 | 1.020 | 0.717 | 1.013 |
| 56. | Q86SV5 | Complement C2 (Fragment) OS=Homo sapiens GN=C2 PE=4 SV=1 - [Q86SV5_HUMAN] | 2.34 | **1.965** |  |  |
| 57. | P01024 | Complement C3 OS=Homo sapiens GN=C3 PE=1 SV=2 - [CO3_HUMAN] | 1105.76 | **1.745** | 0.854 | 1.063 |
| 58. | A0A0G2JPR0 | Complement C4-A OS=Homo sapiens GN=C4A PE=1 SV=1 - [A0A0G2JPR0_HUMAN] | 603.27 |  | **0.341** |  |
| 59. | P0C0L5 | Complement C4-B OS=Homo sapiens GN=C4B PE=1 SV=2 - [CO4B_HUMAN] | 616.90 | **1.388** | **5.974** | **1.376** |
| 60. | P01031 | Complement C5 OS=Homo sapiens GN=C5 PE=1 SV=4 - [CO5_HUMAN] | 7.01 | **2.038** | 0.720 | 0.819 |
| 61. | B7Z550 | Complement component 8, beta polypeptide, isoform CRA_b OS=Homo sapiens GN=C8B PE=2 SV=1 - [B7Z550_HUMAN] | 5.96 | **1.362** |  |  |
| 62. | A0A024R035 | Complement component 9, isoform CRA_a OS=Homo sapiens GN=C9 PE=4 SV=1 - [A0A024R035_HUMAN] | 14.11 | **1.302** | **2.502** | 1.101 |
| 63. | P07357 | Complement component C8 alpha chain OS=Homo sapiens GN=C8A PE=1 SV=2 - [CO8A_HUMAN] | 41.08 | **2.105** | 0.764 | **1.653** |
| 64. | P00751 | Complement factor B OS=Homo sapiens GN=CFB PE=1 SV=2 - [CFAB_HUMAN] | 23.64 | **1.326** | **0.697** | **1.268** |
| 65. | B1AKG0 | Complement factor H-related protein 1 OS=Homo sapiens GN=CFHR1 PE=1 SV=1 - [B1AKG0_HUMAN] | 20.52 | 0.849 | **0.491** | **0.640** |
| 66. | B1N7B6 | Cryocrystalglobulin CC1 heavy chain variable region (Fragment) OS=Homo sapiens PE=2 SV=1 - [B1N7B6_HUMAN] | 5.82 | **1.363** | 0.999 | 0.977 |
| 67. | A0A140DHN6 | Cytochrome c oxidase subunit 1 OS=Homo sapiens GN=COX1 PE=3 SV=1 - [A0A140DHN6_HUMAN] | 2.58 | 0.760 |  |  |
| 68. | Q5JSK4 | E3 ubiquitin-protein ligase RNF128 (Fragment) OS=Homo sapiens GN=RNF128 PE=1 SV=1 - [Q5JSK4_HUMAN] | 2.06 | **1.965** | **1.653** | **0.319** |
| 69. | D3DP16 | Fibrinogen gamma chain, isoform CRA_a OS=Homo sapiens GN=FGG PE=4 SV=1 - [D3DP16_HUMAN] | 75.55 |  | 0.719 | 1.117 |
| 70. | Q86TT1 | Full-length cDNA clone CS0DD006YL02 of Neuroblastoma of Homo sapiens (human) OS=Homo sapiens PE=2 SV=1 - [Q86TT1_HUMAN] | 164.47 | **1.856** | 0.714 | 0.862 |
| 71. | A0A0X9TDD0 | GCT-A1 light chain variable region (Fragment) OS=Homo sapiens PE=2 SV=1 - [A0A0X9TDD0_HUMAN] | 29.63 | **0.238** | 0.826 | 1.131 |
| 72. | A0A120HG46 | GCT-A10 heavy chain variable region (Fragment) OS=Homo sapiens PE=2 SV=1 - [A0A120HG46_HUMAN] | 8.37 | **0.402** | **0.539** | 1.137 |
| 73. | A0A087X1J7 | Glutathione peroxidase OS=Homo sapiens GN=GPX3 PE=1 SV=1 - [A0A087X1J7_HUMAN] | 6.26 | **1.325** |  |  |
| 74. | P00738 | Haptoglobin OS=Homo sapiens GN=HP PE=1 SV=1 - [HPT_HUMAN] | 372.64 | **1.656** | **1.525** | **1.523** |
| 75. | A0A024R962 | HCG40889, isoform CRA_b OS=Homo sapiens GN=hCG_40889 PE=4 SV=1 - [A0A024R962_HUMAN] | 182.85 | **1.966** | **1.209** | **0.596** |
| 76. | Q3Y9I8 | Hemoglobin beta (Fragment) OS=Homo sapiens GN=HBB PE=3 SV=1 - [Q3Y9I8_HUMAN] | 65.42 | **1.362** | 0.891 | 0.751 |
| 77. | Q6J1Z7 | Hemoglobin beta (Fragment) OS=Homo sapiens GN=HBB PE=3 SV=1 - [Q6J1Z7_HUMAN] | 17.16 |  | **0.603** |  |
| 78. | P02790 | Hemopexin OS=Homo sapiens GN=HPX PE=1 SV=2 - [HEMO_HUMAN] | 310.88 | **0.343** | 0.947 | **1.229** |
| 79. | P05546 | Heparin cofactor 2 OS=Homo sapiens GN=SERPIND1 PE=1 SV=3 - [HEP2_HUMAN] | 34.46 | **1.341** | **1.659** | **1.336** |
| 80. | A0A087WUS7 | Ig delta chain C region OS=Homo sapiens GN=IGHD PE=1 SV=1 - [A0A087WUS7_HUMAN] | 2.87 |  | **2.474** | **2.721** |
| 81. | A0A087WYC5 | Ig gamma-1 chain C region OS=Homo sapiens GN=IGHG1 PE=1 SV=1 - [A0A087WYC5_HUMAN] | 1200.39 | **1.625** | 0.915 | **1.357** |
| 82. | A0A087WXL8 | Ig gamma-3 chain C region OS=Homo sapiens GN=IGHG3 PE=1 SV=1 - [A0A087WXL8_HUMAN] | 531.34 | **1.865** | 0.828 | **1.429** |
| 83. | P01861 | Ig gamma-4 chain C region OS=Homo sapiens GN=IGHG4 PE=1 SV=1 - [IGHG4_HUMAN] | 509.05 | **0.567** | 1.015 | **3.447** |
| 84. | P01767 | Ig heavy chain V-III region BUT OS=Homo sapiens PE=1 SV=1 - [HV306_HUMAN] | 32.68 | **1.995** | 1.077 | **2.098** |
| 85. | P01781 | Ig heavy chain V-III region GAL OS=Homo sapiens PE=1 SV=1 - [HV320_HUMAN] | 25.37 | **0.065** | **6.084** | **4.414** |
| 86. | P04430 | Ig kappa chain V-I region BAN OS=Homo sapiens PE=1 SV=1 - [KV122_HUMAN] | 29.07 | 1.120 | 1.148 | **1.439** |
| 87. | P01598 | Ig kappa chain V-I region EU OS=Homo sapiens PE=1 SV=1 - [KV106_HUMAN] | 30.08 | 1.029 | 0.986 | **1.299** |
| 88. | P01603 | Ig kappa chain V-I region Ka OS=Homo sapiens PE=1 SV=1 - [KV111_HUMAN] | 10.22 | 1.175 | **0.608** | 0.728 |
| 89. | P01612 | Ig kappa chain V-I region Mev OS=Homo sapiens PE=1 SV=1 - [KV120_HUMAN] | 10.45 | 1.096 |  |  |
| 90. | P01613 | Ig kappa chain V-I region Ni OS=Homo sapiens PE=1 SV=1 - [KV121_HUMAN] | 39.17 | **0.696** | **0.584** | **1.231** |
| 91. | P01608 | Ig kappa chain V-I region Roy OS=Homo sapiens PE=1 SV=1 - [KV116_HUMAN] | 36.04 | **1.887** | **0.639** | 0.980 |
| 92. | P01620 | Ig kappa chain V-III region SIE OS=Homo sapiens PE=1 SV=1 - [KV302_HUMAN] | 91.10 | 1.098 | 0.834 | 1.020 |
| 93. | P80748 | Ig lambda chain V-III region LOI OS=Homo sapiens PE=1 SV=1 - [LV302_HUMAN] | 57.45 | 1.023 | **0.193** | **0.482** |
| 94. | P01714 | Ig lambda chain V-III region SH OS=Homo sapiens PE=1 SV=1 - [LV301_HUMAN] | 7.48 | **0.564** | 0.793 | **1.203** |
| 95. | A0A087X2C0 | Ig mu chain C region OS=Homo sapiens GN=IGHM PE=1 SV=1 - [A0A087X2C0_HUMAN] | 222.59 | 1.083 | **0.622** | 0.982 |
| 96. | P04220 | Ig mu heavy chain disease protein OS=Homo sapiens PE=1 SV=1 - [MUCB_HUMAN] | 100.41 | **1.355** | **0.450** | 0.761 |
| 97. | S6BGD4 | IgG H chain OS=Homo sapiens PE=2 SV=1 - [S6BGD4_HUMAN] | 117.16 | **1.293** | 0.871 | 1.133 |
| 98. | S6BGE0 | IgG H chain OS=Homo sapiens PE=2 SV=1 - [S6BGE0_HUMAN] | 223.58 | **0.659** | **0.405** | **1.221** |
| 99. | S6BGD6 | IgG L chain OS=Homo sapiens PE=1 SV=1 - [S6BGD6_HUMAN] | 238.87 | **1.710** | 0.880 | 1.142 |
| 100. | Q6P5S8 | IGK@ protein OS=Homo sapiens GN=IGK@ PE=1 SV=1 - [Q6P5S8_HUMAN] | 816.81 |  | 0.729 | 1.058 |
| 101. | Q8N355 | IGL@ protein OS=Homo sapiens GN=IGL@ PE=1 SV=1 - [Q8N355_HUMAN] | 266.27 | 0.926 | **0.494** | **1.401** |
| 102. | Q6NS95 | IGL@ protein OS=Homo sapiens GN=IGL@ PE=2 SV=1 - [Q6NS95_HUMAN] | 168.32 | **1.831** | **0.498** | **1.504** |
| 103. | Q0ZCH9 | Immunglobulin heavy chain variable region (Fragment) OS=Homo sapiens PE=4 SV=1 - [Q0ZCH9_HUMAN] | 94.19 | **1.569** | 1.041 | **1.382** |
| 104. | Q9NPP6 | Immunoglobulin heavy chain variant (Fragment) OS=Homo sapiens PE=2 SV=1 - [Q9NPP6_HUMAN] | 151.22 | 0.770 | 0.928 | **0.677** |
| 105. | C9JA05 | Immunoglobulin J chain (Fragment) OS=Homo sapiens GN=JCHAIN PE=1 SV=1 - [C9JA05_HUMAN] | 17.57 |  | 0.815 | **1.218** |
| 106. | F5H4I5 | Integral membrane protein 2C (Fragment) OS=Homo sapiens GN=ITM2C PE=1 SV=1 - [F5H4I5_HUMAN] | 2.75 |  |  |  |
| 107. | Q5T985 | Inter-alpha-trypsin inhibitor heavy chain H2 OS=Homo sapiens GN=ITIH2 PE=1 SV=1 - [Q5T985_HUMAN] | 94.99 | **1.362** | 0.883 | 1.051 |
| 108. | H0YLI6 | Isocitrate dehydrogenase [NAD] subunit alpha, mitochondrial (Fragment) OS=Homo sapiens GN=IDH3A PE=1 SV=1 - [H0YLI6_HUMAN] | 2.28 |  |  |  |
| 109. | P20851-2 | Isoform 2 of C4b-binding protein beta chain OS=Homo sapiens GN=C4BPB - [C4BPB_HUMAN] | 6.06 | **1.362** | **1.809** | **3.497** |
| 110. | P02671-2 | Isoform 2 of Fibrinogen alpha chain OS=Homo sapiens GN=FGA - [FIBA_HUMAN] | 157.11 |  | 0.886 | 1.140 |
| 111. | O43151-2 | Isoform 2 of Methylcytosine dioxygenase TET3 OS=Homo sapiens GN=TET3 - [TET3_HUMAN] | 2.30 |  |  |  |
| 112. | Q96M20-3 | Isoform 3 of Cyclic nucleotide-binding domain-containing protein 2 OS=Homo sapiens GN=CNBD2 - [CNBD2_HUMAN] | 30.37 | 0.897 |  |  |
| 113. | P10909-4 | Isoform 4 of Clusterin OS=Homo sapiens GN=CLU - [CLUS_HUMAN] | 90.67 | **1.625** | **1.752** | **1.292** |
| 114. | P02679-2 | Isoform Gamma-A of Fibrinogen gamma chain OS=Homo sapiens GN=FGG - [FIBG_HUMAN] | 92.32 | **0.564** | **0.621** | **1.321** |
| 115. | F5H8G2 | Lamin tail domain-containing protein 1 (Fragment) OS=Homo sapiens GN=LMNTD1 PE=4 SV=7 - [F5H8G2_HUMAN] | 3.38 |  |  |  |
| 116. | F8WCZ7 | Leucine-rich repeat-containing protein 75B OS=Homo sapiens GN=LRRC75B PE=4 SV=1 - [F8WCZ7_HUMAN] | 2.20 |  |  |  |
| 117. | E9PKI9 | Leukocyte cell-derived chemotaxin 1 (Fragment) OS=Homo sapiens GN=LECT1 PE=1 SV=1 - [E9PKI9_HUMAN] | 2.65 |  |  |  |
| 118. | P51884 | Lumican OS=Homo sapiens GN=LUM PE=1 SV=2 - [LUM_HUMAN] | 11.56 | **0.523** | 0.756 | **1.678** |
| 119. | A0A0A0MSR6 | Lysine-specific demethylase 4C (Fragment) OS=Homo sapiens GN=KDM4C PE=1 SV=1 - [A0A0A0MSR6_HUMAN] | 3.43 | 0.752 |  |  |
| 120. | A0A109PSY4 | MS-A1 light chain variable region (Fragment) OS=Homo sapiens PE=2 SV=1 - [A0A109PSY4_HUMAN] | 44.37 |  | 0.908 | **2.103** |
| 121. | A0A125U0U7 | MS-C1 heavy chain variable region (Fragment) OS=Homo sapiens PE=2 SV=1 - [A0A125U0U7_HUMAN] | 20.91 | 1.063 | **1.259** | 1.088 |
| 122. | A0A0X9TD47 | MS-D1 light chain variable region (Fragment) OS=Homo sapiens PE=2 SV=1 - [A0A0X9TD47_HUMAN] | 43.69 | 0.881 | 0.849 | 1.131 |
| 123. | A0A0X9UWK7 | MS-D4 heavy chain variable region (Fragment) OS=Homo sapiens PE=2 SV=1 - [A0A0X9UWK7_HUMAN] | 24.38 | 1.070 | **1.535** | 0.852 |
| 124. | A0A125U0V1 | MS-F1 heavy chain variable region (Fragment) OS=Homo sapiens PE=2 SV=1 - [A0A125U0V1_HUMAN] | 2.82 | 1.024 | **1.784** | **1.879** |
| 125. | A0A0X9V9B3 | MS-F1 light chain variable region (Fragment) OS=Homo sapiens PE=2 SV=1 - [A0A0X9V9B3_HUMAN] | 13.51 |  | **1.516** | **1.265** |
| 126. | P13533 | Myosin-6 OS=Homo sapiens GN=MYH6 PE=1 SV=5 - [MYH6_HUMAN] | 6.40 | **1.413** | 0.981 | **1.229** |
| 127. | Q9UL88 | Myosin-reactive immunoglobulin heavy chain variable region (Fragment) OS=Homo sapiens PE=2 SV=1 - [Q9UL88_HUMAN] | 27.84 | **1.222** | **0.668** | **0.707** |
| 128. | Q9UL89 | Myosin-reactive immunoglobulin heavy chain variable region (Fragment) OS=Homo sapiens PE=2 SV=1 - [Q9UL89_HUMAN] | 17.25 | **0.616** | **1.251** | **1.793** |
| 129. | Q9UL90 | Myosin-reactive immunoglobulin heavy chain variable region (Fragment) OS=Homo sapiens PE=2 SV=1 - [Q9UL90_HUMAN] | 22.86 | 1.039 | **0.372** | **1.654** |
| 130. | Q9UL83 | Myosin-reactive immunoglobulin light chain variable region (Fragment) OS=Homo sapiens PE=2 SV=1 - [Q9UL83_HUMAN] | 25.81 | 1.161 | **1.508** | **2.284** |
| 131. | Q96PD5 | N-acetylmuramoyl-L-alanine amidase OS=Homo sapiens GN=PGLYRP2 PE=1 SV=1 - [PGRP2_HUMAN] | 43.02 | 1.137 | **0.710** | 1.092 |
| 132. | H0Y6T6 | Nurim (Fragment) OS=Homo sapiens GN=NRM PE=1 SV=1 - [H0Y6T6_HUMAN] | 2.19 |  |  |  |
| 133. | U3KQU8 | Oxidoreductase NAD-binding domain-containing protein 1 OS=Homo sapiens GN=OXNAD1 PE=1 SV=1 - [U3KQU8_HUMAN] | 2.94 | **1.632** |  |  |
| 134. | P80108 | Phosphatidylinositol-glycan-specific phospholipase D OS=Homo sapiens GN=GPLD1 PE=1 SV=3 - [PHLD_HUMAN] | 3.52 | **1.625** | 1.078 | 0.913 |
| 135. | P36955 | Pigment epithelium-derived factor OS=Homo sapiens GN=SERPINF1 PE=1 SV=4 - [PEDF_HUMAN] | 58.03 | **1.695** | **0.402** | 0.899 |
| 136. | B2R7F8 | Plasminogen OS=Homo sapiens PE=2 SV=1 - [B2R7F8_HUMAN] | 99.93 |  | 1.070 | 1.131 |
| 137. | A0A0C4DG60 | Probable bifunctional methylenetetrahydrofolate dehydrogenase/cyclohydrolase 2 OS=Homo sapiens GN=MTHFD2L PE=1 SV=1 - [A0A0C4DG60_HUMAN] | 2.72 | **1.533** | **0.238** | **0.580** |
| 138. | P02760 | Protein AMBP OS=Homo sapiens GN=AMBP PE=1 SV=1 - [AMBP_HUMAN] | 92.90 | **1.611** | 0.881 | **1.360** |
| 139. | A0A087WU91 | Protein IGHV3-49 OS=Homo sapiens GN=IGHV3-49 PE=1 SV=1 - [A0A087WU91_HUMAN] | 7.76 |  | **1.235** | **1.420** |
| 140. | A0A0U1RRF4 | Protein IGKV1-8 OS=Homo sapiens GN=IGKV1-8 PE=4 SV=1 - [A0A0U1RRF4_HUMAN] | 352.62 | **1.633** |  |  |
| 141. | A0A0B4J2D9 | Protein IGKV1D-13 (Fragment) OS=Homo sapiens GN=IGKV1D-13 PE=1 SV=1 - [A0A0B4J2D9_HUMAN] | 9.38 | **0.465** |  |  |
| 142. | A0A0A0MTQ6 | Protein IGKV2D-28 OS=Homo sapiens GN=IGKV2D-28 PE=1 SV=1 - [A0A0A0MTQ6_HUMAN] | 58.71 | **2.196** | **1.235** | **1.259** |
| 143. | A0A087X0P6 | Protein IGKV2D-29 OS=Homo sapiens GN=IGKV2D-29 PE=1 SV=1 - [A0A087X0P6_HUMAN] | 36.44 |  |  |  |
| 144. | A0A075B6H7 | Protein IGKV3-7 (Fragment) OS=Homo sapiens GN=IGKV3-7 PE=4 SV=1 - [A0A075B6H7_HUMAN] | 17.64 | **1.633** | **0.465** | **0.658** |
| 145. | A0A0A0MT36 | Protein IGKV6D-21 (Fragment) OS=Homo sapiens GN=IGKV6D-21 PE=4 SV=1 - [A0A0A0MT36_HUMAN] | 2.81 | **0.633** | **2.196** | **2.980** |
| 146. | E9PIT3 | Prothrombin OS=Homo sapiens GN=F2 PE=1 SV=1 - [E9PIT3_HUMAN] | 150.94 | **0.695** | **0.694** | 0.898 |
| 147. | Q6MZQ6 | Putative uncharacterized protein DKFZp686G11190 OS=Homo sapiens GN=DKFZp686G11190 PE=2 SV=1 - [Q6MZQ6_HUMAN] | 1198.85 |  | 0.760 | 0.760 |
| 148. | Q6N093 | Putative uncharacterized protein DKFZp686I04196 (Fragment) OS=Homo sapiens GN=DKFZp686I04196 PE=2 SV=1 - [Q6N093_HUMAN] | 666.49 |  | **0.603** | 0.784 |
| 149. | Q6MZV6 | Putative uncharacterized protein DKFZp686L19235 OS=Homo sapiens GN=DKFZp686L19235 PE=2 SV=1 - [Q6MZV6_HUMAN] | 153.63 |  |  |  |
| 150. | Q68D25 | Putative uncharacterized protein DKFZp686N08236 (Fragment) OS=Homo sapiens GN=DKFZp686N08236 PE=2 SV=1 - [Q68D25_HUMAN] | 2.53 | **2.039** | **3.131** | **2.487** |
| 151. | Q75MU0 | Putative uncharacterized protein LIMK1 (Fragment) OS=Homo sapiens GN=LIMK1 PE=4 SV=1 - [Q75MU0_HUMAN] | 1.95 | **1.695** | **1.236** | **1.363** |
| 152. | A0A0C4DGV7 | Retinol-binding protein 4 OS=Homo sapiens GN=RBP4 PE=1 SV=1 - [A0A0C4DGV7_HUMAN] | 73.16 | **1.235** | 1.154 | **1.740** |
| 153. | A2J1N5 | Rheumatoid factor RF-ET6 (Fragment) OS=Homo sapiens PE=2 SV=1 - [A2J1N5_HUMAN] | 53.95 |  | 1.081 | **1.203** |
| 154. | A2J1M8 | Rheumatoid factor RF-IP12 (Fragment) OS=Homo sapiens PE=2 SV=1 - [A2J1M8_HUMAN] | 12.94 | **1.363** | **2.033** | **1.936** |
| 155. | F8W8I8 | Septin-8 OS=Homo sapiens GN=SEPT8 PE=1 SV=1 - [F8W8I8_HUMAN] | 2.43 |  |  |  |
| 156. | Q5UGI6 | Serine/cysteine proteinase inhibitor clade G member 1 splice variant 2 (Fragment) OS=Homo sapiens GN=SERPING1 PE=2 SV=1 - [Q5UGI6_HUMAN] | 83.24 |  | 1.065 | **1.398** |
| 157. | M0R3G6 | Serine/threonine-protein kinase PAK 4 (Fragment) OS=Homo sapiens GN=PAK4 PE=1 SV=1 - [M0R3G6_HUMAN] | 2.79 |  |  |  |
| 158. | P02787 | Serotransferrin OS=Homo sapiens GN=TF PE=1 SV=3 - [TRFE_HUMAN] | 1515.67 | **0.591** | **0.656** | **0.602** |
| 159. | A0A0K0Q2Z1 | Serpin peptidase inhibitor clade C member 1 OS=Homo sapiens GN=SERPINC1 PE=2 SV=1 - [A0A0K0Q2Z1_HUMAN] | 192.61 | **1.235** | 1.087 | **1.399** |
| 160. | P02743 | Serum amyloid P-component OS=Homo sapiens GN=APCS PE=1 SV=2 - [SAMP_HUMAN] | 21.74 | **1.320** | **1.280** | 1.066 |
| 161. | P27169 | Serum paraoxonase/arylesterase 1 OS=Homo sapiens GN=PON1 PE=1 SV=3 - [PON1_HUMAN] | 30.98 | **1.451** | **1.695** | **2.208** |
| 162. | Q65ZC9 | Single-chain Fv (Fragment) OS=Homo sapiens GN=scFv PE=2 SV=1 - [Q65ZC9_HUMAN] | 32.23 |  |  |  |
| 163. | H0YCA5 | Spermatogenesis-associated protein 5-like protein 1 (Fragment) OS=Homo sapiens GN=SPATA5L1 PE=1 SV=1 - [H0YCA5_HUMAN] | 2.85 |  |  |  |
| 164. | E9PHK0 | Tetranectin OS=Homo sapiens GN=CLEC3B PE=1 SV=1 - [E9PHK0_HUMAN] | 2.69 | **1.695** | **1.544** | **1.479** |
| 165. | J3KNC0 | Transcription initiation factor IIA subunit 1 OS=Homo sapiens GN=GTF2A1 PE=1 SV=1 - [J3KNC0_HUMAN] | 2.06 |  |  |  |
| 166. | P02766 | Transthyretin OS=Homo sapiens GN=TTR PE=1 SV=1 - [TTHY_HUMAN] | 91.64 | **1.343** | **1.755** | **1.355** |
| 167. | Q68CN4 | Uncharacterized protein OS=Homo sapiens GN=DKFZp686E23209 PE=1 SV=2 - [Q68CN4_HUMAN] | 499.18 |  |  | 1.168 |
| 168. | Q6N095 | Uncharacterized protein OS=Homo sapiens GN=DKFZp686K03196 PE=1 SV=1 - [Q6N095_HUMAN] | 965.28 |  | **0.546** | 1.057 |
| 169. | Q569I7 | Uncharacterized protein OS=Homo sapiens PE=2 SV=1 - [Q569I7_HUMAN] | 356.87 | **1.965** | **1.385** | 1.143 |
| 170. | Q6GMV8 | Uncharacterized protein OS=Homo sapiens PE=2 SV=1 - [Q6GMV8_HUMAN] | 257.02 |  |  |  |
| 171. | E7ETN3 | Uncharacterized protein OS=Homo sapiens PE=3 SV=1 - [E7ETN3_HUMAN] | 13.15 |  |  |  |
| 172. | Q5NV88 | V1-22 protein (Fragment) OS=Homo sapiens GN=V1-22 PE=1 SV=1 - [Q5NV88_HUMAN] | 2.09 |  | 1.179 | **2.054** |
| 173. | A2MYD4 | V2-7 protein (Fragment) OS=Homo sapiens GN=V2-7 PE=4 SV=1 - [A2MYD4_HUMAN] | 11.39 |  | **1.564** | **1.834** |
| 174. | P02774 | Vitamin D-binding protein OS=Homo sapiens GN=GC PE=1 SV=1 - [VTDB_HUMAN] | 159.83 | 1.162 | 0.810 | **1.377** |
| 175. | P04004 | Vitronectin OS=Homo sapiens GN=VTN PE=1 SV=1 - [VTNC_HUMAN] | 138.61 | **1.405** | 1.148 | **1.406** |
| 176. | P25311 | Zinc-alpha-2-glycoprotein OS=Homo sapiens GN=AZGP1 PE=1 SV=2 - [ZA2G_HUMAN] | 74.32 | **1.519** | **1.307** | **1.405** |

**Additional Table 3 Summary of audiometry statistical data analysis repot**

|  | Dunnett's multiple comparisons test | Predicted (LS) mean diff. | 95.00% CI of diff. | Adjusted P Value | Predicted (LS) mean diff. | 95.00% CI of diff. | Adjusted P Value |
| --- | --- | --- | --- | --- | --- | --- | --- |
| 0.125 kHz | Normal vs. Mild | -4.62 | -6.36 to -2.88 | <0.0001 | -3.33 | -6.79 to 0.133 | 0.0633 |
|  | Normal vs. Moderate | -1.68 | -3.72 to 0.374 | 0.1419 | -3.16 | -8.42 to 2.10 | 0.3777 |
|  | Normal vs. Severe | -1.98 | -4.25 to 0.283 | 0.1044 | -3.13 | -8.13 to 1.88 | 0.3433 |
| 0.250 kHz | Normal vs. Mild | -4.87 | -6.61 to -3.13 | <0.0001 | -3.32 | -6.78 to 0.141 | 0.0642 |
|  | Normal vs. Moderate | -1.6 | -3.65 to 0.451 | 0.1719 | -3.61 | -8.87 to 1.65 | 0.2668 |
|  | Normal vs. Severe | -0.91 | -3.18 to 1.36 | 0.6947 | -2.77 | -7.78 to 2.23 | 0.4472 |
| 0.500 kHz | Normal vs. Mild | -3.02 | -4.76 to -1.28 | 0.0001 | -1.04 | -4.51 to 2.42 | 0.8421 |
|  | Normal vs. Moderate | -2.3 | -4.35 to -0.248 | 0.0225 | -1.49 | -6.75 to 3.77 | 0.8646 |
|  | Normal vs. Severe | -0.672 | -2.94 to 1.60 | 0.849 | -1.94 | -6.95 to 3.06 | 0.7158 |
| 0.750 kHz | Normal vs. Mild | -2.91 | -4.65 to -1.17 | 0.0002 | -2.04 | -5.50 to 1.42 | 0.3944 |
|  | Normal vs. Moderate | -1.83 | -3.88 to 0.217 | 0.0938 | -1.88 | -7.14 to 3.38 | 0.7625 |
|  | Normal vs. Severe | -1.46 | -3.73 to 0.803 | 0.3154 | -3.13 | -8.14 to 1.87 | 0.3414 |
| 1 kHz | Normal vs. Mild | -2.26 | -4.00 to -0.520 | 0.006 | -1.64 | -5.10 to 1.82 | 0.5763 |
|  | Normal vs. Moderate | -1.22 | -3.27 to 0.833 | 0.387 | -2.21 | -7.47 to 3.05 | 0.6654 |
|  | Normal vs. Severe | -0.834 | -3.10 to 1.43 | 0.7478 | -3.99 | -9.00 to 1.01 | 0.157 |
| 1.5 kHz | Normal vs. Mild | -2.15 | -3.89 to -0.412 | 0.0097 | -2.24 | -5.71 to 1.22 | 0.3133 |
|  | Normal vs. Moderate | -2.25 | -4.30 to -0.201 | 0.0263 | -2.06 | -7.32 to 3.20 | 0.7093 |
|  | Normal vs. Severe | -0.941 | -3.21 to 1.33 | 0.6728 | -5.04 | -10.0 to -0.0332 | 0.048 |
| 2 kHz | Normal vs. Mild | -2.25 | -3.99 to -0.514 | 0.0062 | -2.09 | -5.55 to 1.38 | 0.374 |
|  | Normal vs. Moderate | -2.61 | -4.66 to -0.562 | 0.0073 | -4.98 | -10.2 to 0.283 | 0.0694 |
|  | Normal vs. Severe | -2.25 | -4.52 to 0.0138 | 0.0519 | -6.52 | -11.5 to -1.51 | 0.0059 |
| 3 kHz | Normal vs. Mild | -5.57 | -7.31 to -3.83 | <0.0001 | -3.45 | -6.91 to 0.0126 | 0.0511 |
|  | Normal vs. Moderate | -6.62 | -8.67 to -4.57 | <0.0001 | -11.4 | -16.6 to -6.10 | <0.0001 |
|  | Normal vs. Severe | -10.3 | -12.6 to -8.04 | <0.0001 | -13.5 | -18.5 to -8.47 | <0.0001 |
| 4 kHz | Normal vs. Mild | -8.84 | -10.6 to -7.10 | <0.0001 | -5.13 | -8.60 to -1.67 | 0.0013 |
|  | Normal vs. Moderate | -12.7 | -14.8 to -10.7 | <0.0001 | -18.4 | -23.6 to -13.1 | <0.0001 |
|  | Normal vs. Severe | -26.7 | -28.9 to -24.4 | <0.0001 | -23 | -28.0 to -18.0 | <0.0001 |
| 6 kHz | Normal vs. Mild | -12.7 | -14.4 to -10.9 | <0.0001 | -9.75 | -13.2 to -6.29 | <0.0001 |
|  | Normal vs. Moderate | -19.9 | -22.0 to -17.9 | <0.0001 | -23.3 | -28.6 to -18.0 | <0.0001 |
|  | Normal vs. Severe | -41.4 | -43.7 to -39.1 | <0.0001 | -40.6 | -45.6 to -35.6 | <0.0001 |
| 8 kHz | Normal vs. Mild | -15.9 | -17.7 to -14.2 | <0.0001 | -6.35 | -9.81 to -2.88 | <0.0001 |
|  | Normal vs. Moderate | -26.4 | -28.4 to -24.3 | <0.0001 | -20.8 | -26.1 to -15.6 | <0.0001 |
|  | Normal vs. Severe | -39.5 | -41.7 to -37.2 | <0.0001 | -44.7 | -49.7 to -39.7 | <0.0001 |

**Additional Table 4.** Individual Wave I-V Latency Statistical Analysis. The table presents the mean latencies, standard deviations, and results from statistical tests comparing latencies between normal and impaired hearing groups (mild, moderate, severe). For each wave, latency values are displayed along with their corresponding 95% confidence intervals (CI) and p-values, reflecting differences in wave latencies at each severity level.

|  | **Peak I** |  | | | **Peak III** | | | **Peak V** | | |
| --- | --- | --- | --- | --- | --- | --- | --- | --- | --- | --- |
| **Decibel** | **Dunnett's multiple comparisons test** | **Predicted (LS) mean diff.** | **95.00% CI of diff.** | **Adjusted P Value** | **Predicted (LS) mean diff.** | **95.00% CI of diff.** | **Adjusted P Value** | **Predicted (LS) mean diff.** | **95.00% CI of diff.** | **Adjusted P Value** |
| 20 | No vs. Mild | -0.0197 | -0.0644 to 0.0251 | 0.62 | -0.373 | -0.427 to -0.319 | <0.001 | -0.462 | -0.537 to -0.387 | <0.001 |
|  | Normal vs. Moderate | -0.0647 | -0.113 to -0.0165 | 0.004 | -0.852 | -0.910 to -0.793 | <0.001 | -0.906 | -0.987 to -0.825 | <0.001 |
|  | Normal vs. Severe | -0.119 | -0.180 to -0.0578 | <0.001 | -1.05 | -1.13 to -0.978 | <0.001 | -1.02 | -1.12 to -0.912 | <0.001 |
| 25 | Normal vs. Mild | -0.0197 | -0.0644 to 0.0251 | 0.62 | -0.32 | -0.374 to -0.266 | <0.001 | -0.409 | -0.484 to -0.334 | <0.001 |
|  | Normal vs. Moderate | -0.0647 | -0.113 to -0.0165 | 0.004 | -0.767 | -0.825 to -0.708 | <0.001 | -0.821 | -0.902 to -0.740 | <0.001 |
|  | Normal vs. Severe | -0.119 | -0.180 to -0.0578 | <0.001 | -0.867 | -0.941 to -0.793 | <0.001 | -0.83 | -0.933 to -0.727 | <0.001 |
| 30 | Normal vs. Mild | -0.0197 | -0.0644 to 0.0251 | 0.62 | -0.495 | -0.549 to -0.441 | <0.001 | -0.484 | -0.560 to -0.409 | <0.001 |
|  | Normal vs. Moderate | -0.0647 | -0.113 to -0.0165 | 0.004 | -0.886 | -0.944 to -0.828 | <0.001 | -0.84 | -0.921 to -0.759 | <0.001 |
|  | Normal vs. Severe | -0.119 | -0.180 to -0.0578 | <0.001 | -0.916 | -0.990 to -0.842 | <0.001 | -0.779 | -0.882 to -0.676 | <0.001 |
| 35 | Normal vs. Mild | -0.0197 | -0.0644 to 0.0251 | 0.62 | -0.447 | -0.501 to -0.393 | <0.001 | -0.386 | -0.461 to -0.311 | <0.001 |
|  | Normal vs. Moderate | -0.0647 | -0.113 to -0.0165 | 0.004 | -0.871 | -0.929 to -0.813 | <0.001 | -0.775 | -0.856 to -0.694 | <0.001 |
|  | Normal vs. Severe | -0.119 | -0.180 to -0.0578 | <0.001 | -0.931 | -1.00 to -0.857 | <0.001 | -0.744 | -0.847 to -0.641 | <0.001 |
| 40 | Normal vs. Mild | -0.0197 | -0.0644 to 0.0251 | 0.62 | -0.55 | -0.604 to -0.496 | <0.001 | -0.509 | -0.585 to -0.434 | <0.001 |
|  | Normal vs. Moderate | -0.0647 | -0.113 to -0.0165 | 0.004 | -0.732 | -0.791 to -0.674 | <0.001 | -0.657 | -0.738 to -0.576 | <0.001 |
|  | Normal vs. Severe | -0.119 | -0.180 to -0.0578 | <0.001 | -0.88 | -0.954 to -0.806 | <0.001 | -0.713 | -0.816 to -0.610 | <0.001 |
| 45 | Normal vs. Mild | -0.0197 | -0.0644 to 0.0251 | 0.62 | -0.443 | -0.497 to -0.389 | <0.001 | -0.432 | -0.507 to -0.356 | <0.001 |
|  | Normal vs. Moderate | -0.0647 | -0.113 to -0.0165 | 0.004 | -0.706 | -0.764 to -0.647 | <0.001 | -0.66 | -0.741 to -0.579 | <0.001 |
|  | Normal vs. Severe | -0.119 | -0.180 to -0.0578 | <0.001 | -0.766 | -0.840 to -0.692 | <0.001 | -0.629 | -0.732 to -0.526 | <0.001 |
| 50 | Normal vs. Mild | -0.0197 | -0.0644 to 0.0251 | 0.62 | -0.57 | -0.625 to -0.516 | <0.001 | -0.459 | -0.535 to -0.384 | <0.001 |
|  | Normal vs. Moderate | -0.0647 | -0.113 to -0.0165 | 0.004 | -0.794 | -0.852 to -0.736 | <0.001 | -0.649 | -0.730 to -0.568 | <0.001 |
|  | Normal vs. Severe | -0.119 | -0.180 to -0.0578 | <0.001 | -0.854 | -0.928 to -0.780 | <0.001 | -0.618 | -0.721 to -0.515 | <0.001 |
| 55 | Normal vs. Mild | -0.0197 | -0.0644 to 0.0251 | 0.62 | -0.326 | -0.380 to -0.272 | <0.001 | -0.215 | -0.290 to -0.140 | <0.001 |
|  | Normal vs. Moderate | -0.0647 | -0.113 to -0.0165 | 0.004 | -0.727 | -0.785 to -0.669 | <0.001 | -0.581 | -0.662 to -0.500 | <0.001 |
|  | Normal vs. Severe | -0.119 | -0.180 to -0.0578 | <0.001 | -0.797 | -0.871 to -0.723 | <0.001 | -0.561 | -0.663 to -0.458 | <0.001 |
| 60 | Normal vs. Mild | -0.0197 | -0.0644 to 0.0251 | 0.62 | -0.294 | -0.348 to -0.240 | <0.001 | -0.0833 | -0.159 to -0.00800 | 0.03 |
|  | Normal vs. Moderate | -0.0647 | -0.113 to -0.0165 | 0.004 | -0.614 | -0.673 to -0.556 | <0.001 | -0.369 | -0.450 to -0.288 | <0.001 |
|  | Normal vs. Severe | -0.119 | -0.180 to -0.0578 | <0.001 | -0.814 | -0.888 to -0.740 | <0.001 | -0.478 | -0.581 to -0.375 | <0.001 |
| 65 | Normal vs. Mild | -0.0197 | -0.0644 to 0.0251 | 0.62 | -0.113 | -0.167 to -0.0592 | <0.001 | -0.0123 | -0.0876 to 0.0629 | 0.97 |
|  | Normal vs. Moderate | -0.0647 | -0.113 to -0.0165 | 0.004 | -0.444 | -0.502 to -0.385 | <0.001 | -0.308 | -0.389 to -0.227 | <0.001 |
|  | Normal vs. Severe | -0.119 | -0.180 to -0.0578 | <0.001 | -0.644 | -0.718 to -0.570 | <0.001 | -0.417 | -0.520 to -0.314 | <0.001 |
| 70 | Normal vs. Mild | -0.0197 | -0.0644 to 0.0251 | 0.62 | -0.103 | -0.157 to -0.0491 | <0.001 | -0.0322 | -0.107 to 0.0431 | 0.64 |
|  | Normal vs. Moderate | -0.0647 | -0.113 to -0.0165 | 0.004 | -0.349 | -0.407 to -0.290 | <0.001 | -0.243 | -0.324 to -0.162 | <0.001 |
|  | Normal vs. Severe | -0.119 | -0.180 to -0.0578 | <0.001 | -0.449 | -0.523 to -0.375 | <0.001 | -0.252 | -0.355 to -0.149 | <0.001 |

**Additional Table 5.** Individual Wave I-V Amplitude Statistical Analysis. The table presents the mean amplitude, standard deviations, and results from statistical tests comparing latencies between normal and impaired hearing groups (mild, moderate, severe). For each wave, amplitude values are displayed along with their corresponding 95% confidence intervals (CI) and p-values, reflecting differences in wave amplitude at each severity level

|  | **Amplitude I-Ia** |  | | | **Amplitude III-IIIa** | | | **Amplitude V-Va** | | |
| --- | --- | --- | --- | --- | --- | --- | --- | --- | --- | --- |
| **Decibel** | **Dunnett's multiple comparisons test** | **Predicted (LS) mean diff.** | **95.00% CI of diff.** | **Adjusted P Value** | **Predicted (LS) mean diff.** | **95.00% CI of diff.** | **Adjusted P Value** | **Predicted (LS) mean diff.** | **95.00% CI of diff.** | **Adjusted P Value** |
| 20 | No vs. Mild | 0.921 | 0.732 to 1.11 | <0.001 | 0.184 | 0.108 to 0.261 | <0.001 | 0.153 | 0.0656 to 0.239 | <0.001 |
|  | Normal vs. Moderate | 1.28 | 1.07 to 1.48 | <0.001 | 0.258 | 0.175 to 0.340 | <0.001 | 0.41 | 0.316 to 0.504 | <0.001 |
|  | Normal vs. Severe | 1.28 | 1.02 to 1.54 | <0.001 | 0.258 | 0.153 to 0.363 | <0.001 | 0.41 | 0.291 to 0.529 | <0.001 |
| 25 | Normal vs. Mild | 0.987 | 0.798 to 1.18 | <0.001 | 0.197 | 0.120 to 0.273 | <0.001 | 0.159 | 0.0721 to 0.246 | <0.001 |
|  | Normal vs. Moderate | 1.62 | 1.41 to 1.82 | <0.001 | 0.326 | 0.244 to 0.409 | <0.001 | 0.428 | 0.334 to 0.521 | <0.001 |
|  | Normal vs. Severe | 1.62 | 1.36 to 1.87 | <0.001 | 0.326 | 0.221 to 0.431 | <0.001 | 0.428 | 0.309 to 0.547 | <0.001 |
| 30 | Normal vs. Mild | 1.09 | 0.903 to 1.28 | <0.001 | 0.217 | 0.140 to 0.293 | <0.001 | 0.135 | 0.0481 to 0.222 | <0.001 |
|  | Normal vs. Moderate | 1.66 | 1.46 to 1.86 | <0.001 | 0.329 | 0.246 to 0.411 | <0.001 | 0.179 | 0.0859 to 0.273 | <0.001 |
|  | Normal vs. Severe | 2.02 | 1.76 to 2.28 | <0.001 | 0.407 | 0.302 to 0.512 | <0.001 | 0.458 | 0.339 to 0.576 | <0.001 |
| 35 | Normal vs. Mild | 1.18 | 0.991 to 1.37 | <0.001 | 0.233 | 0.157 to 0.310 | <0.001 | 0.141 | 0.0539 to 0.228 | <0.001 |
|  | Normal vs. Moderate | 1.72 | 1.52 to 1.92 | <0.001 | 0.336 | 0.254 to 0.419 | <0.001 | 0.132 | 0.0380 to 0.225 | 0.003 |
|  | Normal vs. Severe | 1.99 | 1.74 to 2.25 | <0.001 | 0.374 | 0.269 to 0.479 | <0.001 | 0.187 | 0.0683 to 0.306 | <0.001 |
| 40 | Normal vs. Mild | 0.678 | 0.489 to 0.866 | <0.001 | 0.129 | 0.0518 to 0.205 | <0.001 | 0.46 | 0.373 to 0.547 | <0.001 |
|  | Normal vs. Moderate | 1.73 | 1.53 to 1.93 | <0.001 | 0.333 | 0.250 to 0.415 | <0.001 | 0.465 | 0.371 to 0.559 | <0.001 |
|  | Normal vs. Severe | 2.03 | 1.77 to 2.28 | <0.001 | 0.359 | 0.254 to 0.464 | <0.001 | 0.575 | 0.456 to 0.694 | <0.001 |
| 45 | Normal vs. Mild | 0.592 | 0.403 to 0.780 | <0.001 | 0.109 | 0.0323 to 0.186 | 0.002 | 0.408 | 0.321 to 0.495 | <0.001 |
|  | Normal vs. Moderate | 1.92 | 1.72 to 2.13 | <0.001 | 0.367 | 0.285 to 0.450 | <0.001 | 0.414 | 0.320 to 0.508 | <0.001 |
|  | Normal vs. Severe | 2.17 | 1.91 to 2.43 | <0.001 | 0.364 | 0.259 to 0.469 | <0.001 | 0.488 | 0.369 to 0.607 | <0.001 |
| 50 | Normal vs. Mild | 1.51 | 1.32 to 1.70 | <0.001 | 0.294 | 0.217 to 0.370 | <0.001 | 0.573 | 0.486 to 0.659 | <0.001 |
|  | Normal vs. Moderate | 2.23 | 2.03 to 2.43 | <0.001 | 0.415 | 0.333 to 0.498 | <0.001 | 0.578 | 0.485 to 0.672 | <0.001 |
|  | Normal vs. Severe | 3.04 | 2.78 to 3.30 | <0.001 | 0.52 | 0.415 to 0.625 | <0.001 | 0.565 | 0.446 to 0.684 | <0.001 |
| 55 | Normal vs. Mild | 1.49 | 1.30 to 1.68 | <0.001 | 0.286 | 0.210 to 0.363 | <0.001 | 0.515 | 0.428 to 0.602 | <0.001 |
|  | Normal vs. Moderate | 2.4 | 2.20 to 2.60 | <0.001 | 0.44 | 0.358 to 0.523 | <0.001 | 0.522 | 0.429 to 0.616 | <0.001 |
|  | Normal vs. Severe | 2.92 | 2.67 to 3.18 | <0.001 | 0.433 | 0.328 to 0.538 | <0.001 | 0.46 | 0.341 to 0.579 | <0.001 |
| 60 | Normal vs. Mild | 1.63 | 1.44 to 1.82 | <0.001 | 0.314 | 0.237 to 0.391 | <0.001 | 0.473 | 0.386 to 0.559 | <0.001 |
|  | Normal vs. Moderate | 2.13 | 1.92 to 2.33 | <0.001 | 0.373 | 0.291 to 0.456 | <0.001 | 0.481 | 0.388 to 0.575 | <0.001 |
|  | Normal vs. Severe | 2.6 | 2.34 to 2.85 | <0.001 | 0.309 | 0.204 to 0.414 | <0.001 | 0.49 | 0.371 to 0.609 | <0.001 |
| 65 | Normal vs. Mild | 1.53 | 1.34 to 1.72 | <0.001 | 0.29 | 0.213 to 0.367 | <0.001 | 0.365 | 0.278 to 0.452 | <0.001 |
|  | Normal vs. Moderate | 2.16 | 1.96 to 2.37 | <0.001 | 0.366 | 0.283 to 0.448 | <0.001 | 0.376 | 0.282 to 0.470 | <0.001 |
|  | Normal vs. Severe | 2.75 | 2.50 to 3.01 | <0.001 | 0.284 | 0.180 to 0.389 | <0.001 | 0.387 | 0.268 to 0.506 | <0.001 |
| 70 | Normal vs. Mild | 0.741 | 0.552 to 0.930 | <0.001 | 0.126 | 0.0490 to 0.202 | <0.001 | 0.568 | 0.481 to 0.654 | <0.001 |
|  | Normal vs. Moderate | 1.4 | 1.20 to 1.60 | <0.001 | 0.193 | 0.111 to 0.276 | <0.001 | 0.58 | 0.486 to 0.673 | <0.001 |
|  | Normal vs. Severe | 2.8 | 2.54 to 3.06 | <0.001 | 0.274 | 0.170 to 0.379 | <0.001 | 0.591 | 0.473 to 0.710 | <0.001 |
